# Mechanical stability and unfolding pathways of parallel tetrameric G-quadruplexes probed by pulling simulations

**DOI:** 10.1101/2024.01.26.577375

**Authors:** Zhengyue Zhang, Vojtěch Mlýnský, Miroslav Krepl, Jiří Šponer, Petr Stadlbauer

## Abstract

Guanine-quadruplex (GQ) is a noncanonical nucleic acid structure formed by guanine-rich DNA and RNA sequences. Folding of GQs is a complex process, many aspects of which remain elusive, despite being important for understanding of structure formation and biological functions of GQs. Pulling experiments are a common tool for acquiring insights into the folding landscape of GQs. Herein, we applied a computational pulling strategy – Steered Molecular Dynamics (SMD) simulations – combined with standard Molecular Dynamics (MD) simulations to explore the unfolding landscapes of tetrameric parallel GQs. We identified anisotropic properties of elastic conformational changes, unfolding transitions, and GQ mechanical stabilities. The vertical component of pulling force (perpendicular to the average G-quartet plane) plays a significant role in disrupting GQ structures and weakening their mechanical stabilities. We demonstrated that the magnitude of vertical force component depends on the pulling anchor positions and the number of G-quartets. Typical unfolding transitions for tetrameric parallel GQs involve base unzipping, opening of the G-stem, strand slippage and rotation to cross-like structures. The unzipping was detected as the first and dominant unfolding event and it usually started at the 3’-end. Furthermore, results from both SMD and standard MD simulations indicate that partial spiral conformations serve as a transient ensemble during the (un)folding of GQs.

## Introduction

DNA Guanine-quadruplex (GQ) is a representative of noncanonical DNA structures, formed by guanine-rich sequences. It contains at least two stacked planar G-quartets and is stabilized by coordinated cations (e.g., Na^+^, K^+^ or NH_4_^+^) in its central channel (Figure 1).^1–6^ GQs participate in vital biological functions, such as gene expression or telomeric maintenance, and thus malfunctioning GQ regulation can lead to various diseases including cancers.^7–15^ The relative structural rigidity of GQs and their continuous ion-binding central channel make them attractive for nanotechnology and biocatalysis.^16,17^

**Figure 1.**
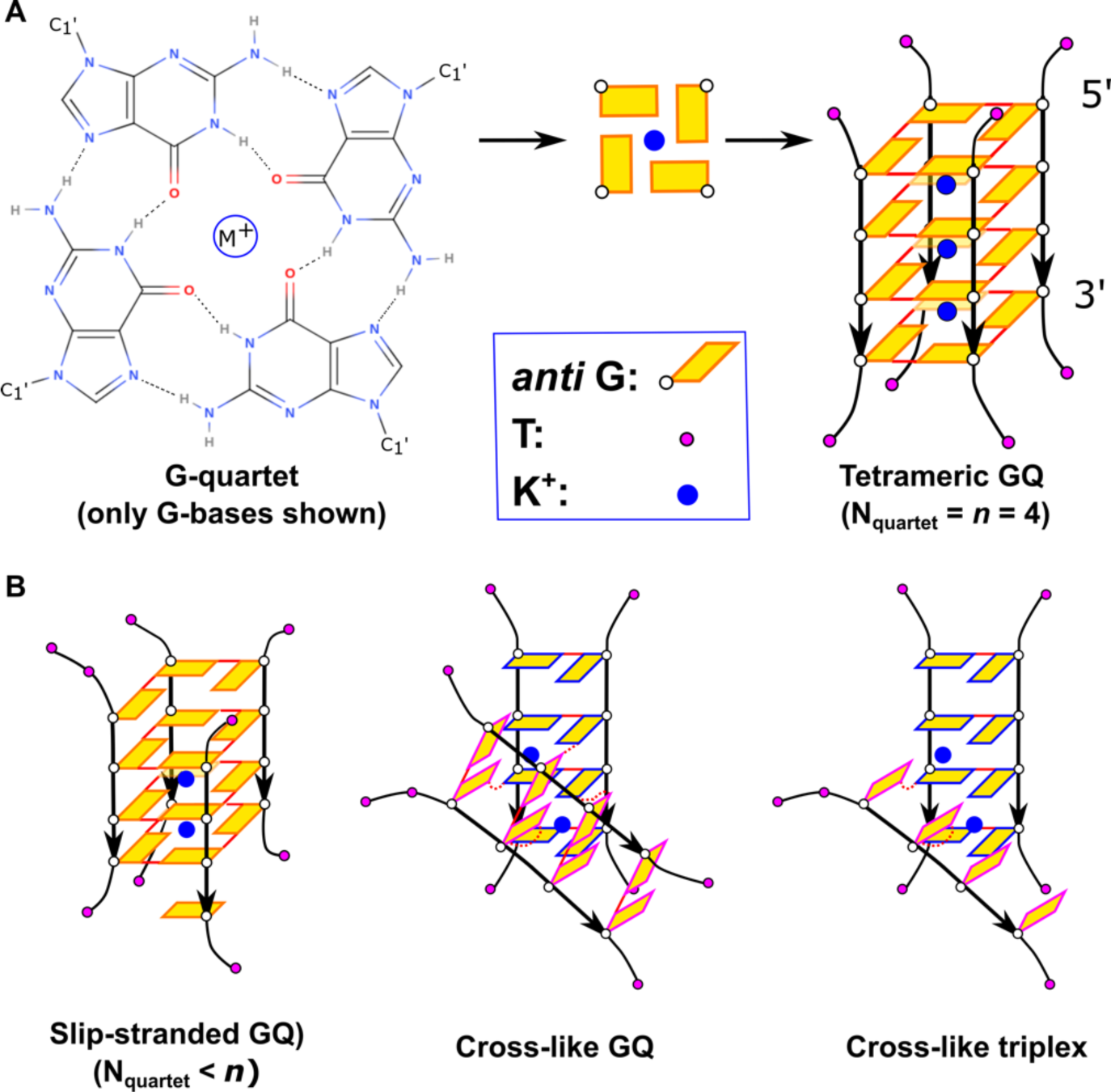
Schemes of G-quartet, tetrameric parallel GQ, and some salient folding intermediates. **(A)** G-quartet composed of four Gs with a coordinated monovalent cation M^+^ (left) stacks with three other G-quartets to form an all-*anti* four-quartet tetrameric GQ (right). In the depicted GQ, one flanking T is connected to all G-tract ends. The cations are commonly located in between the neighbouring G-quartets but can also be in the quartet plane. **(B)** Suggested intermediates in GQ folding. Slip-stranded GQ, i.e., GQ having fewer G-quartets than the G-tract length, can exist on its own, or it may vertically stack and interlock with other slip-stranded GQs to form multimeric species (notice that more than one G-tract can be slipped). Cross-like GQ and triplex are representative examples of cross-like structures, which have G-tract(s) rotated with respect to the native GQ structure.

Monomolecular (intramolecular) GQs formed by a single DNA chain folded on itself can adopt various structures due to many possible combinations of G-tract directions (parallel, antiparallel), loop types, lengths and sequences, patterns of glycosidic torsional angles and the overall number of G-quartets.^1,5,18–23^ Preferred GQ topology may also be influenced by the flanking bases,^24,25^ ionic strength and types,^4,26,27^ pH,^28,29^ or molecular crowding.^28,30^ For instance, the human telomeric (hTel) GGG(TTAGGG)_3_ sequence owns extremely high diversities, and at least six topologies have been detected under diverse experimental conditions.^2,14,27^

Bimolecular and tetramolecular GQs are formed by shorter chains of a few nucleotides (Figure 1). Furthermore, GQ units can stack together and form supramolecular multimeric assemblies.^31–36^ Tetramolecular GQs have been studied on truncated segments of genomically important sequences.^37–39^ Nevertheless, their actual biological relevance is rather negligible, because the probability that four independent chains or distant parts of a long nucleic acid chain would come together is small. On the other hand, they represent fundamental systems to understand the basic physico-chemical properties of GQs and they have been widely studied.^38,40–43^ Structures based on the [d(G_n_)]_4_ sequence, with *n* usually about 4, are tetramolecular GQs which have been characterised numerous times both in the DNA and RNA variants.^40,44^ T’s (or U’s) usually flank the G-sequence on one or both ends, to prevent vertical stacking of individual GQ units and to supress formation of loosely defined higher order multimeric structures with slipped strands (interlocked GQs; Figure 1).^45–49^ Structurally, the oligomer forms a parallel stranded GQ with all guanines in the *anti* conformation, although a small population of a *syn*-quartet at the 5’-end has been detected in solution experiment;^44,50^ the *syn*-quartet can be promoted by absence of 5’-flanking nucleotides^38,51,52^ and by some ligands.^53^ Thermodynamic stability of the GQ increases with the increasing number of G-quartets, i.e. with increasing *n*.^54,55^ The dissociation life-time of [d(TG_n_T)]_4_ GQs increases by 5-6 orders of magnitude per one added G-quartet.^54^ At the same time, increasing *n* promotes formation and stability of mismatched GQs with the number of quartets smaller than *n* or assemblies containing more than four strands.^54,56–59^ RNA GQs have been found more stable than their DNA counterparts.^55,60^ Parallel tetramolecular GQs are of interest in nanotechnology for their rod-like shape and the central channel that holds cations, so it is assumed they may act as nanowires or nanopores. While K^+^ is best for the GQ stability,^48,61–65^ computational studies focused on ion movement through the channel have predicted that Na^+^ is better for conductivity.^66–69^

Due to their importance, GQs have been widely studied by molecular dynamics (MD) simulation methods.^70,71^ In this context, high resolution structures of tetramolecular GQs obtained by X-ray crystallography serve as genuine and vital benchmarks for verification and development of DNA and RNA empirical potentials (i.e., molecular mechanics force fields).^72–75^ MD simulations of tetramolecular GQs have indeed revealed several force field artifacts, for example, excessive ion-ion repulsion in the ion channel and hydrogen bond bifurcation in the G-quartets,^76–79^ pointing to inherent limitations of standard non-polarizable force fields. Theoretical calculations have also helped to understand the *syn*/*anti* preferences of the glycosidic torsion angle *χ* in GQ stems.^51,52,80^

Both experimental and theoretical approaches have been used to study the GQ folding. For intramolecular GQs, the process can be understood within the concept of kinetic partitioning, with multiple long-living states on the landscape, acting with respect to each other as off-pathway kinetic species.^71,81–86^ On the other hand, the folding (formation) kinetics of tetramolecular GQs is specifically complicated by oligonucleotide and cation concentration dependence, because the process is of a higher reaction order with respect to both. The order is thought to be between 3 and 4, and it also depends on the cation type (Na^+^, K^+^) present in the solution.^54,55,61,62,87–90^ Indirect experimental evidence has led to proposal of several folding mechanisms: (i) sequential strand association, (ii) formation of two independent G-duplexes and their subsequent fusion, and (iii) sequential strand association into a triplex, fusion of two such triplexes into a six-stranded intermediate, followed by release of two strands and GQ formation.^88,90,91^ MD simulations have not ruled out any of the scenarios, however, they suggested that the intermediates would not be “ideal” G-duplexes and triplexes with native base pairing, but rather cross-like intermediates with rotated strands (Figure 1).^92^ The simulations have also revealed a strand slippage mechanism, which allows individual strands in a GQ to slide vertically along the other strands especially when the channel is not fully occupied by cations and the GQ is slip-stranded GQ (Figure 1).^77,92^

Single molecule pulling experimental techniques offer a way to probe folding and unfolding pathways of biomolecules. Magnetic tweezer, optical tweezer and atomic force microscopy (AFM), which can be further integrated with single molecule fluorescence resonance energy transfer technique, are the most popular experimental methods that have been used multiple times to investigate folding and unfolding mechanisms of various intramolecular GQs.^11,93–102^ As a computational analogue to experimental pulling methods, steered molecular dynamics (SMD) simulation^103,104^ drags the system out from an initial configuration by applying the external force. SMD techniques have indeed been applied to investigate unfolding of the hTel GQs^105,106,107^ and the thrombin binding aptamer.^108^

The tetrameric GQs are important systems for understanding the basic structural properties and GQ folding mechanisms, but they are technically challenging for single-molecule experiments. Hence, in this work, we used explicit solvent all-atom SMD simulations for detailed investigation of the tetrameric parallel GQ having all-*anti*-G patterns without any connecting loops. Our simulations employed a slow constant-velocity pulling regime with the pulling velocity and force constant already within the reach of modern fast AFM experimental setups.^109,110^ We analysed the effects of external force in SMD simulations for different number of G-quartets and force directions. The simulations revealed effects of the force on GQ helical structure, e.g., helical twist. During unfolding, we identified various unfolding transitions, with single G unzipping playing a prominent role. Importantly, the probability of various unfolding transitions depends on the direction of the applied force. The results indicate that the mechanism of unfolding could be significantly affected by the applied pulling protocol, e.g., choice of collective variable (CV) used for the unfolding. Besides structural transitions caused by the applied force, we discuss the effect of water model and presence of force field artifacts. Last but not the least, we selected a few unfolding intermediates frequently sampled during SMD simulations and probed them by unbiased MD simulations to obtain further insights into late stages of the tetrameric GQ formation. We observed that a partially spiral-shaped structure might be involved in unfolding as well as refolding into the native GQ. Thus, the partial spiral structure could be significantly populated within transition ensembles of folding and unfolding.

## Methods

### System Preparation

We studied three tetrameric parallel-stranded GQs: (i) we took the four-quartet structure PDB ID 352D^111^ (i.e., the sequence [d(TGGGGT)]_4_) as the starting model and replaced the Na^+^ cations by K^+^; (ii) GQ with five G-quartets [d(TGGGGGT)]_4_ was prepared by manually mutating the T-quartet on the 5’-terminal of 352D to a G-quartet, adding a K^+^ into the new site in the channel and modelling a new 5’-terminal T-quartet; (iii) GQ with three G-quartets [d(TGGGT)]_4_ was derived from 352D by removal of the 5’-terminal T-quartet, mutation of the adjacent G-quartet to T’s and removal of the extra channel K^+^. In all GQ systems, one additional T was attached to the 5’-end of the first strand of GQ. The schemes of all systems with the nomenclature specifying the number of G-quartets “nQ” in each system and the strand termini indexing from 1 to 8 are presented in Figure 2. All models (as PDB files) are attached in Supporting Information.

**Figure 2.**
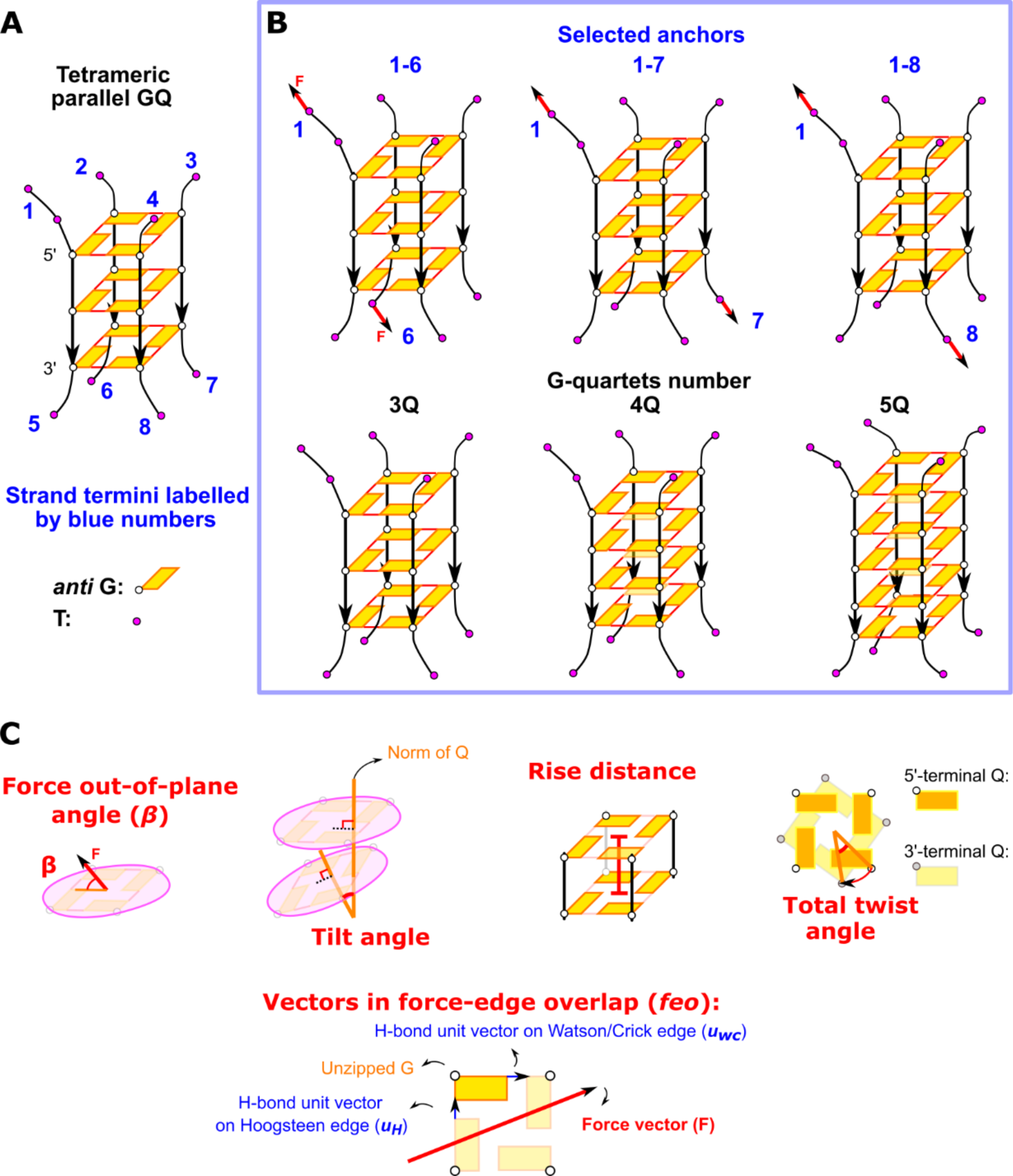
Starting GQ structures and parameters characterizing structural changes. **(A)** Labelling of strand termini. **(B)** Designation of SMD simulations based on pulled strands (pulling anchors) and on the number of quartets in the GQ models. The pulling anchors in the text are specified by superscript, i.e., an abbreviation 3Q^1–8^ means a three-quartet GQ with force-anchoring positions 1 and 8. **(C)** Visual representations of force out-of-plane angle, monitored GQ structural parameters (tilt, rise and twist), and vectors used in the calculation of force-edge overlap (*feo*). *Feo* measures alignment of the pulling force with direction of H-bonds and the formula for its calculation is given in the text.

### Steered Molecular Dynamics simulations

The starting structures were solvated in a truncated octahedral periodic boundary box with OPC water,^112^ where the minimum distance from GQ to the box boundary was at least 5 nm considering the scale extension of unfolded GQ. We also tested the SPC/E water^113^ model for the 3Q-GQ (three-quartet GQ) and 4Q-GQ systems in order to investigate possible impacts of the solvent models. Cl^−^ and K^+^ ions with Joung&Cheatham parameters^114^ were added to the boxes to neutralize the systems and reach the physiological KCl concentration (∼0.15 M). We employed the AMBER OL15 force field.^72,115,116^ The systems were prepared by the Leap module of AMBER18.^117^ Next, we performed initial relaxation and thermalization (details in Supporting Information). Topologies and coordinates after equilibration were converted into GROMACS inputs using the ParmEd module.^117^

SMD simulations were run with three replicates for each system using GROMACS-2018^118^ in combination with Plumed-2.5.^119^ We used Particle Mesh Ewald summation^120^ to calculate the electrostatic energy, and the distance cut-off for non-bonded interactions was set to 1 nm. The velocity rescaling algorithm^121^ and Parrinello-Rahman method^122^ were applied for temperature and pressure control, respectively. SHAKE algorithm^123^ and Hydrogen Mass Repartition method^124^ were used for stabilizing the motions of hydrogens, enabling the 4 fs timestep in the simulations. Pulling was specified in PLUMED as a dynamic harmonic distance restraint between the two spring anchors, defined as geometric centers of C2, C4 and C6 atoms of the two corresponding terminal Ts (see Figure 2), and with the force constant equal to 180 kJ mol^-1^ nm^-2^. All SMD runs used ∼5.4 nm/µs constant pulling speed specified by the dynamic harmonic distance restraint (Table S1). Pulling range and pulling timescale were designed to allow complete detachment of a strand from the GQ. Period between the first unfolding event, which alters H-bonding in the GQ stem, and the detachment of any strand was defined as the unfolding period. The unfolding pathways consisted of series of consecutive individual unfolding events. Maximum averaged force (running average over 2 ns window) ahead of the first unfolding event was recorded as the GQ rupture force. Likewise, we recorded the averaged force inducing each subsequent unfolding event along the pathway.

### Unbiased MD simulations

We extracted important GQ intermediates observed during SMD simulations for subsequent relaxation in standard unbiased MD simulations. Extracted intermediates are listed and shown in Table S2 and Figure S1; all of them had experienced at least two transitions from the native GQ. The GQ coordinates from the SMD trajectories were re-solvated with OPC water in a truncated octahedral periodic box with the distance from the solute to the boundary equal to 1.2 nm. The SPC/E water was loaded for some cases to investigate the effects of different water models. The AMBER OL15 force field^72,115,116^ was used and additional K^+^ and Cl^−^ were added for neutralization and to reach the physiological ionic strength (∼0.15 M). The systems underwent equilibration before the production stage (see Supporting Information), which was performed in AMBER18 using pmemd.cuda.^117^ Two replicates were run for each extracted structure, 1-µs long each.

We also ran standard MD simulations of the native 3Q-, 4Q- and 5Q-GQs in OPC water, two replicates 1-µs long each, to inspect the GQ structural parameters in the unbiased systems. Table S2 summarizes all standard MD simulations performed in this study.

### Definitions of force out-of-plane angle and GQ structural features

We used a set of structural parameters to monitor the pulling force and GQ conformation during the simulations (Figure 2C). **Force out-of-plane angle (*β*)** is defined as the angle between the spring force (given by the spring anchors) and the average GQ quartets’ plane. **Planarity (*P*)** of a quartet is the RMSD value of the distances (*z_i_*) of all heavy base atoms in the quartet from its reference plane (eq. 1); the reference plane is calculated by singular value decomposition (SVD) with the coordinates of bases’ heavy atoms,

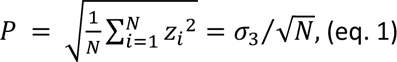

where *N* is the number of heavy atoms, *z_i_* is the distance of the *i*^th^ atom from the best fit quartet plane, and *σ*_3_ is the third singular value. **Tilt angle** is the angle between the two norms of neighbouring quartets, and **rise distance** refers to the distance between their planes, calculated as a sum of distances of the two quartets’ geometric centers to the two quartets’ average best fit plane. **Total twist angle** is the relative rotation angle of the 3’-quartet to the 5’-quartet around the GQ axis, reflecting the (right-)handedness of the GQ helix. We also calculated **force-edge overlap (*feo*)** (difference between two overlaps, to be more precise) before each initial unzipping event; we take the absolute dot product of the pulling force vector (***F***) and H-bonds unit vector on either Watson/Crick or Hoogsteen edge of the (first) unzipped G (***u_wc_*** or ***u_H_***) and subtract the two, then scale the difference by the force magnitude, 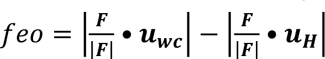. For further details on the calculation of the parameters, see Supporting Information.

### Trajectory analyses

The structures and trajectories from both SMD and unbiased simulations were visualized by PyMOL (version 2.0 Schrödinger, LLC), VMD (version 1.9.3)^125^ and UCSF Chimera.^126^ The force out-of-plane and GQ conformational parameters were calculated by the MDAnalysis^127,128^ and Numpy^129^ Python packages. The K^+^ binding site occupancy was calculated as a percentage of frames having K^+^ between the two adjacent Qs (analysis performed by cpptraj^130^ and an in-house program). The data processing and visualization were done in RStudio.^131^

## Results

### Overview of unfolding transitions

Figure 3 summarizes the most common unfolding transition steps seen in the present study based on their characteristic structural changes, namely unzipping, strand slippage, opening, detachment, formation of (partial) spiral conformation, and rotation. We also observed some other transitions not presented in the figure because they were either too infrequent or too complex to have understandable visualization. The detailed descriptions of all transitions are summarized in Supporting Information.

**Figure 3.**
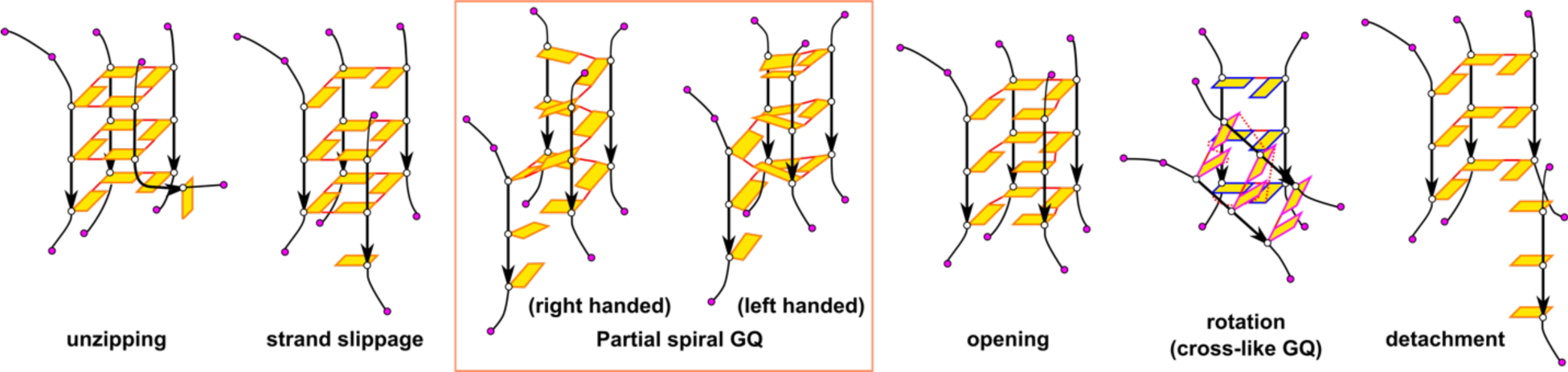
Diagram of the common GQ unfolding transitions. The conformational changes in the figure are shown using the model of 3Q-GQ. The labelling of anti-G and T bases are consistent with those in Figure 2. See the text for detailed descriptions of these transitions.

Unzipping is the most common transition, occurring in all simulated structures. Beside that the unfolding expectedly began by unstacking of terminal Ts that were used as force anchors, rupture process of the actual GQ stem was always (except for one case) initiated by unzipping of a G from GQ terminal quartet, regardless of the pulling directions or the number of quartets in the stem. The initial G unzipping was mostly (36 out of 45 simulations) initiated from the 3’-end of the GQ. The transitions involving changes in multiple Qs usually occurred later (Figure 4). Among them the strand slippage appeared sporadically, but its “halfway-stopped” variant ending in the (partial) spiral GQ conformation was rather common; interestingly, about 40% of the spirals were left-handed and 60% right-handed. The spiral conformation can eventually finish the strand slippage event, or can progress elsewhere, e.g., to unzipping. Of note, only a few minor refolding events were observed in the whole pulling simulations set (Figure 4). Supporting information summarizes the overall statistics of the different transitions (Table S3) and sequence of all transitions in all individual simulations (Tables S4 and S5).

**Figure 4.**
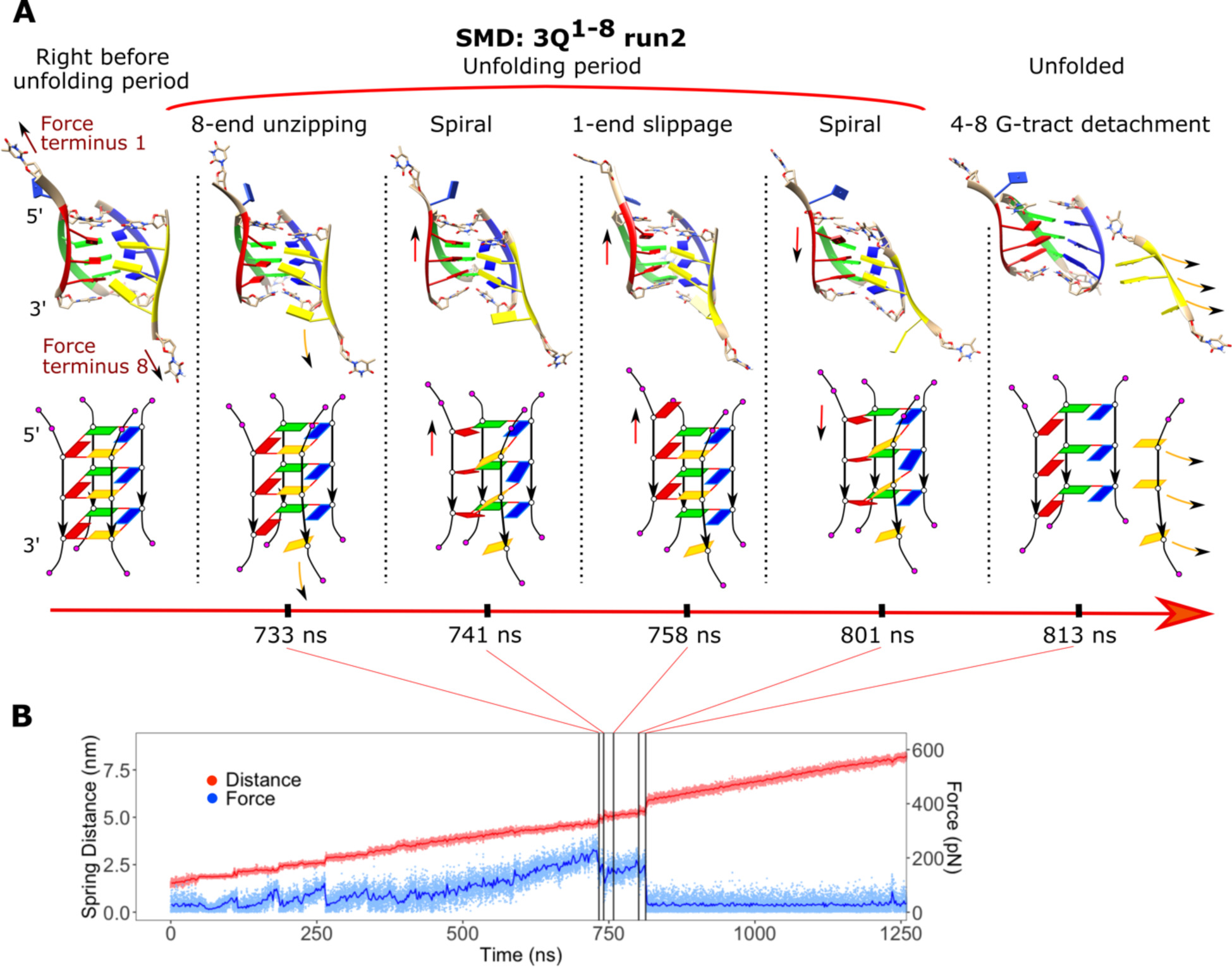
Example of GQ unfolding and the rupture forces. **(A)** Unfolding transitions of the second run of the 3Q^1–8^ system during SMD simulation. The structures are shown both as a cartoon model and simplified scheme. Each strand has a unique colour. **(B)** The distance between pulling centers and the exerted force along the mechanical unfolding; occurrence of the events in (A) is labelled by the grey vertical lines. Force/extension-time graphs of the other SMD simulations are shown in Figure S2-S4.

### 1-8 pulling is most diverse

Qualitative outcome of the unfolding process was influenced by the position of pulling anchors. The most varied outcome of the simulations was observed for the 1-8 pulling (Supporting Information Tables S3–S5). Unzipping was most common at the first stages, but later strand slippage and opening combined with formation of cross-like GQ occurred relatively frequently. On the other hand, GQ unfolding in 1-6 and 1-7 pulling was rather uniform, with the unzipping mechanism being dominant. Nevertheless, even in these pulling directions strand slippage was possible. Of note, the 1-8 pulling direction corresponds to the pulling on the end of intramolecular parallel stranded GQs, such as the human telomeric GQ or various promoter ones. Note also that in the SPC/E water model, strand slippage occurred more frequently than in OPC (Tables S3–S5).

### Unfolding diversity increases with the number of quartets

The number of quartets present in the GQ was another factor that affected the unfolding pathways. Increasing number of quartets led to higher complexity of the unfolding beyond the simple assumption that one additional quartet would mean an additional unfolding event. The occurrence of more complicated transitions, such as GQ opening or strand slippage, was higher in pulling of the 4Q and 5Q systems (Tables S3–S5). In addition, the transitions in 4Q and 5Q system were not necessarily as “clean” as in the 3Q system, i.e., they could happen only partially, for example, slippage of a single or two G(s) instead of the whole G-column, or formation of a cross-like GQ in just a part of the GQ structure.

A G-triplex often remained after a GQ was pulled apart. Three-layered triplexes tended to transition into the cross-like triplex, which has one strand rotated with respect to the other two; it was sometimes occurring even when the GQ was not yet fully unfolded, i.e., the pulled strand was not fully detached. On the other hand, four-layered G-triplexes displayed remarkable stability (five-layered G-triplex was not formed in the simulations). Notice that while the pulling was technically still active, no pulling force was effectively exerted on the remaining triplex state, so it behaved rather as in a standard unbiased MD simulation.

### Transition-inducing force does not scale linearly with the number of quartets

The magnitude of force needed to promote the GQ unfolding varied with both the number of quartets in the GQ and with the pulling direction. The highest transition-inducing forces were in general required in 1-6 pulling direction, followed by 1-7 and 1-8, as evidenced by the maximum forces recorded in individual runs (433, 317, and 252 pN, respectively, on average among replicates), as well as by their general distributions (Figure 5A). The order was the same regardless of the number of quartets, and interestingly, we did not even observe a clear dependence of the rupture force magnitude on the number of quartets. While the 3Q GQ was weakest in general (∼320 pN on average), forces needed to disrupt the 4Q and 5Q systems were on par (∼338 and ∼344 pN, respectively). Examination of the transition-inducing forces in relation to the number of remaining quartets revealed that the transitions originating from 5Q GQs require smaller force compared to GQ stems with three or four remaining quartets (Figure 5D).

**Figure 5.**
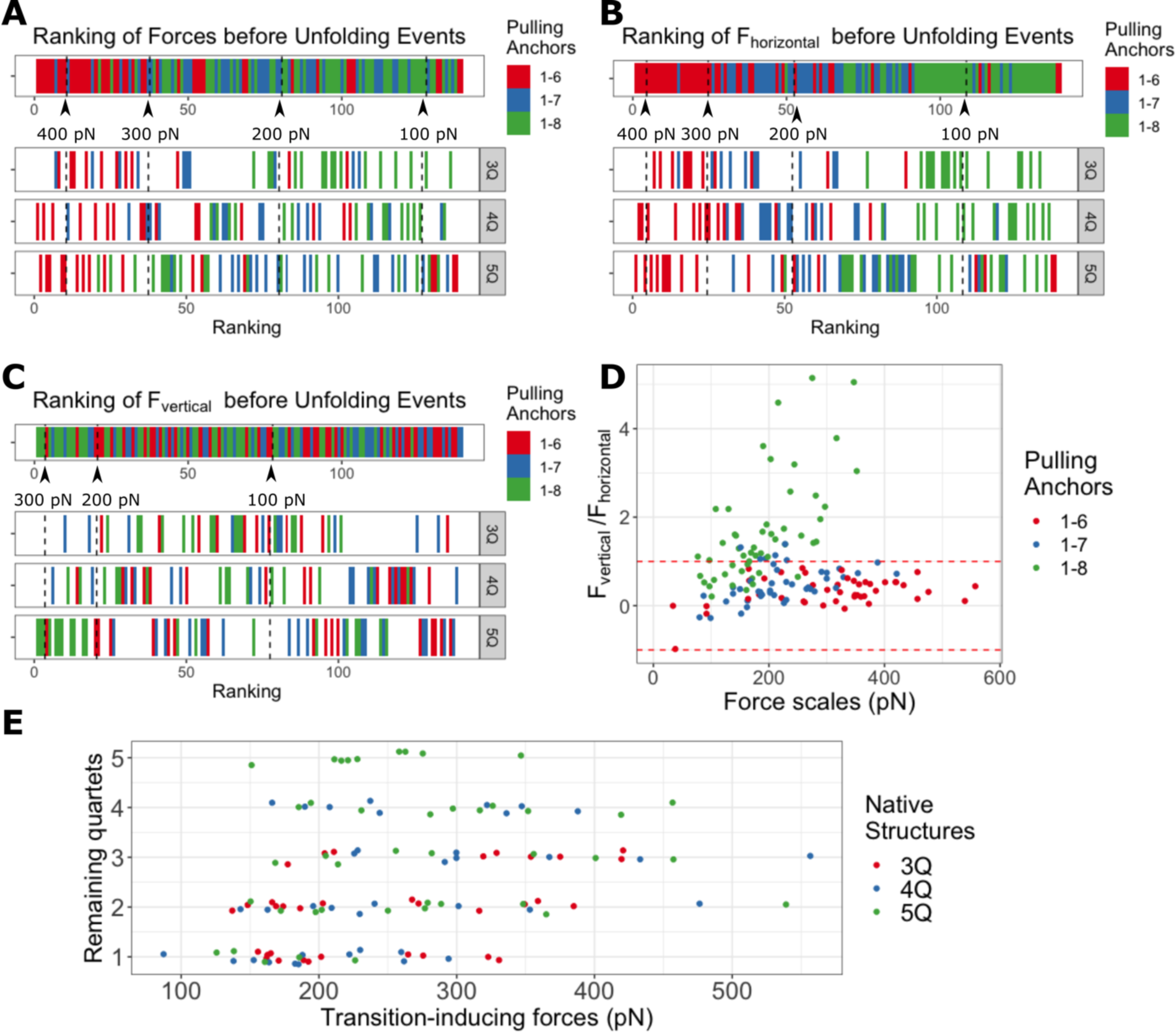
Comparison of transition-inducing forces in simulations with OPC water model. **(A)** The ranking of all force magnitudes (ordered from the highest to the smallest). The figure includes all transition-inducing forces, colored by the three pulling anchor pairs; the first row captures all events in all simulation while the three remaining rows show decomposition of the first row for GQs with different starting numbers of quartets. **(B)** The ranking of the horizontal components of all transition-inducing forces. **(C)** The ranking of the vertical components of all transition-inducing forces. **(D)** The force distribution in the dimension of force magnitude and the vertical-to-horizontal force components ratio. The region between the red dashed lines has the horizontal force component higher than the vertical one. Note that the data with negative ratio have the relative positions of 5’/3’-anchors upside down with respect to the reference GQ plane. **(E)** The distributions of transition-inducing forces under different number of remaining quartets, i.e. quartets actually present in the structure right before the transition, not in the GQ at start of the simulation. The data points are colored according to the native (initial) GQ structures.

Decomposition of the force into its horizontal (parallel to the average GQ plane) and vertical (orthogonal to the plane) component provides further insights. The force components were calculated from the total force and the force out-of-plane angle β (Figures S5–S7) by trigonometric formulas (*F_horizontal_* = *F* × cos *β*, *F_vertical_* = *F* × sin *β*). The vertical component dominates during the events happening in 1-8 pulling, while in 1-6 and 1-7 pulling it is smaller than the horizontal component (Figure 5). This observation is documented by the distributions of all recorded forces in the dimension of the force magnitude vs. the ratio of the vertical and horizontal component (which is equal to the tangent of the angle of force incidence). The events in 1-6 and 1-7 pulling, in Figure 5D, are distributed nearly along lines with small ratio values, while the 1-8 pulling data lie in a different region, where the vertical force component is larger than the horizontal one.

### Elastic deformations of the GQ helical structure ahead of the first unfolding event

Monitoring of GQ structural parameters during the simulation reveals that the force firstly induces conformational changes finer than rupture events. Specifically, the pulling work is absorbed by elastic deformation of the GQ helix. The most prominent is total helical twist decrease in conjunction with helical rise (quartet-quartet vertical distance) increase (Figure 6A,B,C and S8-19, Table S6). The extent of the effect depends on the position of the anchors relative to the GQ center, i.e., both the number of quartets and pulled strands play a role simultaneously (Figure 6A,B and S8-S19). We observed that the deformation was most profound for 1-8 pulling, showing largest helical unwinding. On the contrary, the 3Q^1–6^ system underwent only small changes, because the force direction promoted helical overwinding, which appears to be opposed by stacking interactions and the backbone. The reversibility of the elastic deformation was manifested when the tension was relieved; for example, after an unzipping event the remaining part of the GQ becomes stress-free and returns to the standard helical form.

**Figure 6.**
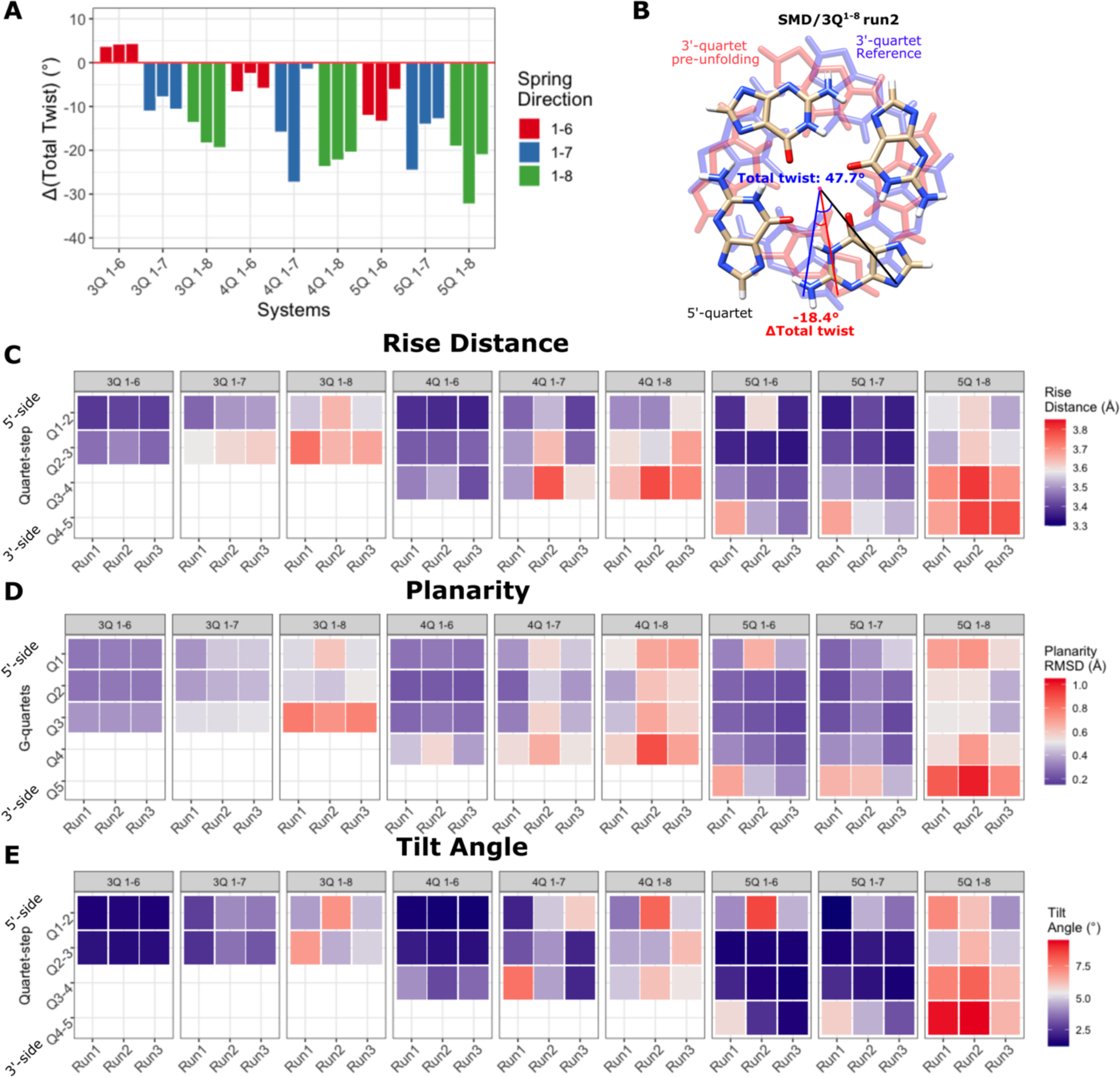
GQ conformational features right before the first unfolding step occurred (simulations in OPC water) **(A)** The changes of total twist angles in all SMD simulations from the equilibrium GQ structure to the GQ ahead of the first unfolding event. **(B)** An example of a total twist angle change from the equilibrium structure to the 2 ns-averaged value right before the unfolding period, taken from the 3Q^1–8^ system simulation. The rise distances **(C)**, planarity **(D)**, and tilt angles **(E)** for all quartet-quartet steps in SMD simulations. See Tables S6-S8 for exact values and the corresponding values in standard MD simulations as references.

### G-quartet deformations and preferred H-bond breaking edge

We further monitored the planarity of individual quartets. The highest non-planarity was seen at 3’-terminal quartet in the 1-8 pulling simulations (Figure 6D, Figure S20-28, Table S7), and it was mostly manifested as a buckled base in the pulled strand. The buckled base also dragged the rest of the quartet, so the whole quartet was tilted with respect to the neighboring quartet (Figure 6E and S29-S37, Table S8). This deformation can likely be attributed to the dominating vertical component of the pulling force; in 1-6 pulling, where the vertical component is less dominant (Figure 5), such quartet deformations were minimal.

In addition to the GQ helical and quartet step parameters, the force direction relative to the GQ center influenced the preference of G base edge where H-bonds would break during the initial unzipping event. The more the force was parallel to the H-bonds on one edge (either Watson-Crick or Hoogsteen), the more likely was it to break there first. (Figure 7A) For example, 3’-unzipping events in 1-6 pulling usually started by breaking the H-bonds at the Hoogsteen edge (from the perspective of the eventually pulled-out G), while in 1-8 pulling the same situation would result in breaking at the Watson-Crick edge (Figure 7B).

**Figure 7.**
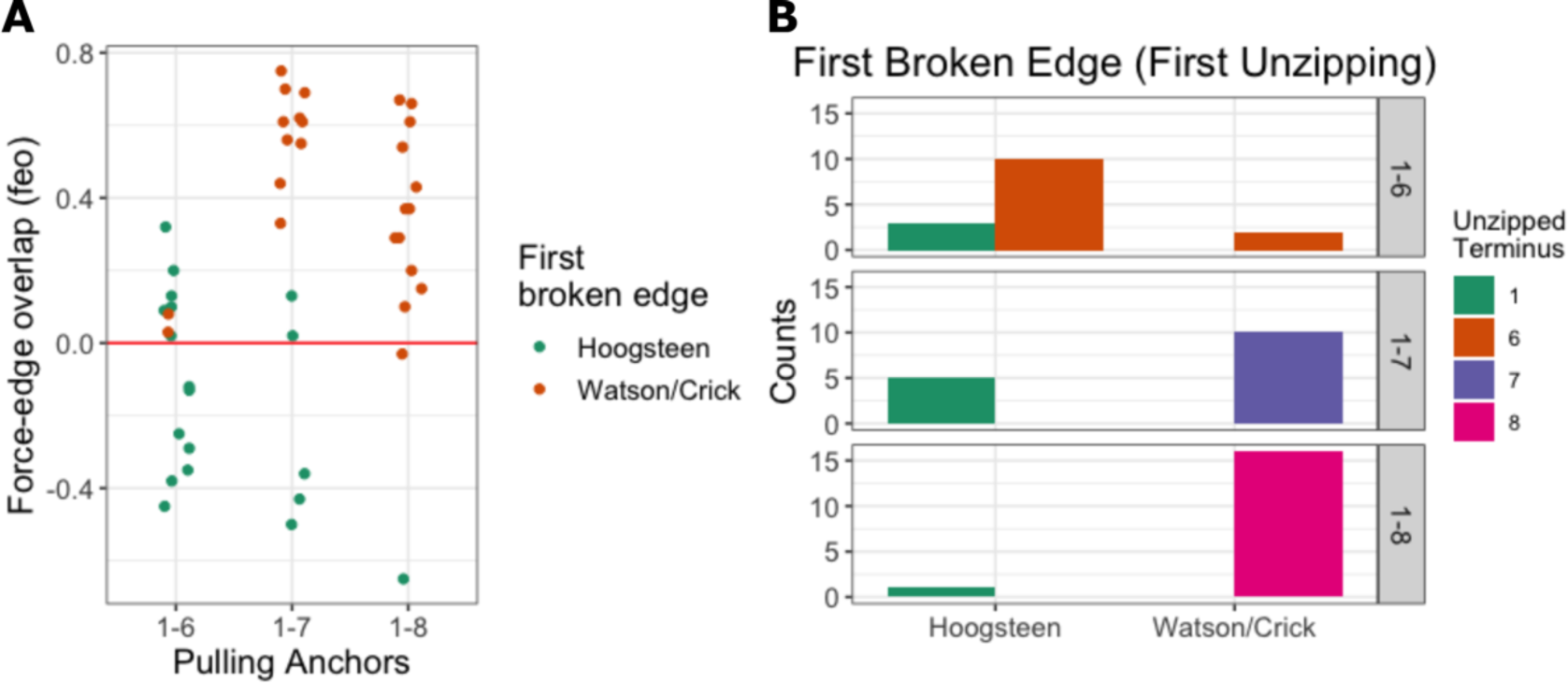
Preference of H-bond breaking edge. **(A)** Force-edge overlap of each unzipped G (see Methods). Positive value means the pulling force was aligned more with the Watson-Crick edge, while negative value indicates better overlap with the Hoogsteen edge. **(B)** Histogram of the first broken edge for all initial unzipping events.

### Partial spiral structure is a transitory ensemble

The bias introduced by the SMD simulations accelerates GQ conformation sampling, but it inherently guides the system via a lower-entropy (restricted) regions of the free energy surface. Thus, to probe the stabilities of some intermediates sampled in the SMD simulations, we performed standard unbiased MD simulations. We mainly focused on the partial spiral conformations since they emerged during strand slippage and unzipping pathways transitions but have not been characterized in detail in previous studies. A few slip-stranded GQs and a structure with rotated strands were also picked up for unbiased simulations (Table S2).

Most importantly, the unbiased MD simulations validated the transient nature of the partial spiral structures sampled by the SMD simulations (Table S9). All the spiral states vanished shortly after the start of the simulations and the spiral state had lifetime up to dozens of nanoseconds. In addition, the partial spiral conformation was – again – sampled for short periods of time during strand slippage events. Both left-handed and right-handed spiral conformations appeared in the unbiased simulations.

Among the total 24 unbiased simulations starting from the spiral conformations, only one unfolded, while there were 10 runs refolding to the native GQ structure; the rest remained partially unfolded. Interestingly, the two independent replicates of the same Spiral-3Q^1–6^-run3 system evolved into completely opposite directions, with one refolded into the native 3Q GQ and the other unfolded (Figure 8). The unfolding development included strand slippage and opening before the strand 1-5 detached from the GQ stem. The refolding simulation showed an intricate process, with multiple strand slippage events and incorporation of two unzipped Gs; the strand slippages could be reversible and not always towards the native GQ conformation.

**Figure 8.**
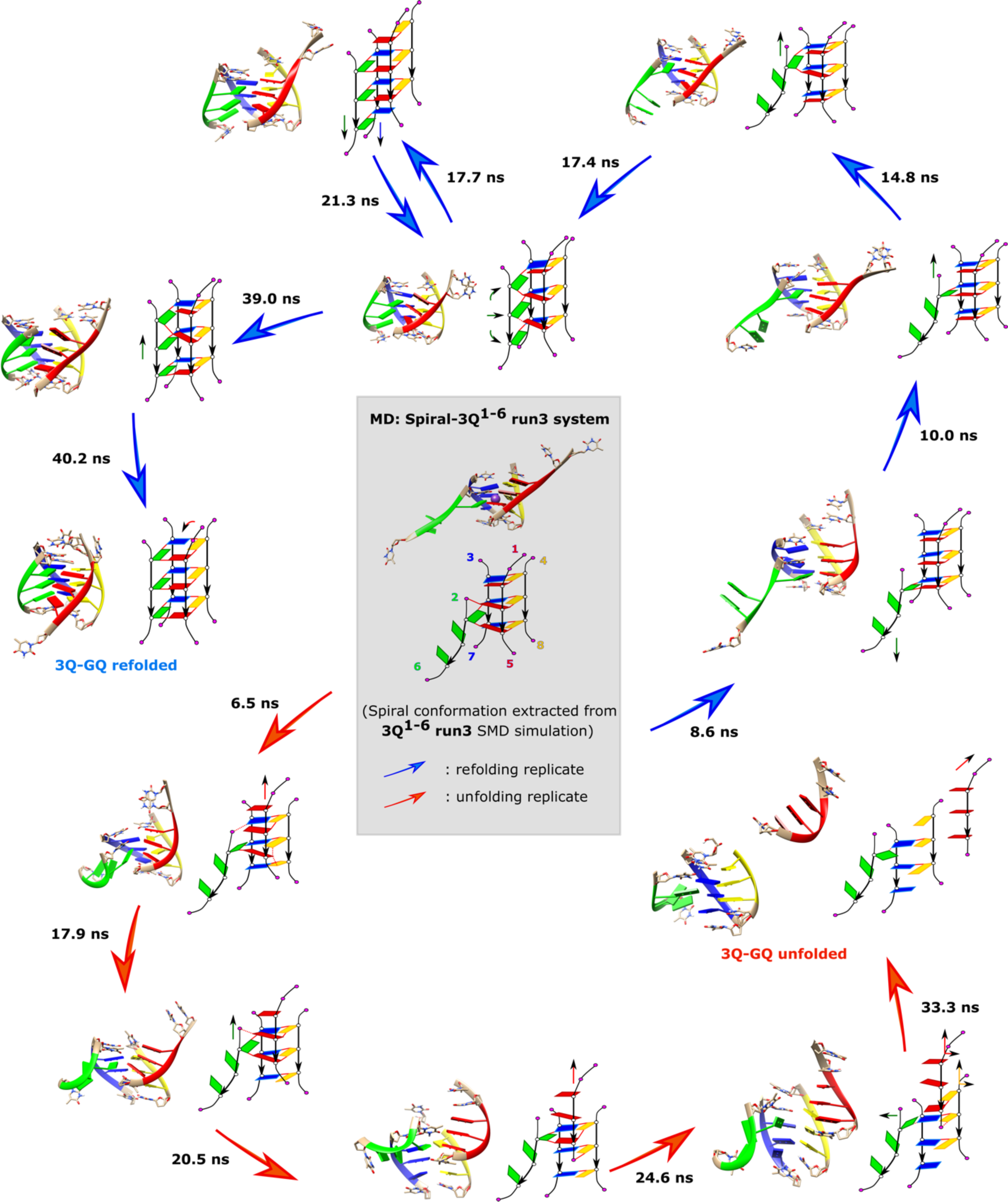
The folding and unfolding transitions in the unbiased simulation of Spiral-3Q^1–6^ run3 system. The starting structure of the simulations is a slip-stranded left-handed spiral conformation, shown in the center of the scheme in grey background. The two replicates show opposite directions of conformational change: one refolds to the original 3Q-GQ conformation (pathway with blue arrows) and the other unfolds since the 1-5 strand detached (pathway with red arrows). The times of the transitions are labelled next to the arrows.

On the contrary, we did not observe significant unfolding or folding tendency in the simulations starting from the slip-stranded and strand-rotated intermediates, the simulations remained locked around the starting conformation (see Supporting Information).

### Water model effect and cation jumping-out events

We performed 27 pulling simulations in the OPC water and 18 (for 3Q and 4Q GQ) in the SPC/E water model. Under the same pulling conditions (position of pulling anchors, number of Qs), the SMD simulations performed in OPC and SPC/E water models were rather similar and showed only minimal differences in most aspects (Cf. Figure 6 and S38); unzipping was the dominant unfolding transition, with strand slippage been slightly more likely in SPC/E water (Table S3). The rupture forces in both water models were comparable (Cf. Figure 5 and S39). 4Q and mainly 5Q GQs suffered from escape of usually one cation from the 3’-peripheral side of the channel. We noticed slightly smaller occupancy of the channel cation binding sites in SPC/E water model (Figure S40). Nevertheless, in SPC/E the terminal quartet was typically stable even without a cation in the adjacent channel binding site. On the other hand, although the cations had higher occupancy in OPC water, once the cation escaped from the channel, this event was more likely to be closely followed by base unzipping. The ion escapes from 4Q parallel tetramolecular GQs in standard simulations was noticed already earlier and was attributed to the exaggerated inter-cation repulsion in pair-additive force fields.^78^ Once a G base was unzipped, it could form an intercalated structure with the flanking Ts, which we suppose might be a force field artifact. We observed higher occurrence and stability of such structures with the SPC/E water model. Formation of such an intercalated intermediate was the reason why one SMD simulation in the SPC/E water did not reach complete strand detachment on the simulation time scale. See Supporting Information for further details.

## Discussion

We investigated GQ mechanical unfolding by slow Steered Molecular Dynamics (SMD) simulations, sampling the unfolding intermediate conformations by mechanically pulling tetrameric parallel GQs, i.e. parallel all-*anti* G-stems in absence of loops. We have systematically studied GQs with 3 to 5 G-quartets while testing all three possible dispositions of the anchored force spring. Altogether we performed 45 pulling simulations. The simulations showed that the complexity of unfolding dynamics and intermediates sampled by SMD simulations depends on both the number of quartets and the pulling anchors used. Unzipping is the main (and initial) unfolding transition under all the setups, but the whole spectrum of observed structural transitions is comparable to pulling of intramolecular GQs reported by us earlier.^107^ We observed substantial quantitative differences in rupture forces among the GQ systems and tested force directions, which means that the response of the GQ stem to the pulling force is anisotropic. Overall, the simulations confirm that at the atomic resolution the unfolding of GQ under external force consists of a series of structural transitions, rather than being a sudden one step rupture of the structure. This aspect of the unfolding process is likely below resolution of the experimental techniques.

### The roles played by different types of transitions in GQ unfolding: the dominant role of G unzipping

Unzipping is the fundamental unfolding transition during mechanical pulling of tetrameric parallel GQ. It is the most common transition. All (except of one) SMD simulations initiated the unfolding process by unzipping of at least one G while unzipping played a role also in the later stages of the pulling. Strand slippage (register shift) played only a considerably lesser role. This was somewhat surprising, since importance of strand slippage in (un)folding of parallel stranded GQs was suggested in several preceding studies.^77,92,106,107^ In the present work most of the strand slippage events occurred when the spring-anchor strand had only two Gs attached to the GQ body, requiring less rebuilding of the H-bonds network. This result does not mean that strand slippage is unimportant for GQ (un)folding in absence of external force, e.g. in thermal unfolding. However, based on the present results we suggest that the strand slippage may be facilitated by a preceding perturbation of one of the terminal quartets.

Besides the role of the number of remaining quartets in the structure, with smaller number increasing the chances of strand slippage, the simulation results appear to be also sensitive to the used water model. Based on the relative distribution of strand slippage events in OPC vs. SPC/E water models, we suggest that the choice of simulation water model affects the balance between unzipping and strand slippage. The OPC favours more unzipping whereas SPC/E (and TIP3P^132^) water model makes the strand slippage somewhat more likely, though G-unzipping is still the first conformational transition (except of one SPC/E simulation in which the unfolding started directly by strand slippage). This effect of water model would explain why in previous standard MD,^77,92,133^ enhanced-sampling MD,^134^ as well as SMD simulations,^106,107^ strand slippage was quite frequent for parallel-stranded GQs (some with loops) in SPC/E and TIP3P water; on the other hand, in OPC spontaneous base unzipping (unbinding) occurred even under no force^78^. Admittedly, we expect that the balance is also affected by the cations, and we reiterate that all simulations with pair-additive force fields inevitably suffer from overestimation of cation-cation repulsion in the stem.^78,135,136^ Nevertheless, the picture emerging from the present SMD simulations is that the first unfolding event is typically unzipping (unbinding) of one or more Gs in the terminal quartets, which opens the door for further structural changes (cf. Figure 3 Figure 4). Supporting information Tables S3-S5 list the statistics and sequence of all unfolding events in all 45 pulling simulations and clearly demonstrate the dominant role of G-unzipping in the unfolding process.

Similar to strand slippage, the other non-unzipping transitions (Figure 3) also occurred after one or several G-bases had unzipped during pulling, suggesting that the breakage of one quartet to a triplet is a bottleneck process of GQ unfolding which facilitates subsequent structural distortions involving even multiple quartets. Overall, despite a few differences, unfolding of the tetrameric GQs by external force shows structural richness comparable to the intramolecular telomeric GQ.^107^

### The role of spiral-like structures

In standard MD runs starting from partially unfolded GQs, unzipping, incorporation of unzipped Gs, and strand slippage were also frequently observed. We propose that these are fundamental late-stage GQ transition folding movements sampled even without the assistance of external forces. Importantly, herein we examined in more detail the partial spiral conformation (Figure 3 Figure 4). We found out that it may serve as a transition ensemble during strand slippage and unzipping, as well as in refolding events despite its relatively short lifetime (up to 5 ns) in both SMD and standard simulations. Its appearance in both standard and SMD simulations, in both folding and unfolding pathways, suggests that it is a relevant transitory ensemble at a crossroad of multiple conformational transitions.

### Anisotropic mechanical stability of tetrameric parallel GQs

The simulations revealed remarkable differences in the GQ mechanical stability depending on the force direction, which is mostly determined by the spatial arrangement of pulling anchors. In general, disruption of a GQ required highest force in 1-6 pulling, whereas smallest in 1-8 pulling. Explanation of this phenomenon requires delving deeper into the GQ structure. Horizontal and vertical components of the acting force, i.e. parallel and perpendicular to the average quartet plane, respectively, vary with two factors: i) position of the anchors and ii) the number of quartets, which determines total GQ helical twist (approx. +30 degrees per quartet-quartet step). The forces first induce elastic changes in the GQ structure and when the GQ cannot sustain the strain it undergoes a local rupture event. The horizontal force component can act on the GQ by a full range between strong unwinding and strong overwinding, depending on the mutual position of pulling anchors (see Supporting Information text and Figure S41). This is similar to unwinding or overwinding of a rope by twisting hands, in which unwinding disrupts the GQ structure while overwinding makes it stiffer. Vertical force component indirectly causes only unwinding; although there is no twisting effect of the force per se, the limited flexibility of the backbone must respond by unwinding to the increasing inter-quartet distance in vertically extended helix. In the 1-6 pulling, the horizontal force component dominates, and results in both overwinding and underwinding depending on the number of quartets. In the 1-8 pulling, the horizontal force can also induce unwinding or overwinding, but the vertical component is larger than the horizontal one and consequently induces strong unwinding very efficiently regardless of initial anchors spatial arrangement. The vertical stretching of the GQ is also correlated with major quartet deformations, such as increased quartet non-planarity (base buckling), which in turn weaken quartets and their stacking. As a result, the vertical component dominating the 1-8 pulling is responsible for the major changes of GQ conformation and the smaller total force needed to disrupt the GQ in comparison to the 1-7 and 1-6 pulling. We thus hypothesize that the significant elastic GQ deformations induced – and determined – by the vertical force component lead to the reduction of total force magnitude required for GQ mechanical unfolding. Similarly, strand slippage is a vertical transition, thus it is associated with the vertical force component, which explains why it was more likely to occur in the 1-8 pulling.

Interestingly, no linear correlation between the number of remaining quartets and GQ mechanical stability was observed. Magnitudes of transition-inducing forces were similar in systems with two, three and four remaining quartets, whereas GQ with five quartets was weaker (Figure 5E). We think the reason is that unzipping – the most common transition – acts relatively independently of the number of quartets, therefore the similar force magnitude is seen among the GQs. The lower stability of 5Q GQ can be explained as a consequence of the channel-cation expulsion artefact (due to over-estimated cation-cation repulsion, see above), which was common in the 5Q GQ simulations and which mostly affected the initial 5Q state. It should be noted that stability of G-quartets is not uniform; no unzipping ever happened in the inner quartets when they were still covered by outer quartets, and the 3’-terminal quartet was more labile that the 5’-terminal one. We link the smaller stability of 3’-quartets to their greater intrinsic structural deformation (Figure 6), which likely weakens both stacking to the neighbouring quartet as well as H-bonding and ion binding within the quartet itself. In experiments, more quartets in a GQ are in general associated with greater thermal and kinetic stability.^54,55^ The apparent discrepancy between such measurements and our simulations stems from different parts of the free-energy surface being explored; the SMD simulations (or pulling experiments) use a chosen collective variable for pulling (end-to-end distance), which restricts the conformational sampling. The SMD simulations thus follow a rather low-entropy pathway. On the other hand, melting experiments are a high–entropy technique, so the GQ is allowed to explore different parts of the free-energy surface during its denaturation. Obviously, force-field artifact (the above-noted over-estimated cation-cation repulsion which increases with the number of ions inside the stem) may also affect the simulation result.

The 3Q tetrameric GQ is a basic scaffold of many biochemically relevant intramolecular GQs, which differ only by the presence of propeller loops. In this sense, the 3Q^1–8^ pulling simulations conducted in SPC/E water resemble the human telomeric parallel GQ (PDB ID: 1KF1^137^) pulling investigated in our previous study.^107^ Notably, the rupture forces measured in 3Q^1–8^ are larger than those in 1KF1 pulling (∼200 pN in tetrameric parallel GQ vs. ∼160 pN in 1KF1 GQ). Even though the SMD protocols differ slightly between these two studies, we think that the propeller loops in 1KF1 weaken the mechanical stability of GQ in simulations. This is obviously a counterintuitive result. We suggest that it may be attributed to imbalance of the force field which spuriously destabilize the propeller loops. It has been suggested by several preceding studies using standard as well as enhanced sampling simulations, where instability of the propeller loops was also observed and attributed to a force-field artefact.^71,138,139^ The origin of this imbalance has not yet been identified.

## Conclusions

We employed slow pulling SMD to investigate the unfolding of tetrameric parallel G-quadruplexes (GQs), with a focus on the impact of the number of quartets (3-5) in the GQ and the pulling force direction. The slow pulling SMD protocol allowed to reveal details of transitions and intermediates of tetrameric parallel GQ unfolding, which would be difficult to examine by single molecule pulling experiments.

All GQ systems exhibited a stepwise mechanical unfolding mechanism comprised of common classes of transitions, namely base unzipping, strand slippage, opening and rotation to cross-like structures. Unlike in the intramolecular GQs, the tetrameric GQ unfolding is dominated by unzipping movements. Unzipping of single base in one of the outer quartets triggers the other types of transitions. Interestingly, the unzipping was preferred at the 3’-end over the 5’-end. Strand slippage is identified as less pronounced than unzipping but still crucial. Before the first unfolding event (the first unzipping) and between subsequent transitions we evidence elastic changes in the GQ structure, typically unwinding of the helical structure and deformation of quartets. We investigated in more detail conformational properties of short-living partial spiral conformations, which occurred as an intermediate during strand slippage and some unzipping events, suggesting that spiral-like structures play a role of transition ensembles.

The unfolding dynamics and mechanical stability of the GQ are significantly affected by the spatial arrangement of pulling anchors with respect to the G-stem, which is determined by the selection of pulling anchors and the number of quartets in the GQ. We suggest that the pulling force with a larger vertical component (i.e. perpendicular to the average quartet plane) disrupts the planarity and stacking of the quartets more efficiently than forces with smaller vertical component. These deformations of GQ structure result in i) increased occurrence of transitions involving multiple quartets (strand slippage) and ii) smaller force required for GQ rupture. These results illustrate the anisotropic character of mechanical unfolding of tetrameric parallel GQs.

In summary, the results provide further evidence for the complexity of GQ structural transitions at the atomistic level of resolution. Most importantly, we demonstrate anisotropic behaviour of GQ and explain how and why different forces induce different changes. Our work broadens the overall understanding of GQ structures and their dynamic changes and properties.

## Associated Content

### Data Availability Statement

All data applied on deriving the results and discussions are available in this text and in the Supporting Information. The PDB files of simulation starting structures are documented in Supporting Information. The simulation input and parameter files, including PLUMED input files defining the pulling protocols, and the force and anchor distance information used for generating the results have been published in Zenodo repository (https://zenodo.org/records/10527801). The AMBER18 package and OL15 force field can be licensed and downloaded from the AMBER (http://ambermd.org/) and OL Force Fields (https://fch.upol.cz/ff_ol/downloads.php) official webpages. The GROMACS-v2018 (https://www.gromacs.org/) and PLUMED-v2.5 (https://www.plumed.org/download/) are available for free. The PyMOL Molecular Graphics System can be licensed from Schrodinger (https://pymol.org/2/). The VMD molecular visualization program can be licensed from UIUC (http://www.ks.uiuc.edu/Research/vmd/). The UCSF Chimera program can be licensed from UCSF (https://www.cgl.ucsf.edu/chimera/).

The trajectories underlying this article will be shared on reasonable request to the corresponding author due to the large size of the raw simulation trajectories.

### Supporting Information

The starting structures of tetrameric parallel G-quadruplexes used for both SMD and standard MD simulations.

More details of simulation protocols, structural parameters of G-quadruplex, unfolding transition, and additional simulation data and analyses.

## Author Information

### Author Contributions

The project discussion and manuscript writing were contributed by all the authors (Z.Z., V.M., M.K., J.S., P.S.). All the authors approved the final version of the manuscript.

### Notes

The authors declare no competing financial interest.

## Supporting information

Starting structure in PDB, compressed into "tar.gz"

## Acknowledgement

This work was supported by the Czech Science Foundation (21-23718S).

## Notes

https://zenodo.org/records/10527801

### Competing Interest Statement

The authors have declared no competing interest.

