## Supplementary text, figures, and tables for "Mechanical stability and unfolding pathways of parallel tetrameric G-quadruplexes probed by pulling simulations"

### *Equilibration protocol*

The equilibration was done in AMBER18 for both SMD and MD simulations.[1] The initial minimization of the system was done while applying a 25 kcal/(mol•Å$^2$) positional restraint on the GQ atoms. The systems were then heated up from 100 K to 300 K while keeping the positional restraint. Afterwards, an alternating series of minimizations and equilibrations was employed for five rounds. In each round, the positional restraint applied to the solute gradually decreased from 5 kcal/(mol•Å$^2$) to 1 kcal/(mol•Å$^2$) with a step size of 1 kcal/(mol•Å$^2$). For the final equilibration, the positional restraint was 0.5 kcal/(mol•Å$^2$). For each minimization, we ran 500 steps of steepest descent followed by 500 steps of conjugated gradient minimization. The equilibrations were run for 50 picoseconds each.

### *Calculation of force out-of-plane angle and GQ structural features*

The **force out-of-plane angle (*β*)**, the angle between the spring force and the average GQ quartets' plane, is calculated from the norm of the average GQ plane (calculated by singular value decomposition), and the spring anchors' vector. During the unfolding period, the formal "average GQ plane" is estimated from the remaining well-shaped quartets, G-triplets or even a single G-base, which are chosen manually by inspecting the trajectory. Thus, the reference GQ plane is obtained specifically in different cases. For the **total twist angle,** we take the pairs of Gs on 5'-Q and 3'-Q on the diagonal strands and obtain their C1'-C1' and N9-N9 vectors. The corresponding averages of the two vectors on both 5' and 3' Qs are projected to the average plane, from which we compute the projection angle. The total twist angle value is further averaged by the two projection angles resulted from the two combinations of diagonal strands (e.g., on 5'-Q, one combination is termini 1,3 and the other one is termini 2,4). In addition to the instantaneous value of all the parameters, we also calculated the running average over 2 ns.

### *Description of common unfolding transitions*

*Unzipping* refers to the process in which a G-base is dragged off from the GQ stem by breaking the H-bonds in the quartet and losing the stacking interactions with the nearby quartet. *Strand slippage* occurs as a single (rarely two) strand shift towards either the 5' or 3' side direction, which involves the complete loss of the original *cis*-Watson-Crick/Hoogsteen (cWH) H-bonds of the sliding strand and rebuilding of new H-bonds in the adjacent quartet layers. *Opening* happens when two neighbouring G-tracts in GQ lose their mutual H-bonds in all quartet layers, leaving the cation binding sites exposed to the solvent. *Detachment* describes removal of a strand from the GQ, which indicates a significant GQ unfolding transition, and is a special case of unzipping. Detachment is the last transition which leads to the entirely unfolded GQ. *Spiral* conformation appears as an intermediate of the strand slippage, in which either the Watson-Crick side or the Hoogsteen side of one G-tract breaks the original H-bonds and buckles up or down to form H-bonds with the Gs in the neighbouring Q layers. This type of conformations is named "spiral" since the new *c*WH H-bonds connect multiple quartet layers of partially unfolded GQ and reshape the structure like a spiral. It could have either left-handed or right-handed sense. We often use the term *partial spiral*, which encompasses structures not adopting the nice spiral shape fully developed in the G-tract vertical sliding movement. Partial spirals can contain imperfections such as one G of the moving G-tract not slipped at all, or only one new *c*WH bond formed instead of two. *Rotation* into the *cross-like GQ* describes two neighbouring strands mutually rotating against the other two strands.

### *Unbiased simulations of slip-stranded and strand-rotated GQs*

Standard simulations starting from the spiral structures revealed a high tendency to diverse structural rearrangements, towards both folding and unfolding. In contrast, lesser folding/unfolding tendency was observed in the unbiased simulations starting from the slip-stranded GQ intermediates (Table S9). The systems Shifted-3Q$^{1\text{-}8}$-r2 and Shifted-3Q$^{1\text{-}8}$-r3, which had two well-shaped quartet layers, remained in the slip-stranded conformation during the simulations; the 5'-G on the 5'-shifted strand formed stable non-Watson-Crick pairing with the 5'-T on the diagonal strand, hindering the shifted strand to slide back. On the other side, the unzipped 3'-G turned to the *syn*-conformation after the unzipping in SMD, which prevented the incorporation of the unzipped 3'-G into the G-stem. Thus, no transitions were observed in simulations of these two systems. In contrast to Shifted-3Q$^{1\text{-}8}$-r2 and Shifted-3Q$^{1\text{-}8}$-r3 structures featuring well-formed quartet planes, Shifted-4Q$^{1\text{-}6}$-r3 and Rotated-4Q$^{1\text{-}8}$-r1 intermediate structures have large quartet planarity deviations. Simulations starting from these structures were found showing more conformational changes, such as incorporations of unzipped G, which improved the planarity and stacking of quartets. Specifically in the Rotated-4Q$^{1\text{-}8}$-r1 simulations, the rotated strands rotated back and formed three quartets with nearly ideal quartet planarity. However, the conformational transitions in these systems were not sufficient to refold them to the native GQ structures.

### Water model in the MD simulations of GQs

The SMD simulations of tetrameric parallel GQs in OPC and SPC/E water models provided similar results. Specifically, there were no significant differences observed in terms of the diversity and preference of the transition events, the distributions of transition-inducing forces, and the elastic conformational changes between the simulations in the two water models. However, the lower occupation rates of cation binding sites still matter in the SMD simulations with SPC/E water. Although the peripheral quartet remained stable for a long period after the escape of the neighbouring channel cation, the absence of the channel cation would destabilize GQ and influence the evaluation of the mechanical stability of GQs. This is the reason why we did not make SMD simulations for the 5Q GQ system in SPC/E water. We assume that for this system it is very likely that it would suffer from the ion escape from the channel which could undermine reliability of the SMD simulations. Thus, while the previous benchmarking simulations recommended SPC/E water model as a safe choice for standard simulations of GQs,[2] we suggest the OPC water model can be a safer choice for SMD simulations of GQ with four or more quartets. Nevertheless, in summary, despite the above-noted differences, the SMD simulations in OPC and SPC/E water models give basically the same unfolding picture.

### Mathematical model for estimating unwinding or overwinding effects given by the horizontal force

The horizontal force component (related to the average GQ plane) is expected to induce unwinding or overwinding on the GQ being pulled. The mathematical model we developed estimates the torque that determines the unwinding or overwinding of the GQ, given the arrangement of the pulling anchors, i.e., 1-6/1-7/1-8 pulling:

$$T_H = F * r^2 * \sin t \,/ \sqrt{s^2 + 4r^2 \sin^2 {}^{t}/_{2}}$$

where $T_H$ is the torque caused by the horizontal force component, *F* is the total force magnitude, *r* is the distance from quartet geometric center to the edge of sugar (approximated as 1 nm). *s* is the vertical distance between quartets calculated as 0.34×(*N*-1), with *N* equal to the number of quartets, and *t* is the total helical twist adjusted to account in the position of anchors as follows:

$$t_{1-6} = (N-1) * \frac{30\pi}{180} + \frac{\pi}{2}$$
$$t_{1-7} = (N-1) * \frac{30\pi}{180} + \pi$$
$$t_{1-8} = (N-1) * \frac{30\pi}{180} - \frac{\pi}{2}$$

See Figure S41A for graphical representation of the terms. The relationship between the torque ($T_H$) and the anchor arrangement, determined by the number of quartets ($N$) and the anchor points (1-6/1-7/1-8), is shown in Figure S41B. According to the model, 1-7 pulling results in unwinding in all GQ structures we constructed. $3Q^{1\text{-}6}$ system has overwinding effects, while $5Q^{1\text{-}6}$ system induces unwinding. In contrast, 1-8 pulling has unwinding and overwinding effects on the GQs with three and five quartets, respectively.

**Supplementary Table**

**Table S1:** Overview of all SMD simulations.

| System name[a] | Simulation time (μs) × number of simulations | Pulling distance (nm)[b] | Pulling speed (nm/μs) |
|---|---|---|---|
| $3Q^{1\text{-}6}$ | 1.1 × 3 | 3 → 9 | 5.45 |
| $3Q^{1\text{-}7}$ | 1.3 × 3 | 2 → 9 | 5.38 |
| $3Q^{1\text{-}8}$ | 1.3 × 3 | 1.5 → 8.5 | 5.38 |
| $3Q^{1\text{-}6}$-spce | 1.1× 3 | 3 → 9 | 5.45 |
| $3Q^{1\text{-}7}$-spce | 1.3 × 3 | 2 → 9 | 5.38 |
| $3Q^{1\text{-}8}$-spce | 1.3× 3 | 1.5 → 8.5 | 5.38 |
| $4Q^{1\text{-}6}$ | 1.7 × 3 | 2 → 11 | 5.29 |
| $4Q^{1\text{-}7}$ | 1.3 × 3 | 2 → 9 | 5.38 |
| $4Q^{1\text{-}8}$ | 1.1 × 3 | 2 → 8 | 5.45 |
| $4Q^{1\text{-}6}$-spce | 1.3 × 2 | 2 → 9 | 5.38 |
| $4Q^{1\text{-}7}$-spce | 1.3 × 2 | 2 → 9 | 5.38 |
| $4Q^{1\text{-}8}$-spce | 1.1 × 2 | 2 → 8 | 5.45 |
| $5Q^{1\text{-}6}$ | 1.7 × 3 | 3 → 12 | 5.29 |
| $5Q^{1\text{-}7}$ | 1.7 × 3 | 2.5 → 11.5 | 5.29 |
| $5Q^{1\text{-}8}$ | 1.3 × 3 | 2.5 → 9.5 | 5.38 |

[a] The name starts by the number of Qs in the GQ, followed by the spring direction in superscript (see Methods). SPC/E water model was used for the systems with "-spce" label, and OPC water model otherwise.

[b] The pulling distance is defined between the two spring-anchored points (the geometric center of C2, C4 and C6 atoms in each selected T, see Methods). It gradually increases during the simulations from the initial value (the number before the arrow) to the final value (after the arrow).

**Table S2:** Overview of all standard MD simulations starting from the unfolding intermediates extracted from the SMD simulations.

| System name[a] | Origin of the starting structure | Simulation time (μs) × number of simulations |
|---|---|---|
| Spiral-3Q$^{1-6}$ run3 | SMD: 3Q$^{1-6}$ run3 | 1 × 2 |
| Spiral-3Q$^{1-7}$ run2 | SMD: 3Q$^{1-7}$ run2 | |
| Spiral-3Q$^{1-8}$ run2 | SMD: 3Q$^{1-8}$ run2 | |
| Spiral-4Q$^{1-7}$ run3 | SMD: 4Q$^{1-7}$ run3 | |
| Spiral-4Q$^{1-8}$ run1 | SMD: 4Q$^{1-8}$ run1 | |
| Spiral-4Q$^{1-8}$ run2 | SMD: 4Q$^{1-8}$ run2 | |
| Spiral-3Q$^{1-6}$ run3-spce | SMD: 3Q$^{1-6}$ run3 | |
| Spiral-3Q$^{1-7}$ run2-spce | SMD: 3Q$^{1-7}$ run2 | |
| Spiral-3Q$^{1-8}$ run2-spce | SMD: 3Q$^{1-8}$ run2 | |
| Spiral-4Q$^{1-7}$ run3-spce | SMD: 4Q$^{1-7}$ run3 | |
| Spiral-4Q$^{1-8}$ run1-spce | SMD: 4Q$^{1-8}$ run1 | |
| Spiral-4Q$^{1-8}$ run2-spce | SMD: 4Q$^{1-8}$ run2 | |
| Shifted-3Q$^{1-8}$ run2 | SMD: 3Q$^{1-8}$ run2 | |
| Shifted-3Q$^{1-8}$ run3 | SMD: 3Q$^{1-8}$ run3 | |
| Shifted-4Q$^{1-6}$ run3 | SMD: 4Q$^{1-6}$ run3 | |
| Rotated-4Q$^{1-8}$ run1 | SMD: 4Q$^{1-8}$ run1 | |
| 3Q-GQ | Starting 3Q-GQ structure | 1 × 1 |
| 4Q-GQ | Starting 4Q-GQ structure | |
| 5Q-GQ | Starting 5Q-GQ structure | |
| 3Q-GQ-spce | Starting 3Q-GQ structure | |
| 4Q-GQ-spce | Starting 4Q-GQ structure | |

[a] "Spiral" and "Shifted" denote the intermediate structure types, spiral G4s conformation and G4s with G-tract slippage, respectively, which were used as the starting structures of standard MD. The number of Qs and spring direction (superscript) indicate the original SMD system of the starting structure, followed by the number specifying one of the three runs in the SMD system. The systems with SPC/E water model are labelled with "-spce", and OPC water model was used otherwise.

**Table S3:** Statistics of unfolding events in each system.

| OPC water | | | | | | |
|---|---|---|---|---|---|---|
| Number of quartets | Pulling anchors | Unzipping | Strand slippage | Partial spiral GQ | Opening | Rotation (cross-like structure) |
| 3 | 1-6 | 6 | 1 | 1 | 0 | 0 |
| 3 | 1-7 | 6 | 0 | 1 | 0 | 0 |
| 3 | 1-8 | 5 | 4 | 3 | 2 | 2 |
| 4 | 1-6 | 11 | 2 | 1 | 0 | 0 |
| 4 | 1-7 | 11 | 1 | 1 | 0 | 0 |
| 4 | 1-8 | 7 | 1 | 3 | 2 | 3 |
| 5 | 1-6 | 15 | 1 | 0 | 3 | 2 |
| 5 | 1-7 | 16 | 1 | 0 | 0 | 0 |
| 5 | 1-8 | 14 | 3 | 1 | 1 | 1 |
| SPC/E water | | | | | | |
| Number of quartets | Pulling anchors | Unzipping | Strand slippage | Partial spiral GQ | Opening | Rotation (cross-like structure) |
| 3 | 1-6 | 7 | 5 | 2 | 2 | 1 |
| 3 | 1-7 | 7 | 1 | 1 | 2 | 1 |
| 3 | 1-8 | 5 | 2 | 1 | 2 | 2 |
| 4 | 1-6 | 11 | 4 | 2 | 1 | 0 |
| 4 | 1-7 | 9 | 1 | 1 | 0 | 0 |
| 4 | 1-8 | 7 | 1 | 3 | 2 | 3 |

**Table S4:** The overview of all unfolding events in SMD simulations with OPC water model.

| System name | Runs | Unfolding transitions [a] (time in ns) |
|---|---|---|
| $3Q^{1-6}$ | 1 | 6-end unzipping (637); 6-end unzipping (702); 2-6 G-tract detachment (717) |
| | 2 | 6-end unzipping (663); 6-end unzipping (690); 2-6 G-tract detachment (694) |
| | 3 | 6-end unzipping (610); 6-end unzipping (697); 6-end slippage (699); L-partial spiral[b] (725); 2-6 G-tract detachment (728) |
| $3Q^{1-7}$ | 1 | 7-end unzipping (788); 7-end unzipping (845); 3-7 G-tract detachment (850) |
| | 2 | 7-end unzipping (869); R-partial spiral (879); 7-end unzipping (884); 3-7 G-tract detachment (894) |
| | 3 | 7-end unzipping (816); 7-end unzipping (852); 3-7 G-tract detachment (855) |
| $3Q^{1-8}$ | 1 | 8-end unzipping (742); 8-end unzipping (751); 4-8 G-tract detachment (768) |
| | 2 | 8-end unzipping (733); R-partial spiral (741); 1-end slippage (758); R-partial spiral (801); 5-end slippage & 4-8 G-tract detachment (813) |
| | 3 | 8-end unzipping (735); R-partial spiral (774); 1-5,4-8 G-tracts opening & rotation into cross-like GQ (775); 1-end slippage (808); 1-5,4-8 G-tracts opening & rotation into cross-like GQ (879); 1-end slippage (882); 8-end unzipping (886); 1-5 G-tract detachment (888) |
| $4Q^{1-6}$ | 1 | 6-end unzipping (899); 6-end unzipping (1001); 6-end unzipping (1008); 2-6 G-tract detachment (1011) |
| | 2 | 6-end unzipping (740); 6-end unzipping (1110); 6-end unzipping & 1-end unzipping (1112); 6-end slippage (1117); 2-6 strand detachment (1121) |
| | 3 | 6-end unzipping (869); 6-end unzipping (964); 1-end unzipping (973); L-partial spiral (987); 6-end slippage (993); 6-end unzipping (1022); 2-6 G-tract detachment (1056) |
| $4Q^{1-7}$ | 1 | 7-end unzipping (857); 7-end unzipping (877); 7-end unzipping (960); 3-7 G-tract detachment (964) |
| | 2 | 1-end unzipping (896); 1-end unzipping (901); 1-end unzipping (912); 1-5 G-tract detachment (927) |
| | 3 | 1-end unzipping (732); 7-end unzipping (886); 1-end unzipping (916); R-partial spiral (994); 7,8-end slippage (1005); 7-end unzipping (1050); 7-end unzipping (1149); 3-7 G-tract detachment (1219) |
| $4Q^{1-8}$ | 1 | 8-end unzipping (646); R-partial spiral & 3-7,4-8 G-tracts rotation into cross-like GQ (784); 1-5,4-8 G-tracts opening (805); 1,2-end slippage (807); 1-end unzipping × 2 (907); 4-8 G-tract detachment (916) |
| | 2 | 8-end unzipping (727); 8-end unzipping (783); R-spiral (816); 1-5,4-8 G-tracts opening and rotation into cross-like GQ (819); 4-8 G-tract detachment (868) |
| | 3 | 8-end unzipping (709); R-partial spiral & 1-5,4-8 G-tracts rotation into cross-like GQ (792); 8-end unzipping (798); 4-8 G-tract detachment (933) |
| $5Q^{1-6}$ | 1 | 6-end unzipping (716); 1-end unzipping (903); 6-end unzipping (976); 6-end unzipping & 1-5,2-6 G-tracts opening (1130); 1-5 G-tract rotation into cross-like GQ & 1-end unzipping (1132); 2-6 G-tract detachment (1134) |
| | 2 | 1-end unzipping (696); 1-end unzipping (817); 6-end unzipping (935); 6-end unzipping & 2-6,3-7 G-tracts opening & 6-end unzipping (974); 2-6 G-tract detachment (981) |
| | 3 | 1-end unzipping (644); 1-end unzipping (898); 6-end unzipping (909); 6-end unzipping & 1-5,2-6 G-tracts opening and rotation into cross-like GQ & 6-end unzipping (974); 6-end slippage (978); 2-6 G-tract detachment (1005) |
| $5Q^{1-7}$ | 1 | 7-end unzipping (628); 1-end unzipping (792); 7-end unzipping (884); 7-end unzipping (925); 7-end unzipping (961); 3-7 G-tract detachment (978) |
| | 2 | 7-end unzipping (696); 7-end unzipping (721); 7-end unzipping (916); 1-end unzipping (1048); 7-end unzipping & 7-end slippage (1059); 1-end unzipping (1199); 3-7 G-tract detachment (1206) |
| | 3 | 1-end unzipping (682); 1-end unzipping & 2-end unzipping (782); 7-end unzipping (853); 1-end unzipping (961); 1-5 G-tract detachment (1015) |
| $5Q^{1-8}$ | 1 | 8-end unzipping (691); 8-end unzipping (871); one 8-end-G slide in 3'-direction (929); 1-5,4-8 G-tracts rotation into cross-like GQ (930); 1-end unzipping × 3 (933); 4-8 G-tract detachment (937) |
| | 2 | 8-end unzipping (812); 8-end unzipping (825); 1-end unzipping (852); 1-5,4-8 G-tracts opening (862); 8-end unzipping (868), 8-end slippage & attachment to the side of 3-7 (873); 4-8 G-tract detachment (883) |
| | 3 | 8-end unzipping (680); 7-end unzipping (802); 8-end unzipping (808); 7,8-end-G-bases intercalation (since 880); 8-end unzipping (885); 8-end unzipping & R-partial spiral (902); 8-end slippage & 7-end rotation into partial cross-like GQ (919); 7,8-end slippage (925) and subsequently distorted with huge conformational changes |

[a]The unfolding transitions listed in the table are summarized in the main text (see Figure 3). The specific transition is labelled by terminus number or terminus numbers of the strand(s) where the transition occurred. For example, “7-end unzipping” means the unzipping happened on the G on the

terminus 7, and “1-5,2-6 G-tracts opening” refers the opening between 1-5 and 2-6 G-tracts. In cases we observed two simultaneous transitions, we use “&”.

[b]The partial spiral conformations are classified into left-handiness and right-handiness, labelled by “L-partial spiral” and “R-partial spiral”, respectively.

**Table S5:** Overview of all unfolding events in SMD simulations with SPC/E water model.

| System name | Runs | Unfolding transitions[a] (time in ns) |
|---|---|---|
| $3Q^{1\text{-}6}$-spce | 1 | 6-end unzipping (686); 6-end unzipping (695); 1-5,2-6 G-tracts opening (698); 6-end slippage (699); 2-end slippage & 1-5 G-tract rotation into cross-like GQ (700); 1-5 G-tract rotated off from GQ core (703); 2-6 G-tract formed bases intercalation (709); 1-5 G-tract detachment (711) |
| | 2 | 6-end unzipping (736); 6-end unzipping & L-partial spiral & 6-end slippage (739); 1-end unzipping (742); L-partial spiral (747); 6-end slippage (748); 1-5,2-6 G-tracts opening (848); 1-end slippage (850); 1-5 G-tract detachment (852) |
| | 3 | 6-end unzipping (689); 6-end unzipping (705); 2-6 G-tract detachment (709) |
| $3Q^{1\text{-}7}$-spce | 1 | 1-end unzipping (830); 1-5,4-8 G-tracts opening and 1-5 G-tract rotation into cross-like GQ (848); 1,2-end slippage & 1-end unzipping & 1-5,4-8 G-tracts closing (859); 7-end unzipping & L-partial spiral (863); no L-partial spiral (876); 1-5,4-8 G-tracts opening (927); 1-5,4-8 G-tracts closing (942); 1-5 G-tract detachment (981) |
| | 2 | 7-end unzipping (849); 7-end unzipping (870); 3-7 G-tract detachment (941) |
| | 3 | 7-end unzipping (775); 7-end unzipping (878); 3-7 G-tract detachment (893) |
| $3Q^{1\text{-}8}$-spce | 1 | 8-end slippage (781); 1-5,4-8 G-tracts opening & 3-7,4-8 G-tracts rotation into cross-like GQ (812); 8-end unzipping (894); 4-8 G-tract detachment (897) |
| | 2 | 8-end unzipping (757); R-partial spiral & 1-5,4-8 opening & 8-end unzipping & 8-end slippage (817); 3-7,4-8 G-tracts detachment (860) |
| | 3 | 8-end unzipping and then refolding (685-712); 8-end unzipping (760); 3-7,4-8 G-tracts rotation into cross-like GQ (828); 8-end unzipping (885); 4-8 G-tract detachment (887) |
| $4Q^{1\text{-}6}$-spce | 1 | 1-end unzipping (803); 1-end unzipping (998); L-partial spiral & 6,7-end slippage (1021); 6-end unzipping (1026); 6,7-end slippage (1151); 1-5,2-6 G-tracts opening (1157); 1-5 G-tract detachment (1191) |
| | 2 | 1-end unzipping and then refolding (711-718); 6-end unzipping (864); 1-end unzipping (985); L-partial spiral & 1-end unzipping & 6-end unzipping (1173); 6-end unzipping & 1-end slippage & 6-end slippage (1175); 2-6 G-tract detachment (1186) |
| | 3 | 6-end unzipping (925); 6-end unzipping (1114); 6-end unzipping (1119); 2-6 G-tract detachment (1124) |
| $4Q^{1\text{-}7}$-spce | 1 | 1-end unzipping (788); 1-end unzipping (867); 1-end unzipping (920); 1-5 G-tract detachment (1065) |
| | 2 | 7-end unzipping (725); 7-end unzipping (928); 7-end unzipping (989); 7-end slippage (1012); L-partial spiral (1018); 3-7 G-tract detachment (1045) |
| | 3 | 7-end unzipping (791); 7-end unzipping (864); 7-end unzipping (993); 3-7 G-tract detachment (1170) |
| $4Q^{1\text{-}8}$-spce | 1 | 8-end unzipping (764); 8-end unzipping (782); 1-5,4-8 G-tracts opening & rotation into cross-like GQ (795); 1-end unzipping (825); 4-8 G-tract detachment (845) |
| | 2 | 1-end unzipping (815); R-partial spiral & 8-end unzipping (854); 1-5,2-6 G-tracts opening & 1-5,4-8 G-tracts rotation into cross-like GQ (864); 8-end slippage (886); complicated unfolding with intercalation of bases in strands 1,2, 1Q remaining at the end of the simulation |
| | 3 | 8-end unzipping and then refolding (618-631); 8-end unzipping (665); unzipped 8-end G intercalated with 7-end T (~677); 7-end G rotation formed partial cross-like structure (777); 4-8 G-tract rotation to cross-like GQ and soon rotated back & 5,8-end partial opening (828); 4-8 G-tract rotation to cross-like GQ & 1-5,4-8 G-tracts opening (838); 8-end unzipping × 2 & the 5′-Q refolding (841); 1-5,4-8 G-tracts opening (846); 4-8 G-tract detachment (850) |

[a]See the note in Table S4.

**Table S6:** The values of quartet-step rise distances right before the unfolding period during SMD simulations with OPC water, with corresponding data from standard MD as reference.

| System name | Runs | Q1-Q2 (5' side, Å) | Q2-Q3 (Å) | Q3-Q4 (Å) | Q4-Q5 (Å) |
|---|---|---|---|---|---|
| $3Q^{1-6}$ | 1 | 3.394 | 3.460 | - | - |
| | 2 | 3.408 | 3.484 | - | - |
| | 3 | 3.403 | 3.450 | - | - |
| $3Q^{1-7}$ | 1 | 3.451 | 3.582 | - | - |
| | 2 | 3.503 | 3.603 | - | - |
| | 3 | 3.508 | 3.617 | - | - |
| $3Q^{1-8}$ | 1 | 3.548 | 3.738 | - | - |
| | 2 | 3.646 | 3.649 | - | - |
| | 3 | 3.567 | 3.668 | - | - |
| 3Q standard MD | 1 | 3.365 | 3.494 | - | - |
| $4Q^{1-6}$ | 1 | 3.382 | 3.438 | 3.478 | - |
| | 2 | 3.374 | 3.441 | 3.522 | - |
| | 3 | 3.362 | 3.446 | 3.421 | - |
| $4Q^{1-7}$ | 1 | 3.449 | 3.503 | 3.506 | - |
| | 2 | 3.533 | 3.639 | 3.772 | - |
| | 3 | 3.406 | 3.450 | 3.598 | - |
| $4Q^{1-8}$ | 1 | 3.490 | 3.597 | 3.636 | - |
| | 2 | 3.485 | 3.564 | 3.787 | - |
| | 3 | 3.590 | 3.677 | 3.717 | - |
| 4Q standard MD | 1 | 3.333 | 3.418 | 3.451 | - |
| $5Q^{1-6}$ | 1 | 3.382 | 3.365 | 3.480 | 3.667 |
| | 2 | 3.599 | 3.357 | 3.447 | 3.517 |
| | 3 | 3.369 | 3.342 | 3.429 | 3.480 |
| $5Q^{1-7}$ | 1 | 3.349 | 3.420 | 3.514 | 3.672 |
| | 2 | 3.415 | 3.398 | 3.494 | 3.568 |
| | 3 | 3.357 | 3.358 | 3.437 | 3.528 |
| $5Q^{1-8}$ | 1 | 3.574 | 3.520 | 3.708 | 3.673 |
| | 2 | 3.608 | 3.619 | 3.808 | 3.798 |
| | 3 | 3.518 | 3.570 | 3.693 | 3.779 |
| 5Q standard MD | 1 | 3.339 | 3.349 | 3.400 | 3.411 |

**Table S7:** The values of planarity right before the unfolding period during SMD simulations with OPC water, with corresponding data from standard MD as reference.

| System name | Runs | Q1 (5' side, Å) | Q2 (Å) | Q3 (Å) | Q4 (Å) | Q5 (Å) |
|---|---|---|---|---|---|---|
| $3Q^{1-6}$ | 1 | 0.293 | 0.289 | 0.357 | - | - |
| | 2 | 0.308 | 0.299 | 0.355 | - | - |
| | 3 | 0.309 | 0.294 | 0.356 | - | - |
| $3Q^{1-7}$ | 1 | 0.355 | 0.369 | 0.4845 | - | - |
| | 2 | 0.451 | 0.408 | 0.487 | - | - |
| | 3 | 0.454 | 0.422 | 0.500 | - | - |
| $3Q^{1-8}$ | 1 | 0.481 | 0.468 | 0.789 | - | - |
| | 2 | 0.602 | 0.444 | 0.725 | - | - |
| | 3 | 0.489 | 0.516 | 0.770 | - | - |
| 3Q standard MD | 1 | 0.325 | 0.321 | 0.405 | - | - |
| $4Q^{1-6}$ | 1 | 0.287 | 0.225 | 0.262 | 0.443 | - |
| | 2 | 0.302 | 0.233 | 0.296 | 0.558 | - |
| | 3 | 0.278 | 0.213 | 0.268 | 0.377 | - |
| $4Q^{1-7}$ | 1 | 0.353 | 0.246 | 0.311 | 0.545 | - |
| | 2 | 0.557 | 0.460 | 0.568 | 0.661 | - |
| | 3 | 0.450 | 0.361 | 0.409 | 0.526 | - |
| $4Q^{1-8}$ | 1 | 0.522 | 0.378 | 0.434 | 0.573 | - |
| | 2 | 0.680 | 0.610 | 0.676 | 0.902 | - |
| | 3 | 0.690 | 0.559 | 0.569 | 0.690 | - |
| 4Q standard MD | 1 | 0.236 | 0.181 | 0.208 | 0.258 | - |
| $5Q^{1-6}$ | 1 | 0.319 | 0.255 | 0.251 | 0.333 | 0.683 |
| | 2 | 0.651 | 0.222 | 0.221 | 0.274 | 0.428 |
| | 3 | 0.389 | 0.202 | 0.179 | 0.221 | 0.336 |
| $5Q^{1-7}$ | 1 | 0.254 | 0.196 | 0.209 | 0.341 | 0.645 |
| | 2 | 0.351 | 0.325 | 0.321 | 0.376 | 0.620 |
| | 3 | 0.462 | 0.254 | 0.192 | 0.228 | 0.417 |
| $5Q^{1-8}$ | 1 | 0.696 | 0.538 | 0.518 | 0.539 | 0.868 |
| | 2 | 0.725 | 0.534 | 0.523 | 0.711 | 1.001 |
| | 3 | 0.550 | 0.403 | 0.398 | 0.537 | 0.755 |
| 5Q standard MD | 1 | 0.230 | 0.185 | 0.167 | 0.185 | 0.260 |

**Table S8:** The values of quartet-step tilt angles right before the unfolding period during SMD simulations with OPC water, with corresponding data from standard MD as reference.

| System name | Runs | Q1-Q2 (5' side, °) | Q2-Q3 (°) | Q3-Q4 (°) | Q4-Q5 (°) |
|---|---|---|---|---|---|
| $3Q^{1-6}$ | 1 | 1.546 | 1.773 | - | - |
| | 2 | 1.607 | 1.681 | - | - |
| | 3 | 1.526 | 1.710 | - | - |
| $3Q^{1-7}$ | 1 | 2.711 | 2.444 | - | - |
| | 2 | 4.006 | 3.606 | - | - |
| | 3 | 3.748 | 3.145 | - | - |
| $3Q^{1-8}$ | 1 | 4.286 | 6.868 | - | - |
| | 2 | 7.030 | 4.530 | - | - |
| | 3 | 4.726 | 5.014 | - | - |
| 3Q standard MD | 1 | 1.421 | 1.515 | - | - |
| $4Q^{1-6}$ | 1 | 1.415 | 1.972 | 4.076 | - |
| | 2 | 1.345 | 1.743 | 2.904 | - |
| | 3 | 1.354 | 1.748 | 3.461 | - |
| $4Q^{1-7}$ | 1 | 2.124 | 3.081 | 7.675 | - |
| | 2 | 4.891 | 4.134 | 4.345 | - |
| | 3 | 5.913 | 2.179 | 2.215 | - |
| $4Q^{1-8}$ | 1 | 3.679 | 4.435 | 4.880 | - |
| | 2 | 7.950 | 4.468 | 6.143 | - |
| | 3 | 4.962 | 6.220 | 5.465 | - |
| 4Q standard MD | 1 | 1.439 | 1.446 | 1.494 | - |
| $5Q^{1-6}$ | 1 | 4.011 | 1.451 | 2.224 | 5.651 |
| | 2 | 8.507 | 1.484 | 1.587 | 2.448 |
| | 3 | 4.591 | 1.436 | 1.371 | 1.519 |
| $5Q^{1-7}$ | 1 | 1.248 | 1.492 | 2.328 | 5.861 |
| | 2 | 4.554 | 1.919 | 1.795 | 4.577 |
| | 3 | 3.566 | 1.714 | 1.426 | 3.049 |
| $5Q^{1-8}$ | 1 | 7.209 | 4.863 | 7.345 | 9.243 |
| | 2 | 6.137 | 6.343 | 7.977 | 9.483 |
| | 3 | 4.073 | 4.912 | 6.310 | 6.371 |
| 5Q standard MD | 1 | 1.363 | 1.220 | 1.264 | 1.414 |

**Table S9:** Overview of all transitions in standard unbiased MD simulations listed in Table S2, starting from the intermediates extracted from SMD simulations.

| System name | Runs | Transition events[a] |
|---|---|---|
| Spiral-3Q$^{1\text{-}6}$-r3 | 1 | 1-end slippage (6.5); L-spiral (17.9); 1-end slippage (20.5); 1,4-end slippage & 1-5,2-6 G-tracts opening (24.6); 1-5 G-tract detachment (33.3); 2-6 G-tract detachment (95.0); Ended with 2 detached strands and one duplex |
| | 2 | 6-end slippage (8.6); 2-end slippage (10.0); L-spiral (14.8); 6-end two G-bases refolding (17.4); 6,7-end slippage (17.7); L-spiral (21.3); bottom Q refolded (39.0); L-spiral finished, refolding completed (40.2) |
| Spiral-3Q$^{1\text{-}7}$-r2 | 1 | 8-end slippage (0.1); 7-end G-base refolding, refolding completed (211) |
| | 2 | 3-end slippage (0.8); 7-end G-base refolding, refolding completed (6.5) |
| Spiral-3Q$^{1\text{-}8}$-r2 | 1 | 5-end slippage (0.8); 8-end G-base refolding, refolding completed (50.0) |
| | 2 | 5-end slippage (0.8); 7-end unzipping (444); remained unfolded |
| Spiral-4Q$^{1\text{-}7}$-r3 | 1 | 4-end slippage (1.0); 7-end G-base refolding (8.1); 1-end G-base refolding (2nd G to the 1st Q, 15.0); L-spiral (only on the first two Qs, 27.4); 5-end slippage, refolding completed (33.7) |
| | 2 | 4-end slippage (15.7); R-spiral (22.4); 4-end slippage (49.0); remained unfolded |
| Spiral-4Q$^{1\text{-}8}$-r1 | 1 | 8-end slippage (143), remained unfolded |
| | 2 | 4-end slippage (15.0); 1-end G-base refolding, refolding completed (54.8) |
| Spiral-4Q$^{1\text{-}8}$-r2 | 1 | 5-end slippage (2.7); remained unfolded |
| | 2 | 5-end slippage (3.0); remained unfolded |
| Spiral-3Q$^{1\text{-}6}$-r3-spce | 1 | 5-end slippage (29.0); R-spiral (34.4); 1-end slippage (35.5); remained unfolded |
| | 2 | 5-end slippage (1.1); L-spiral (7.0); 5-end slippage (212); remained unfolded |
| Spiral-3Q$^{1\text{-}7}$-r2-spce | 1 | 3-end slippage (2.3); 7-end G-base refolding, refolding completed (119) |
| | 2 | 3-end slippage (0.5); 7-end G-base refolding, refolding completed (72.0) |
| Spiral-3Q$^{1\text{-}8}$-r2-spce | 1 | 1-end slippage (1.2); remained unfolded |
| | 2 | 5-end slippage (3.2); 1-5,4-8 G-tract opening (184); 1-end slippage (186); 5-end slippage, 1-5,4-8 G-tracts closing (198); remained unfolded |
| Spiral-4Q$^{1\text{-}7}$-r3-spce | 1 | 5-end slippage (5.7); R-spiral (6.2); 2-end slippage (198); remained unfolded |
| | 2 | 2-end slippage (27.9); remained unfolded |
| Spiral-4Q$^{1\text{-}8}$-r1-spce | 1 | 5-end slippage (15.5); 1-end G-base refolding (16.0); 8-end G-base refolding (but the bottom Q is not well-formed, 154); deformed GQ lasted till the end |
| | 2 | 5-end slippage (133); remained unfolded, 8-end unzipped G-base kept syn-glycosidic angle |
| Spiral-4Q$^{1\text{-}8}$-r2-spce | 1 | 5-end slippage (3.3); two 8-end G-bases refolding, refolding completed (25.8) |
| | 2 | 5-end slippage (2.2); two 8-end G-bases refolding, refolding completed (30.0) |
| Shifted-3Q$^{1\text{-}8}$-r2 | 1 | No transition detected |
| | 2 | No transition detected |
| Shifted-3Q$^{1\text{-}8}$-r3 | 1 | No transition detected |
| | 2 | No transition detected |
| Shifted-4Q$^{1\text{-}6}$-r3 | 1 | 6-end G-base refolding (126); remained unfolded |
| | 2 | 6-end G-base refolding (61.0); remained unfolded |
| Rotated-4Q$^{1\text{-}8}$-r1 | 1 | 3-7,4-8-G-tract rotating back & 3,4-end slippage (60.2); 1-end G-base refolding (68.0); 1-end G-base refolding (81.0); remained unfolded |
| | 2 | 3-7,4-8-G-tract rotating back (4.6); remained unfolded |

[a]See the note in Table S3. The "G-base refolding" event is the reverse action of unzipping. The standard MD simulation results did not always end with native or completely unfolded GQs, and thus, we noted down the ending of each run.

**Table S10:** The values of force out-of-plane angle right before the unfolding period.

| System name | Runs | Force out-of-plane angle (β, °) |
|---|---|---|
| $3Q^{1-6}$ | 1 | 27 |
| | 2 | 28 |
| | 3 | 27 |
| $3Q^{1-7}$ | 1 | 34 |
| | 2 | 34 |
| | 3 | 34 |
| $3Q^{1-8}$ | 1 | 56 |
| | 2 | 46 |
| | 3 | 57 |
| $4Q^{1-6}$ | 1 | 30 |
| | 2 | 29 |
| | 3 | 30 |
| $4Q^{1-7}$ | 1 | 39 |
| | 2 | 45 |
| | 3 | 43 |
| $4Q^{1-8}$ | 1 | 76 |
| | 2 | 65 |
| | 3 | 72 |
| $5Q^{1-6}$ | 1 | 34 |
| | 2 | 34 |
| | 3 | 36 |
| $5Q^{1-7}$ | 1 | 53 |
| | 2 | 50 |
| | 3 | 55 |
| $5Q^{1-8}$ | 1 | 77 |
| | 2 | 75 |
| | 3 | 81 |

**Table S11:** The values of quartet-step rise distances right before the unfolding period during SMD simulations with SPC/E water, with corresponding data from standard MD as reference.

| System name | Runs | Q1-Q2 (5' side, Å) | Q2-Q3 (Å) | Q3-Q4 (Å) |
|---|---|---|---|---|
| $3Q^{1\text{-}6}$-spce | 1 | 3.456 | 3.513 | - |
| | 2 | 3.367 | 3.510 | - |
| | 3 | 3.431 | 3.451 | - |
| $3Q^{1\text{-}7}$-spce | 1 | 3.561 | 3.609 | - |
| | 2 | 3.489 | 3.681 | - |
| | 3 | 3.479 | 3.767 | - |
| $3Q^{1\text{-}8}$-spce | 1 | 3.515 | 3.750 | - |
| | 2 | 3.511 | 3.700 | - |
| | 3 | 3.520 | 4.022 | - |
| 3Q standard MD | 1 | 3.384 | 3.462 | - |
| $4Q^{1\text{-}6}$-spce | 1 | 3.430 | 3.587 | 3.660 |
| | 2 | 3.356 | 3.509 | 3.588 |
| | 3 | 3.437 | 3.515 | 3.621 |
| $4Q^{1\text{-}7}$-spce | 1 | 3.459 | 3.612 | 3.614 |
| | 2 | 3.479 | 3.520 | 3.571 |
| | 3 | 3.469 | 3.580 | 3.666 |
| $4Q^{1\text{-}8}$-spce | 1 | 3.600 | 3.642 | 3.665 |
| | 2 | 3.943 | 3.620 | 3.733 |
| | 3 | 3.534 | 3.566 | 3.689 |
| 4Q standard MD | 1 | 3.342 | 3.411 | 3.435 |

**Table S12:** The values of planarity right before the unfolding period during SMD simulations with SPC/E water, with corresponding data from standard MD as reference.

| System name | Runs | Q1 (5' side, Å) | Q2 (Å) | Q3 (Å) | Q4 (Å) |
|---|---|---|---|---|---|
| $3Q^{1-6}$-spce | 1 | 0.300 | 0.326 | 0.422 | - |
| | 2 | 0.251 | 0.264 | 0.363 | - |
| | 3 | 0.296 | 0.309 | 0.433 | - |
| $3Q^{1-7}$-spce | 1 | 0.574 | 0.462 | 0.498 | - |
| | 2 | 0.445 | 0.460 | 0.540 | - |
| | 3 | 0.316 | 0.327 | 0.605 | - |
| $3Q^{1-8}$-spce | 1 | 0.622 | 0.602 | 0.773 | - |
| | 2 | 0.501 | 0.524 | 0.805 | - |
| | 3 | 0.565 | 0.574 | 1.155 | - |
| 3Q standard MD | 1 | 0.269 | 0.261 | 0.333 | - |
| $4Q^{1-6}$-spce | 1 | 0.308 | 0.245 | 0.402 | 0.517 |
| | 2 | 0.272 | 0.268 | 0.282 | 0.665 |
| | 3 | 0.321 | 0.294 | 0.350 | 0.688 |
| $4Q^{1-7}$-spce | 1 | 0.597 | 0.499 | 0.581 | 0.645 |
| | 2 | 0.293 | 0.198 | 0.274 | 0.645 |
| | 3 | 0.340 | 0.367 | 0.460 | 0.637 |
| $4Q^{1-8}$-spce | 1 | 1.137 | 0.721 | 0.886 | 1.067 |
| | 2 | 1.870 | 0.744 | 0.687 | 0.849 |
| | 3 | 0.443 | 0.411 | 0.539 | 0.947 |
| 4Q standard MD | 1 | 0.263 | 0.184 | 0.198 | 0.260 |

**Table S13:** The values of quartet-step tilt angles right before the unfolding period during SMD simulations with SPC/E water, with corresponding data from standard MD as reference.

| System name | Runs | Q1-Q2 (5' side, °) | Q2-Q3 (°) | Q3-Q4 (°) |
|---|---|---|---|---|
| $3Q^{1-6}$-spce | 1 | 1.959 | 2.371 | - |
| | 2 | 1.437 | 2.733 | - |
| | 3 | 1.700 | 2.210 | - |
| $3Q^{1-7}$-spce | 1 | 4.343 | 3.217 | - |
| | 2 | 3.602 | 3.045 | - |
| | 3 | 2.373 | 6.735 | - |
| $3Q^{1-8}$-spce | 1 | 5.133 | 7.543 | - |
| | 2 | 3.578 | 5.513 | - |
| | 3 | 3.505 | 11.873 | - |
| 3Q standard MD | 1 | 1.552 | 1.693 | - |
| $4Q^{1-6}$-spce | 1 | 1.959 | 1.029 | 1.415 |
| | 2 | 2.326 | 0.528 | 2.400 |
| | 3 | 2.089 | 1.676 | 4.899 |
| $4Q^{1-7}$-spce | 1 | 9.205 | 0.844 | 2.573 |
| | 2 | 0.945 | 0.128 | 4.089 |
| | 3 | 2.844 | 2.723 | 4.182 |
| $4Q^{1-8}$-spce | 1 | 9.345 | 4.828 | 5.095 |
| | 2 | 4.253 | 5.956 | 9.532 |
| | 3 | 4.179 | 3.836 | 6.542 |
| 4Q standard MD | 1 | 1.373 | 1.362 | 1.557 |

**Supplementary figures:**

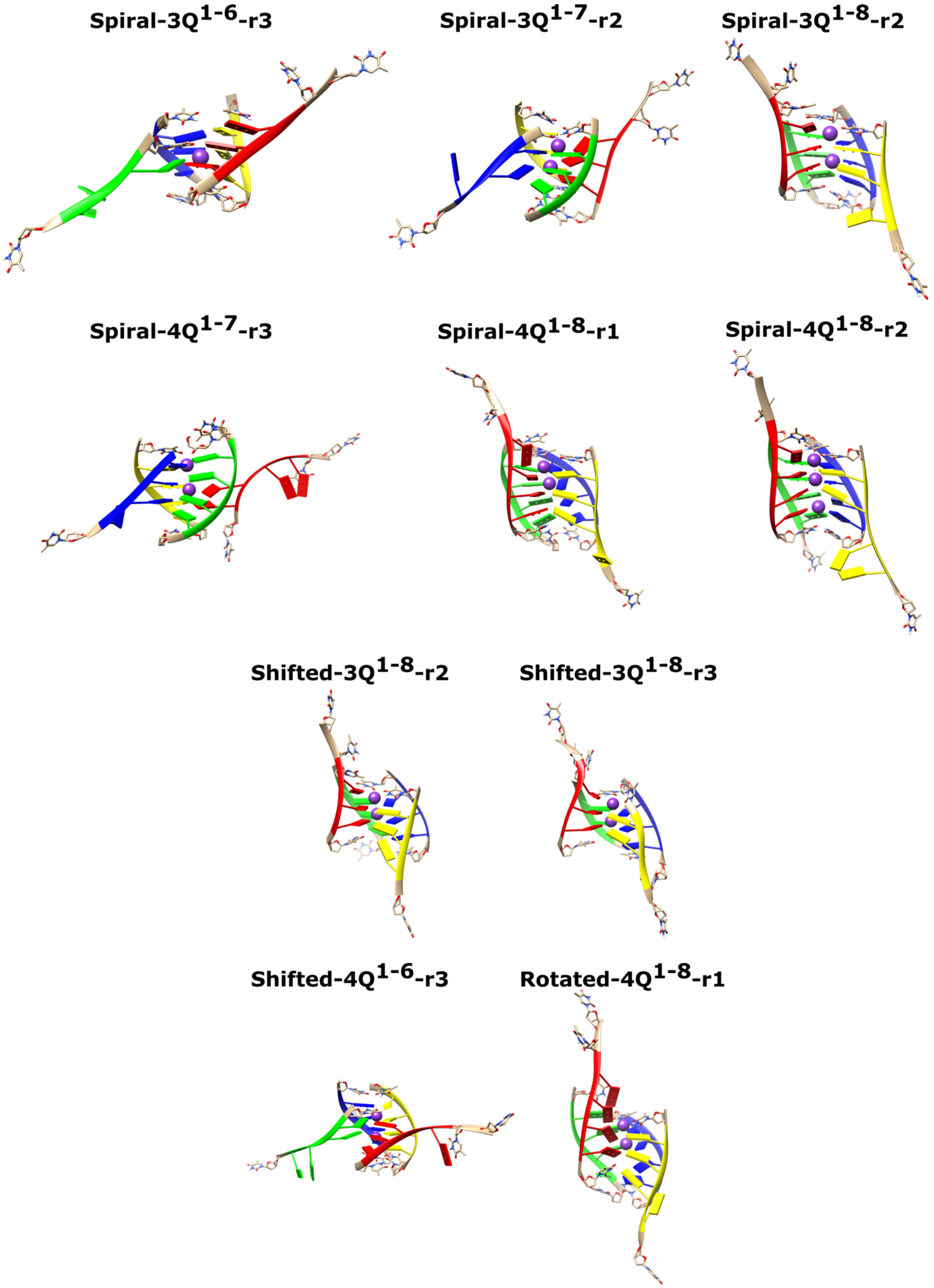


**Figure S1. Starting structures of the unbiased MD simulations, extracted from the SMD simulations.** The naming and the structural origin of each system are detailed in Table S2.

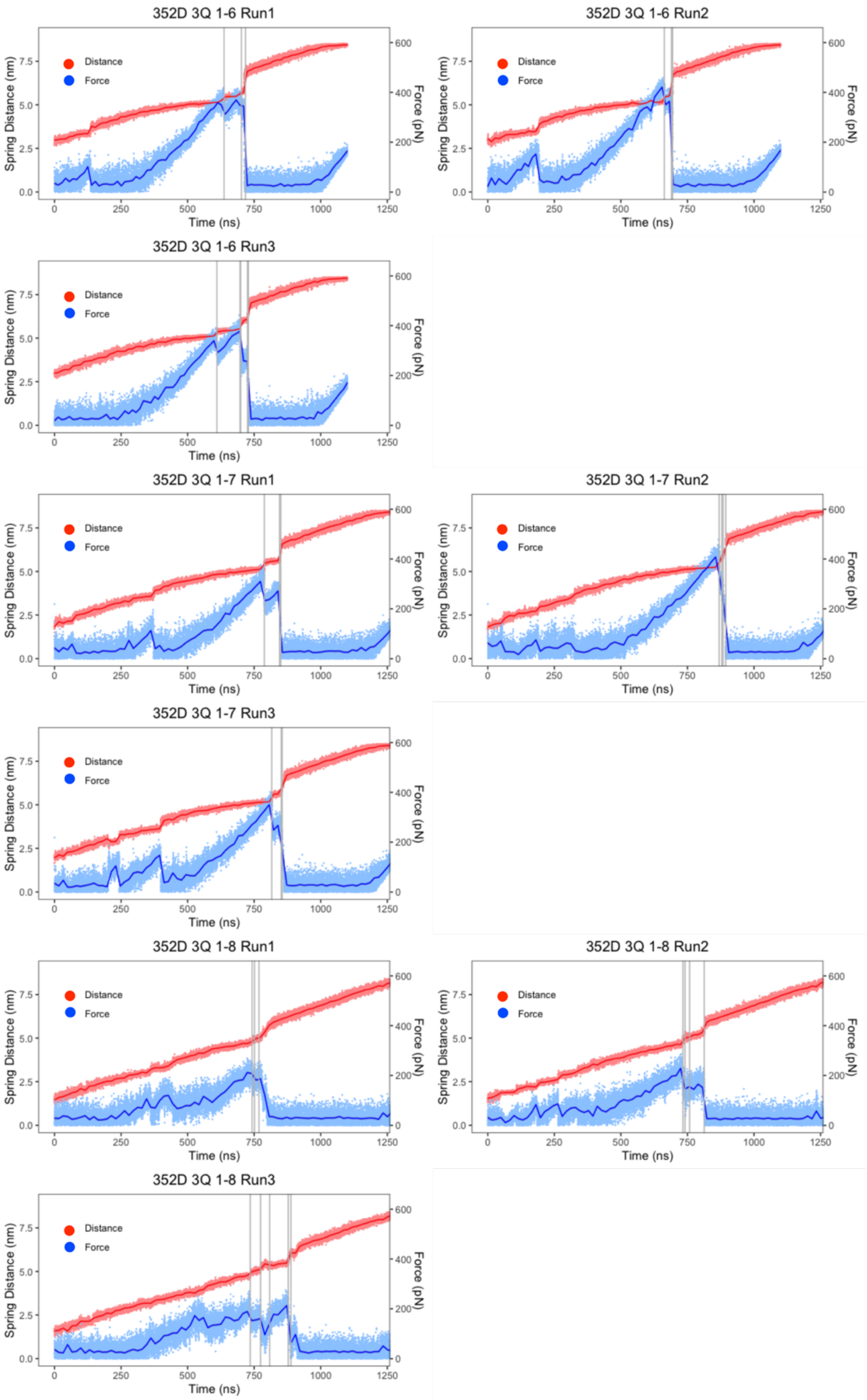


**Figure S2.** Evolution of force and spring anchors distance of 3Q-GQs during pulling. All transition events are labelled as grey vertical lines, and the time-averaged distance and force are in solid lines. Note that the steep increase in force near the very end of 1-6 and 1-7 pulling simulations is caused by physical size limits of the periodic boundary conditions and is meaningless and not affecting the previous unfolding.

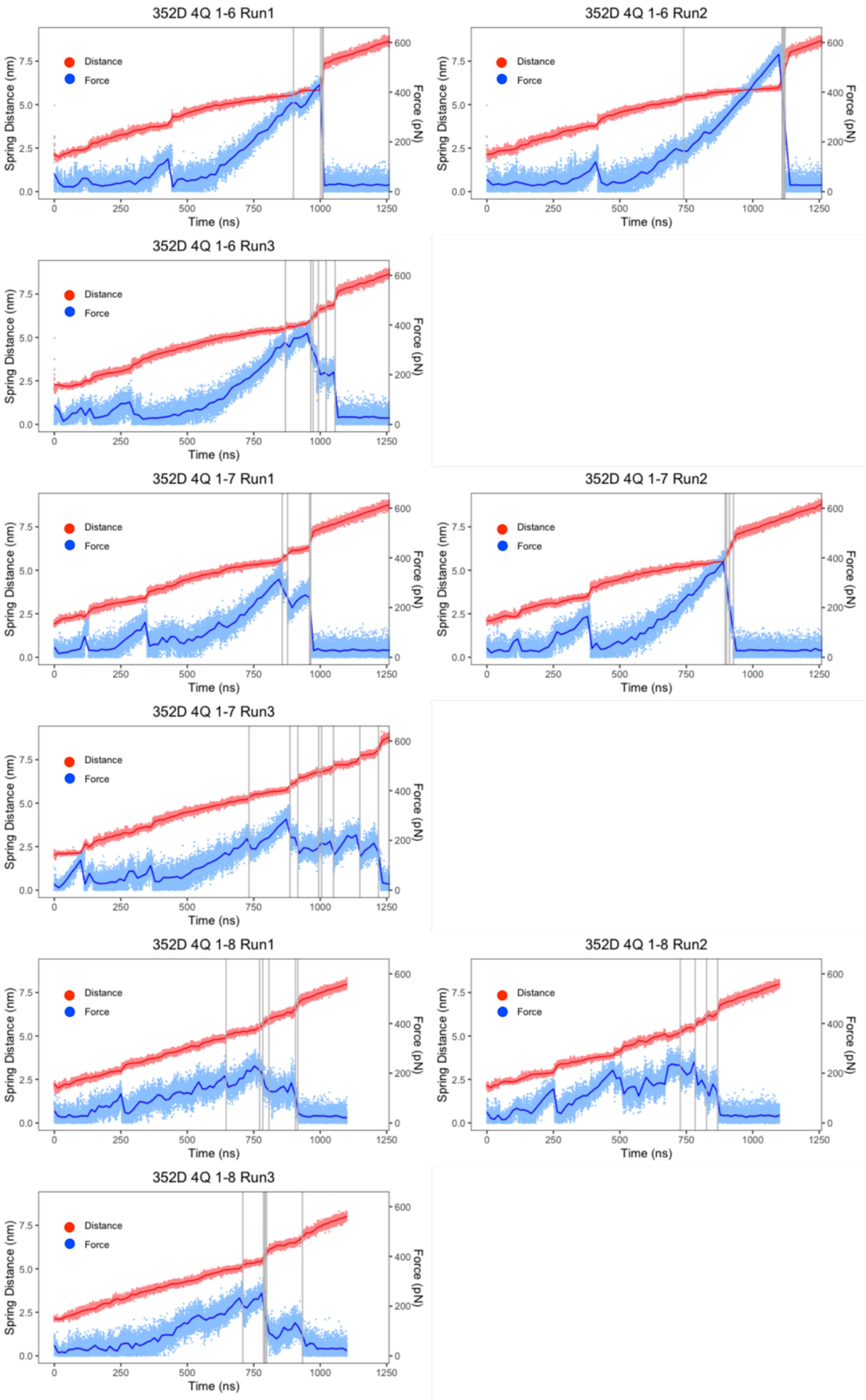


**Figure S3.** Evolution of force and spring anchors distance of 4Q-GQs during pulling. All transition events are labelled as grey vertical lines, and the time-averaged distance and force are in solid lines.

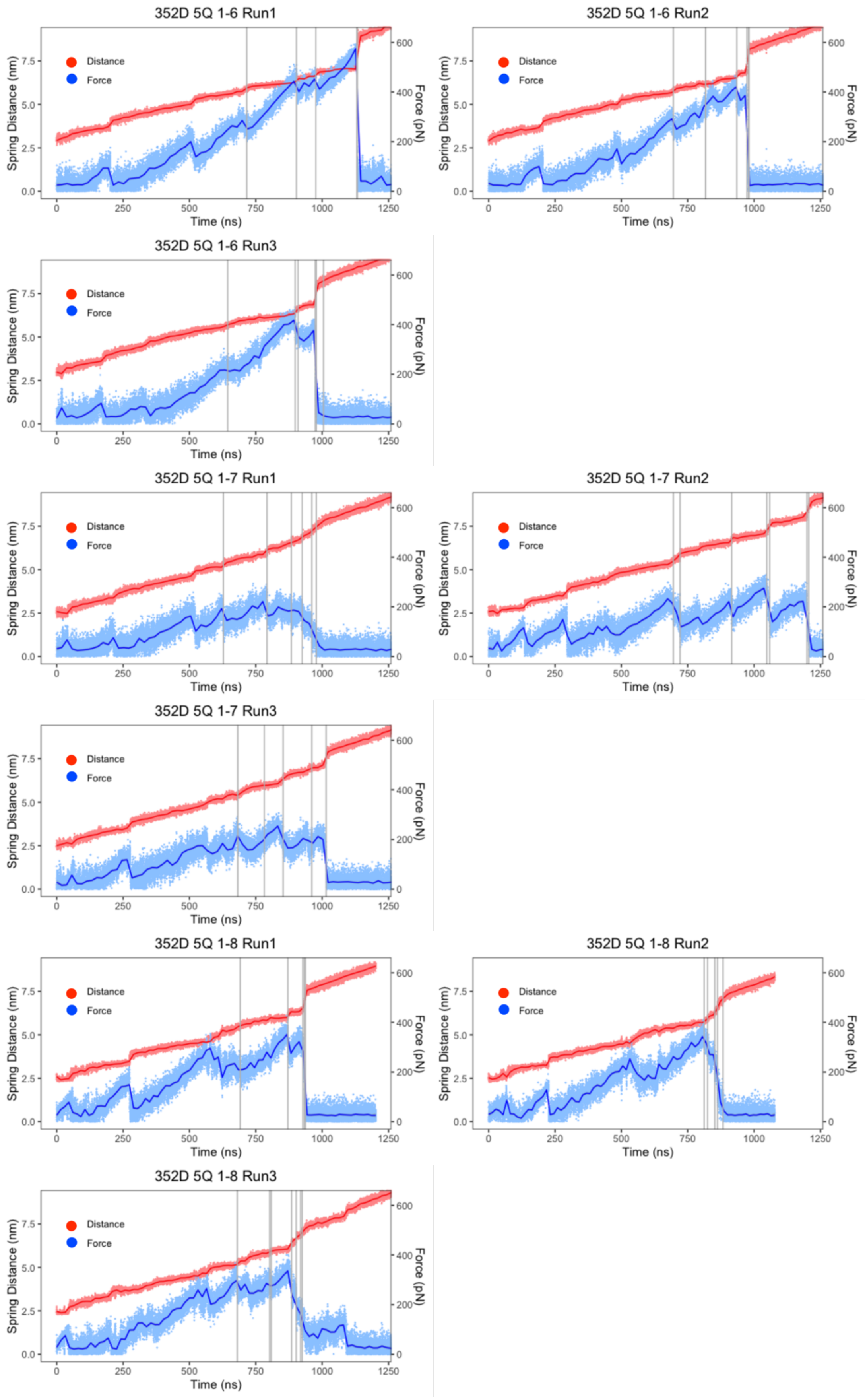


**Figure S4.** Evolution of force and spring anchors distance of 5Q-GQs during pulling. All transition events are labelled as grey vertical lines, and the time-averaged distance and force are in solid lines.

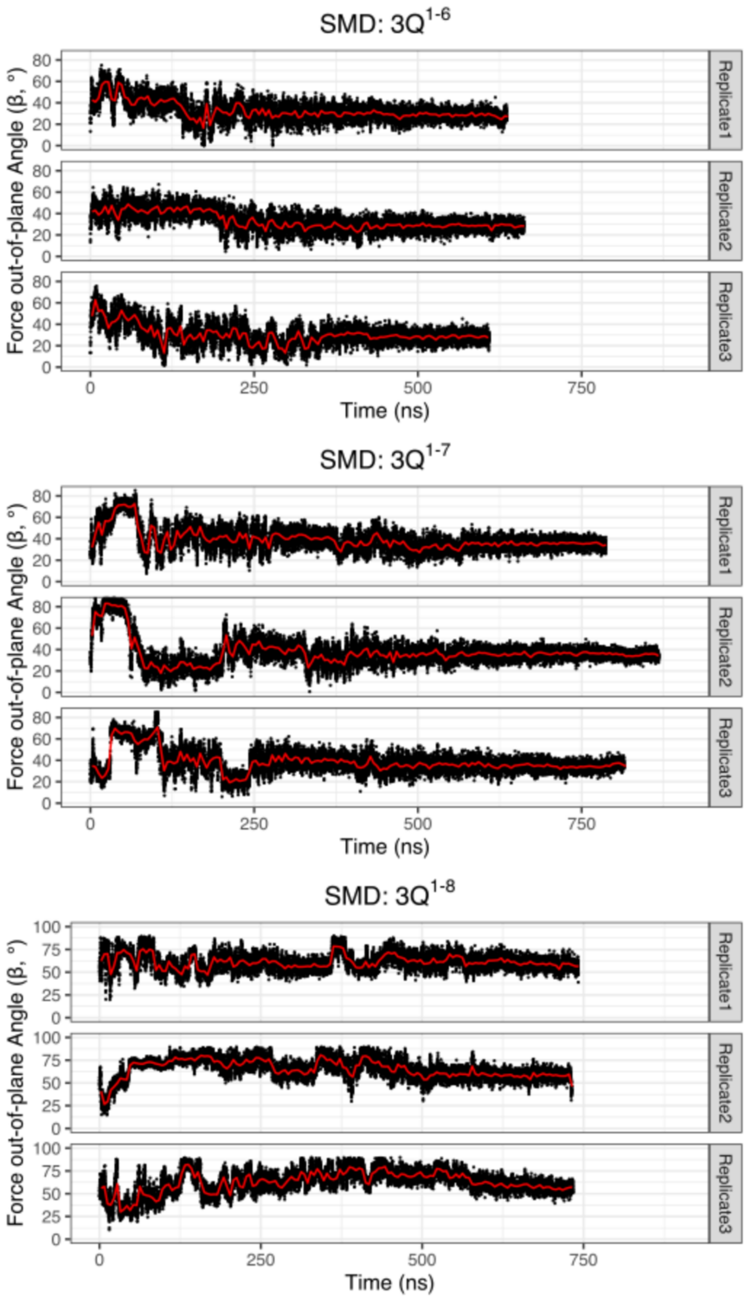


**Figure S5.** Evolution of force out-of-plane angle of 3Q-GQs during pulling, and the red solid lines show the time-averaged values.

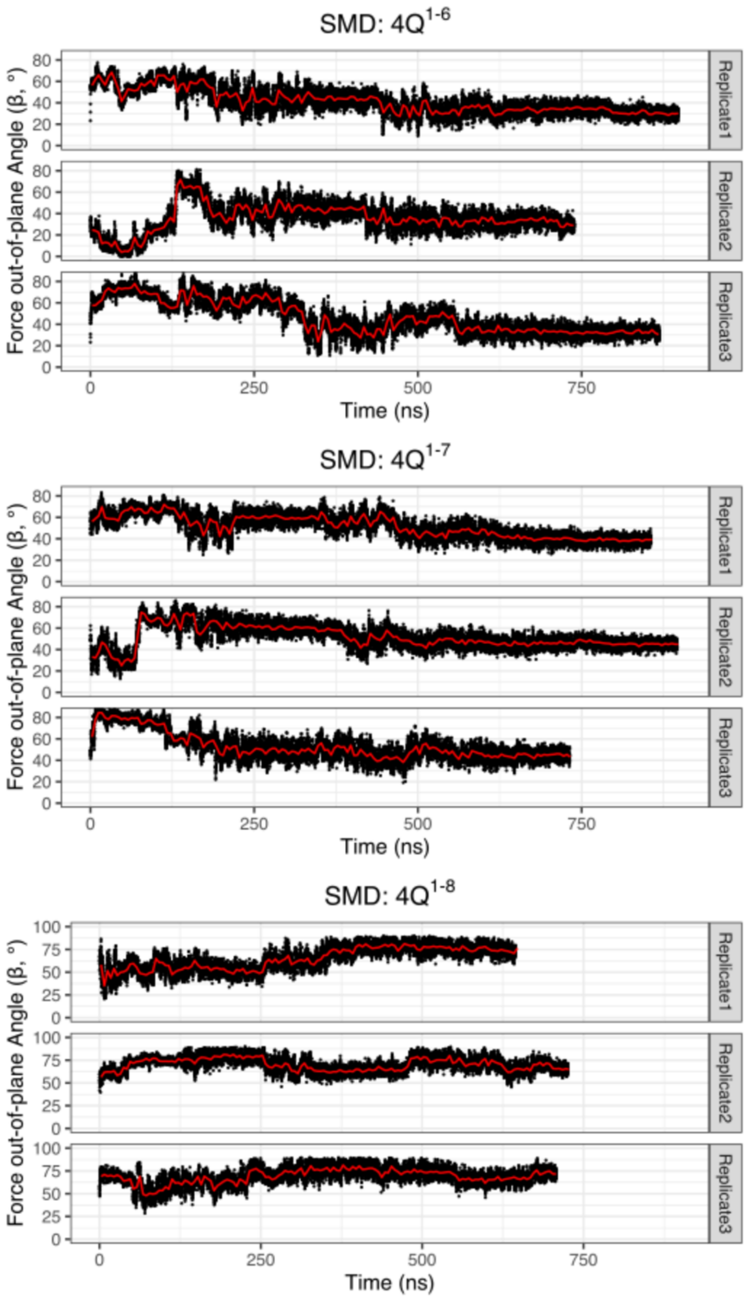


**Figure S6.** Evolution of force out-of-plane angle of 4Q-GQs during pulling, and the red solid lines show the time-averaged values.

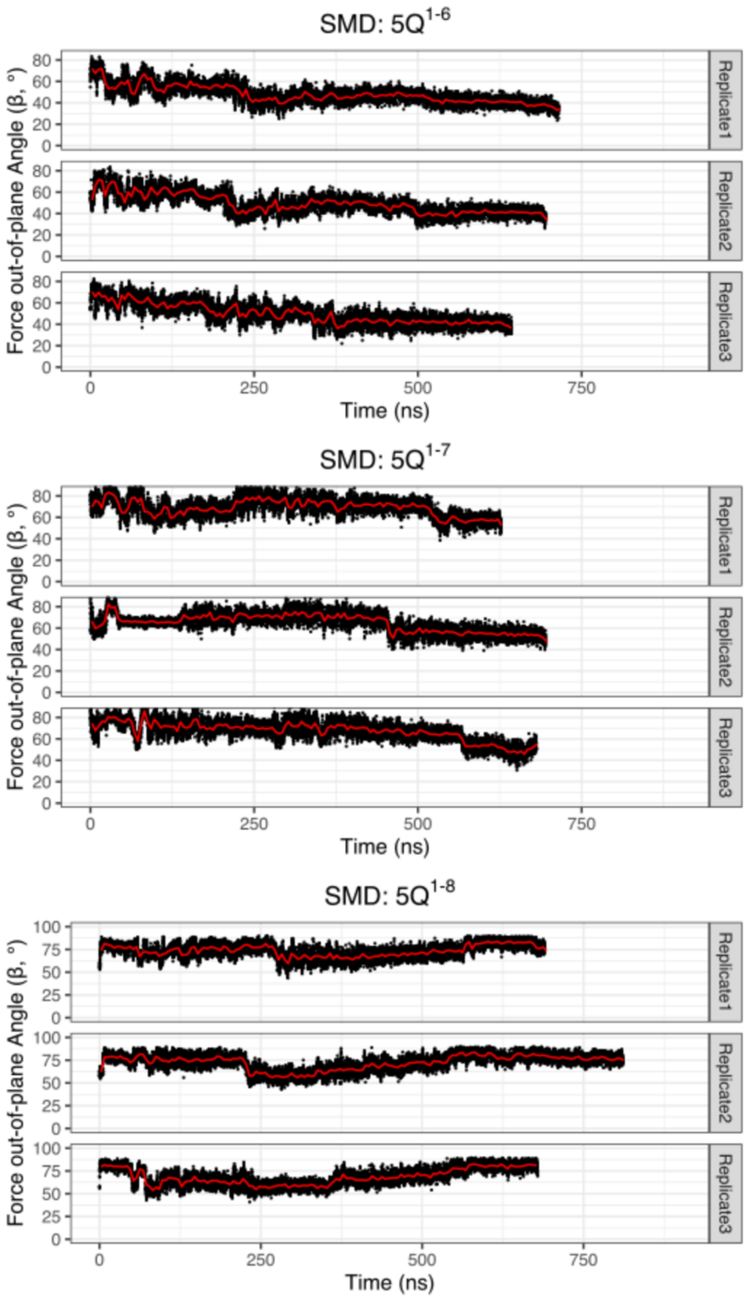


**Figure S7.** Evolution of force out-of-plane angle of 5Q-GQs during pulling, and the red solid lines show the time-averaged values.

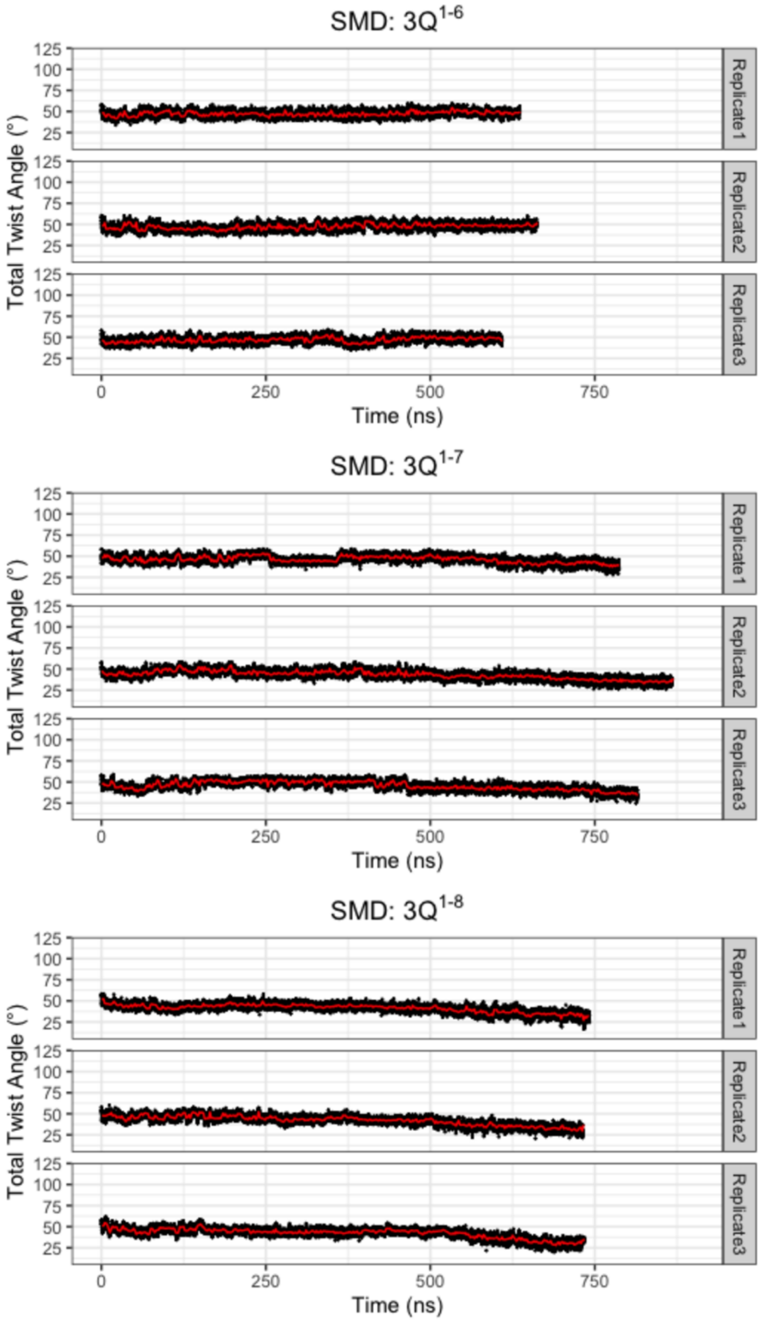


**Figure S8.** Evolution of total twist angles in SMD of 3Q-GQs, and the red solid lines show the time-averaged values.

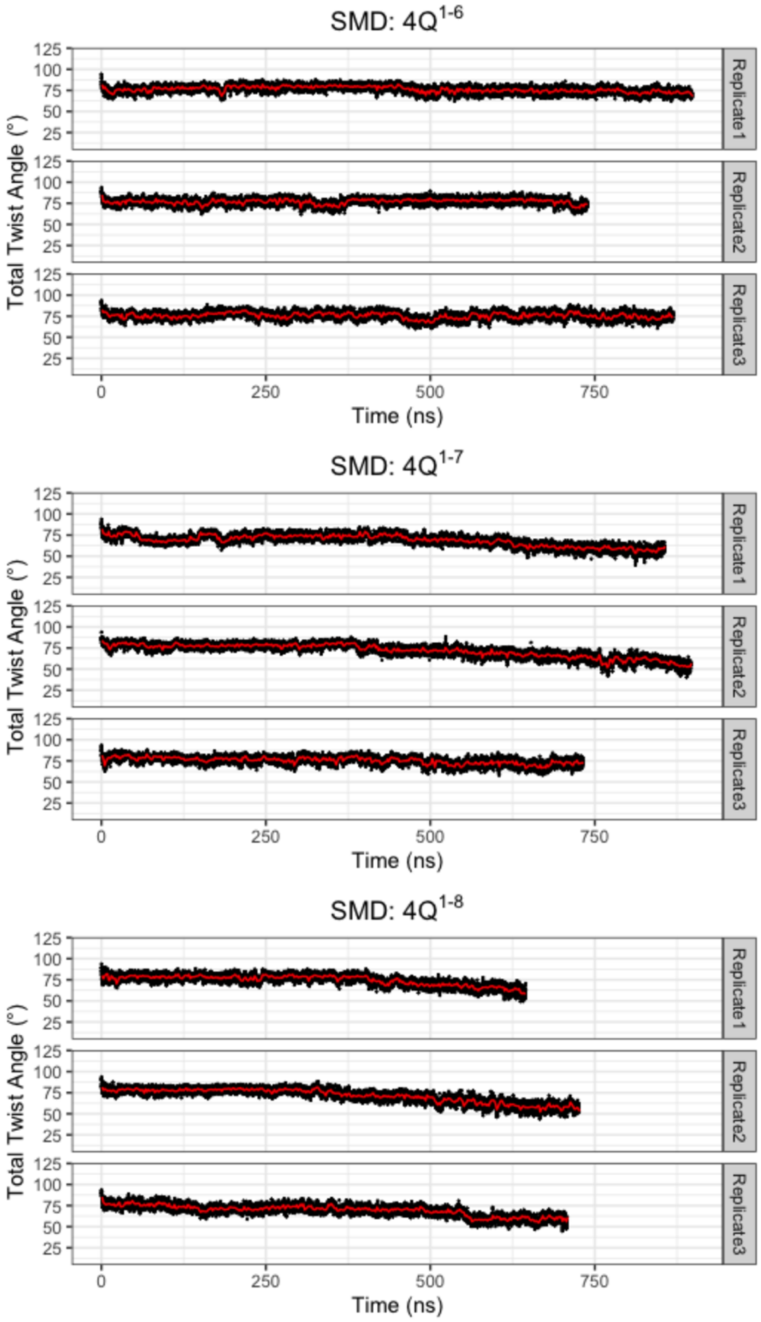


**Figure S9.** Evolution of total twist angles in SMD of 4Q-GQs, and the red solid lines show the time-averaged values.

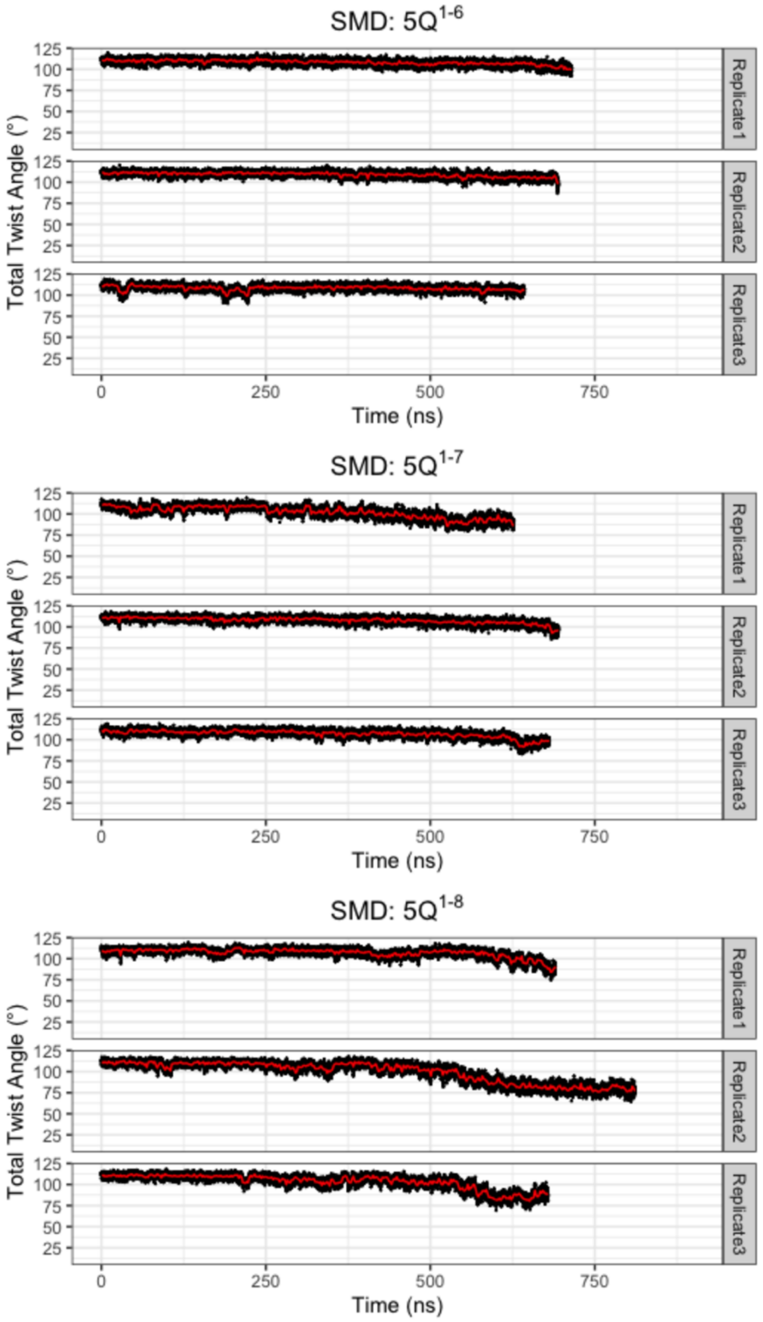


**Figure S10.** Evolution of total twist angles in SMD of 5Q-GQs, and the red solid lines show the time-averaged values.

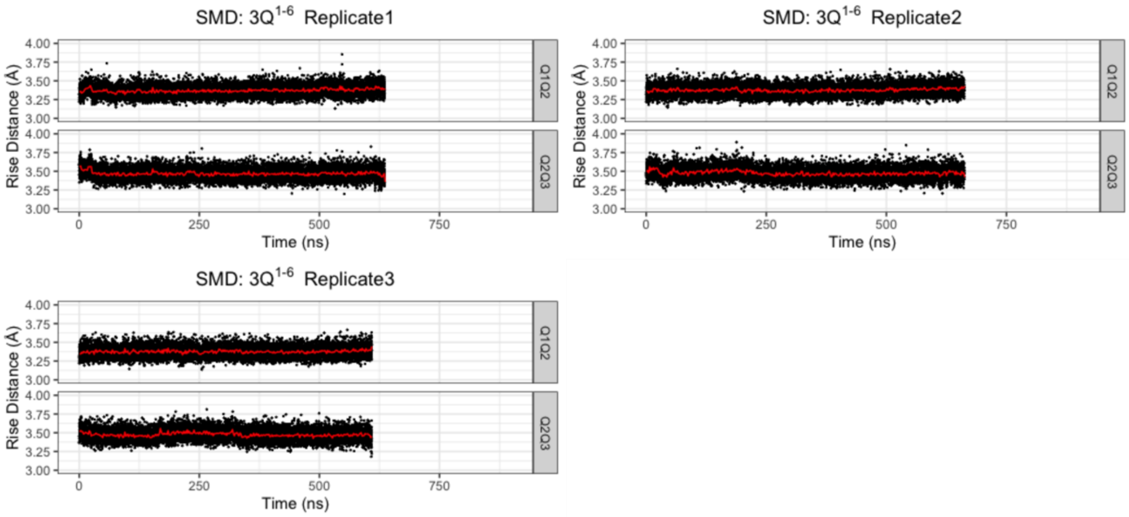


**Figure S11.** Evolution of rise distances in SMD of $3Q^{1-6}$, and the red solid lines show the time-averaged values.

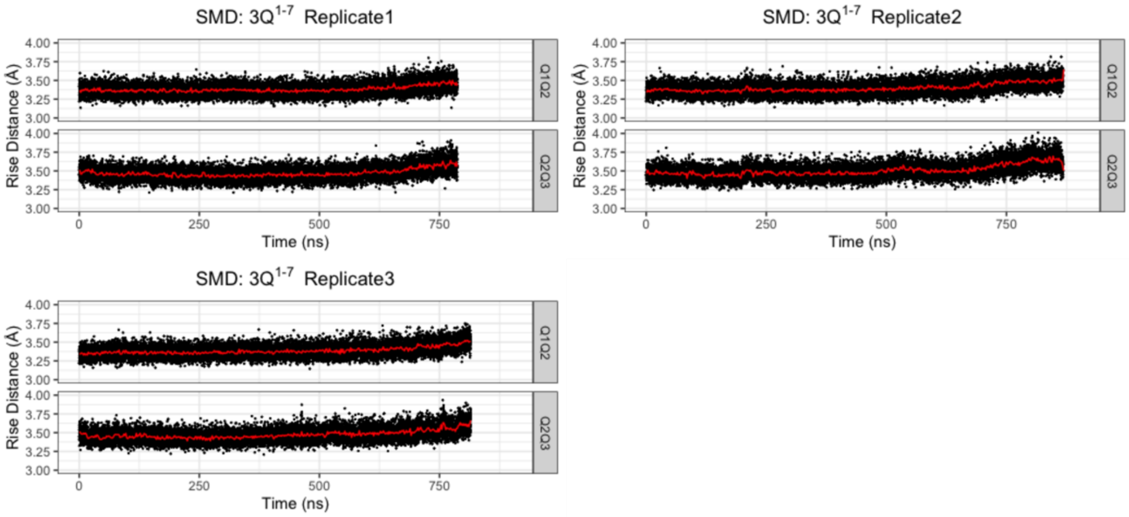


**Figure S12.** Evolution of rise distances in SMD of $3Q^{1-7}$, and the red solid lines show the time-averaged values.

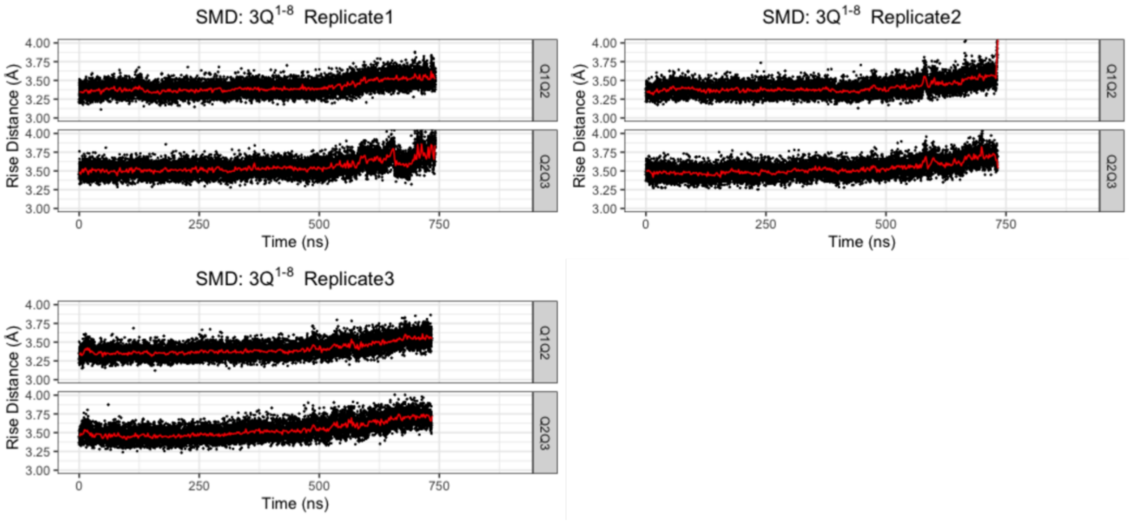

**Figure S13.** Evolution of rise distances in SMD of $3Q^{1-8}$, and the red solid lines show the time-averaged values.

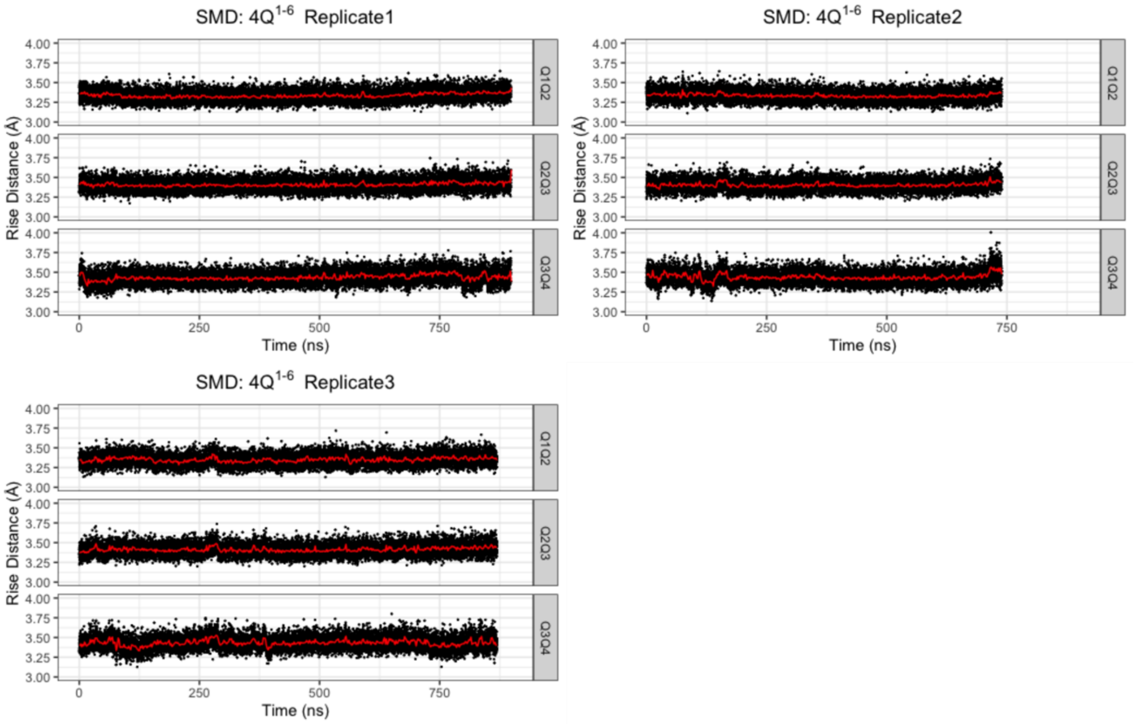


**Figure S14.** Evolution of rise distances in SMD of $4Q^{1-6}$, and the red solid lines show the time-averaged values.

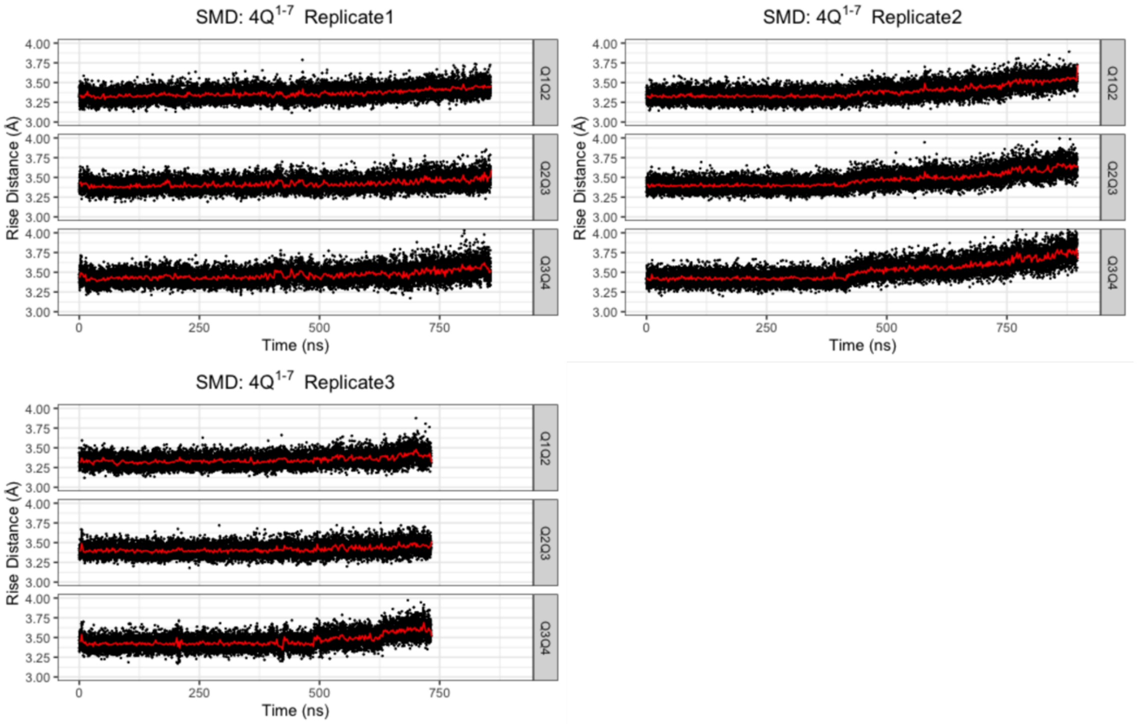


**Figure S15.** Evolution of rise distances in SMD of $4Q^{1-7}$, and the red solid lines show the time-averaged values.

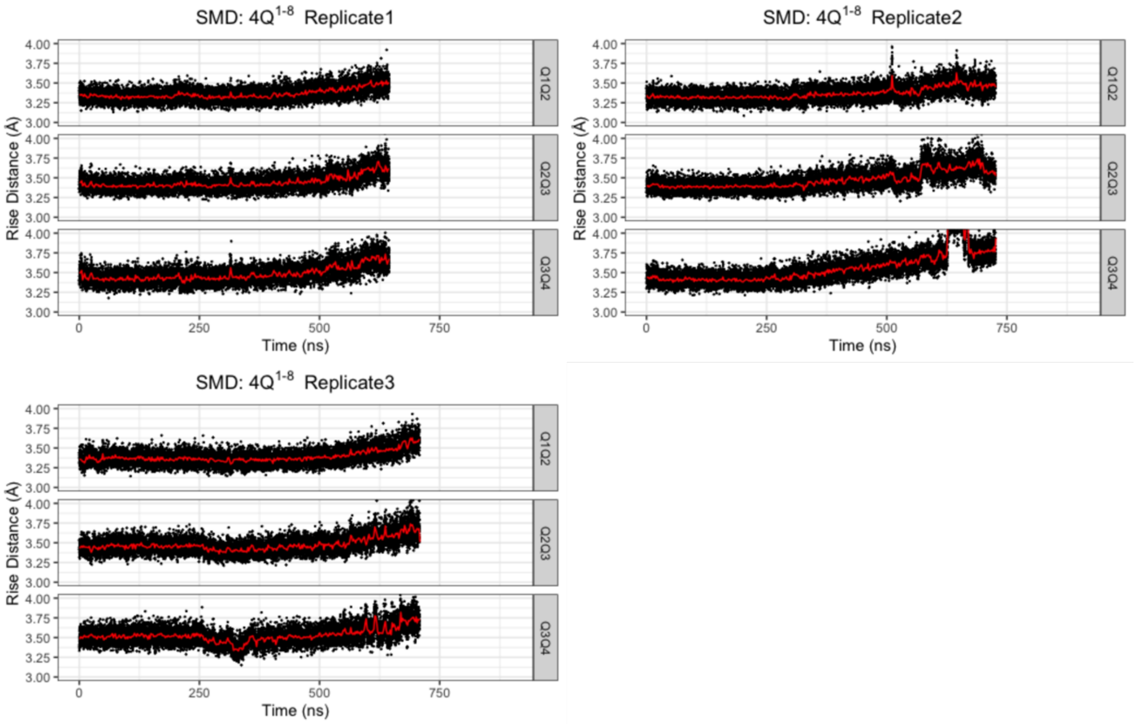


**Figure S16.** Evolution of rise distances in SMD of $4Q^{1\text{-}8}$, and the red solid lines show the time-averaged values.

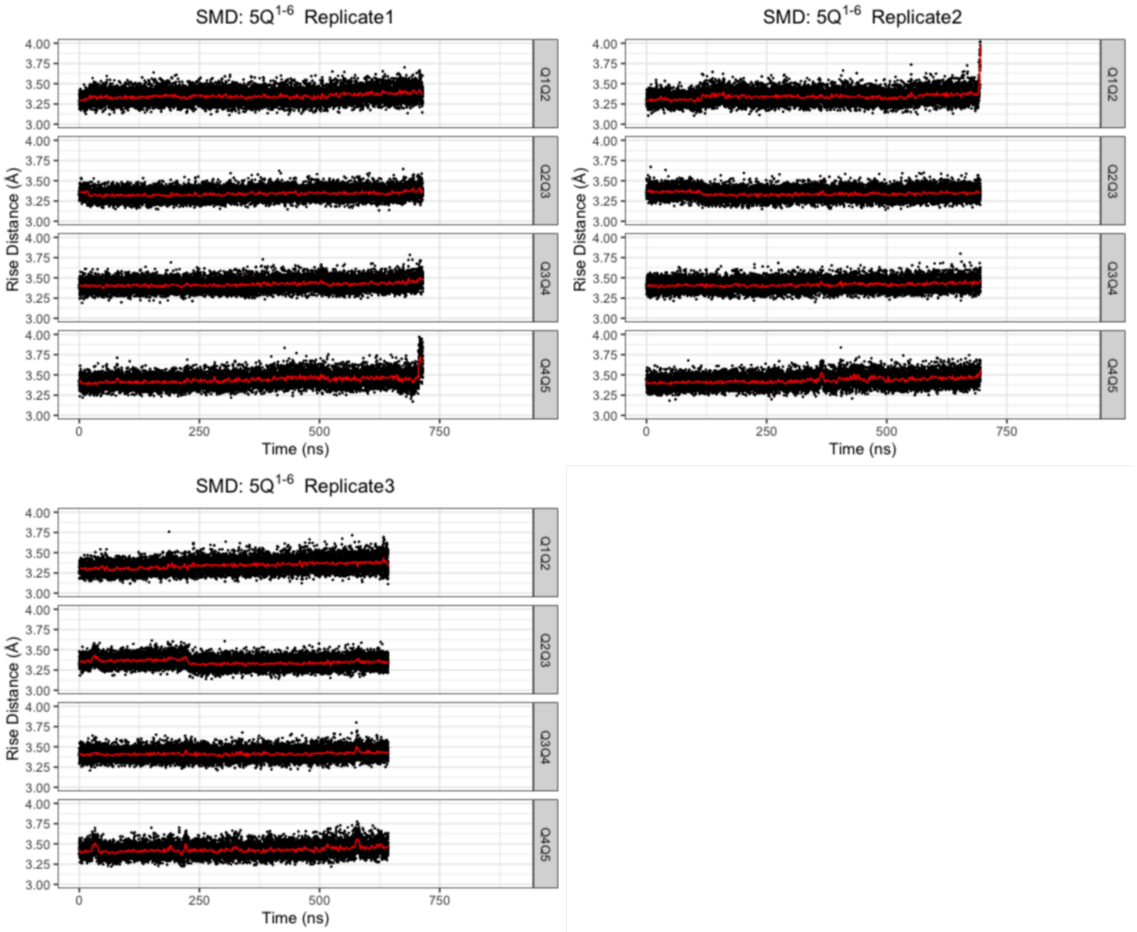


**Figure S17.** Evolution of rise distances in SMD of $5Q^{1\text{-}6}$, and the red solid lines show the time-averaged values.

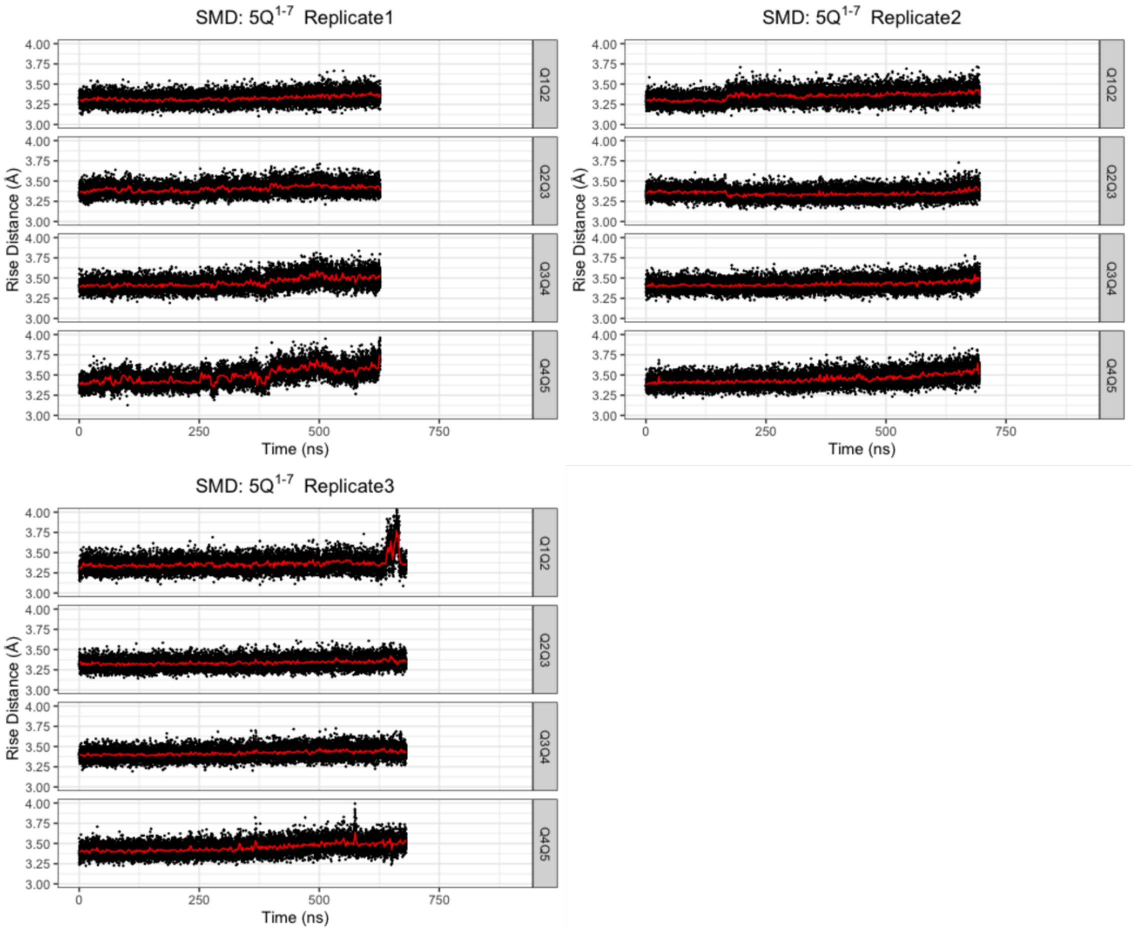


**Figure S18.** Evolution of rise distances in SMD of 5Q$^{1\text{-}7}$, and the red solid lines show the time-averaged values.

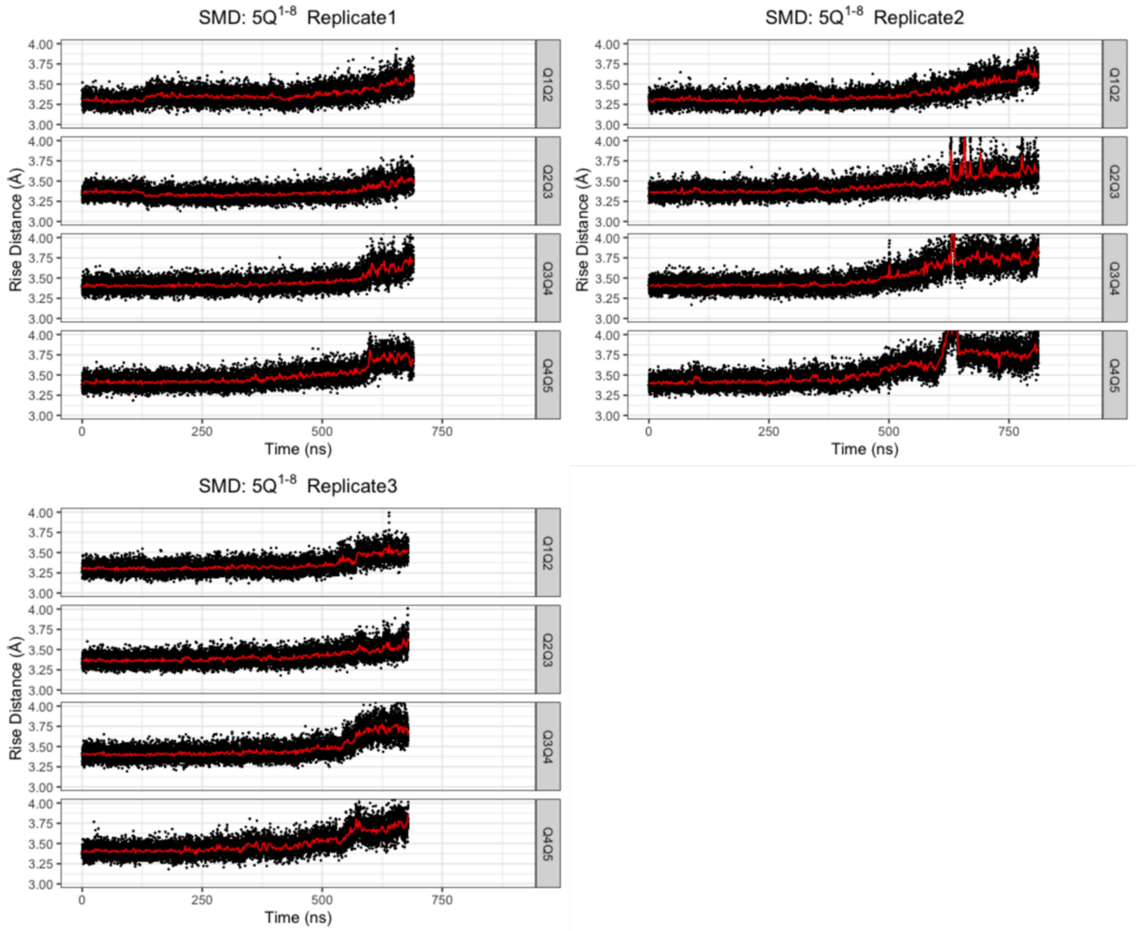


**Figure S19.** Evolution of rise distances in SMD of 5Q$^{1-8}$, and the red solid lines show the time-averaged values.

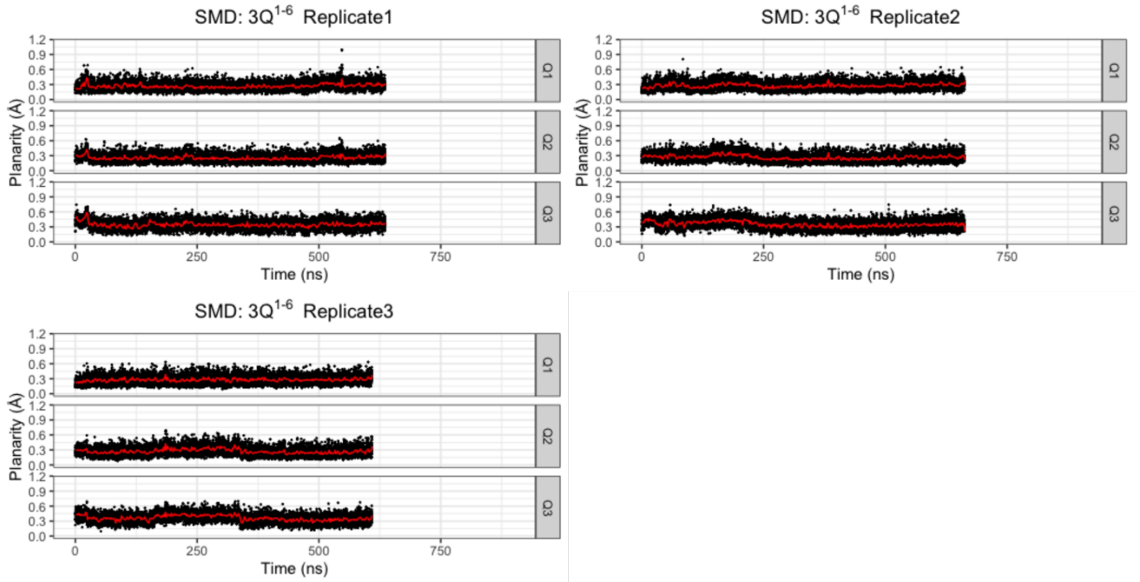


**Figure S20.** Evolution of planarity in SMD of 3Q$^{1-6}$, and the red solid lines show the time-averaged values.

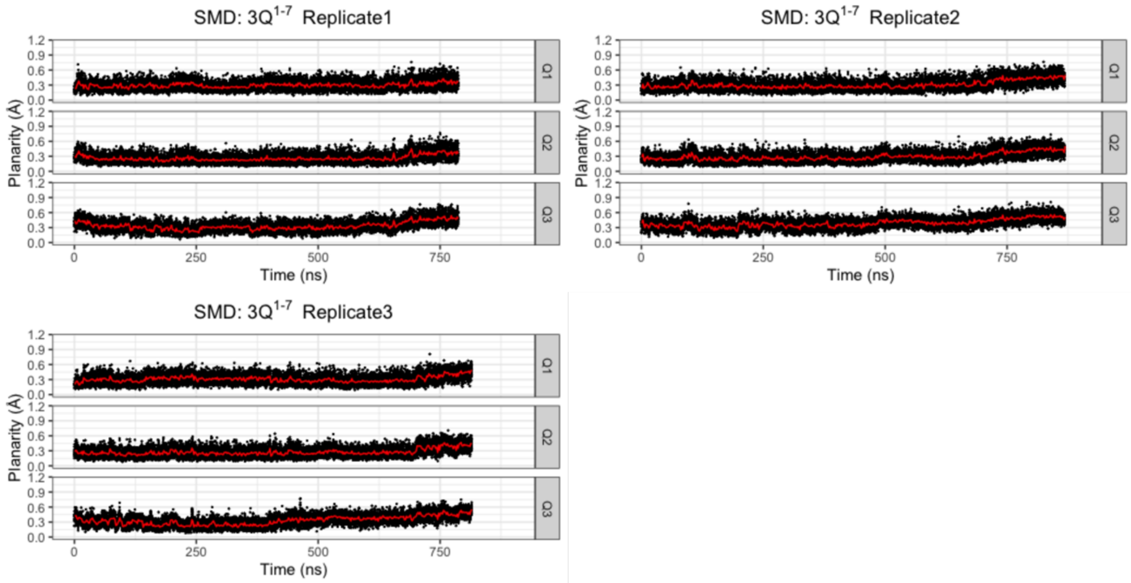


**Figure S21.** Evolution of planarity in SMD of 3Q[1-7], and the red solid lines show the time-averaged values.

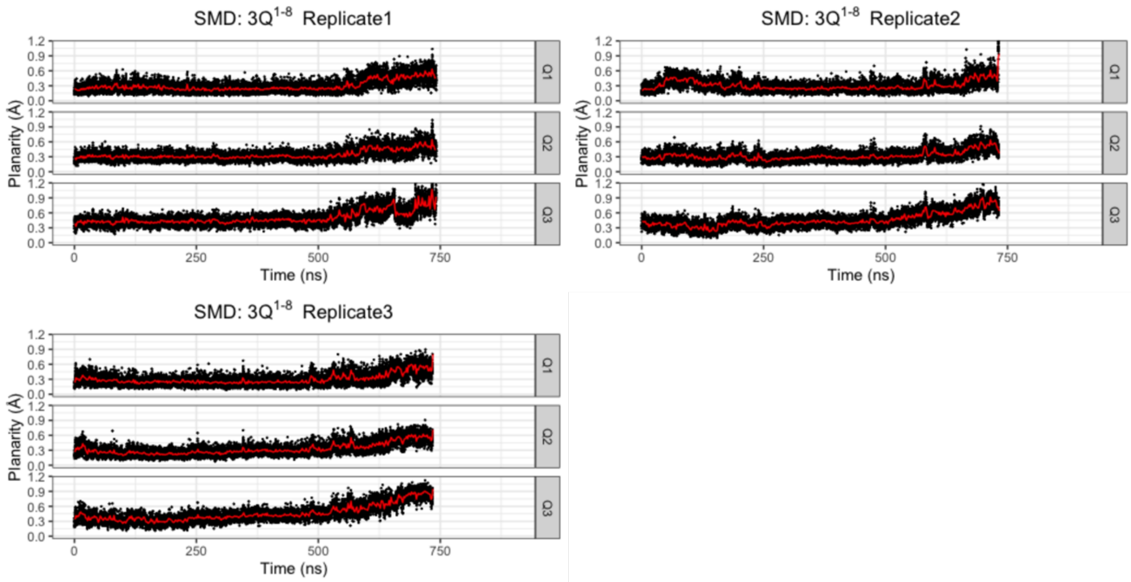


**Figure S22.** Evolution of planarity in SMD of 3Q[1-8], and the red solid lines show the time-averaged values.

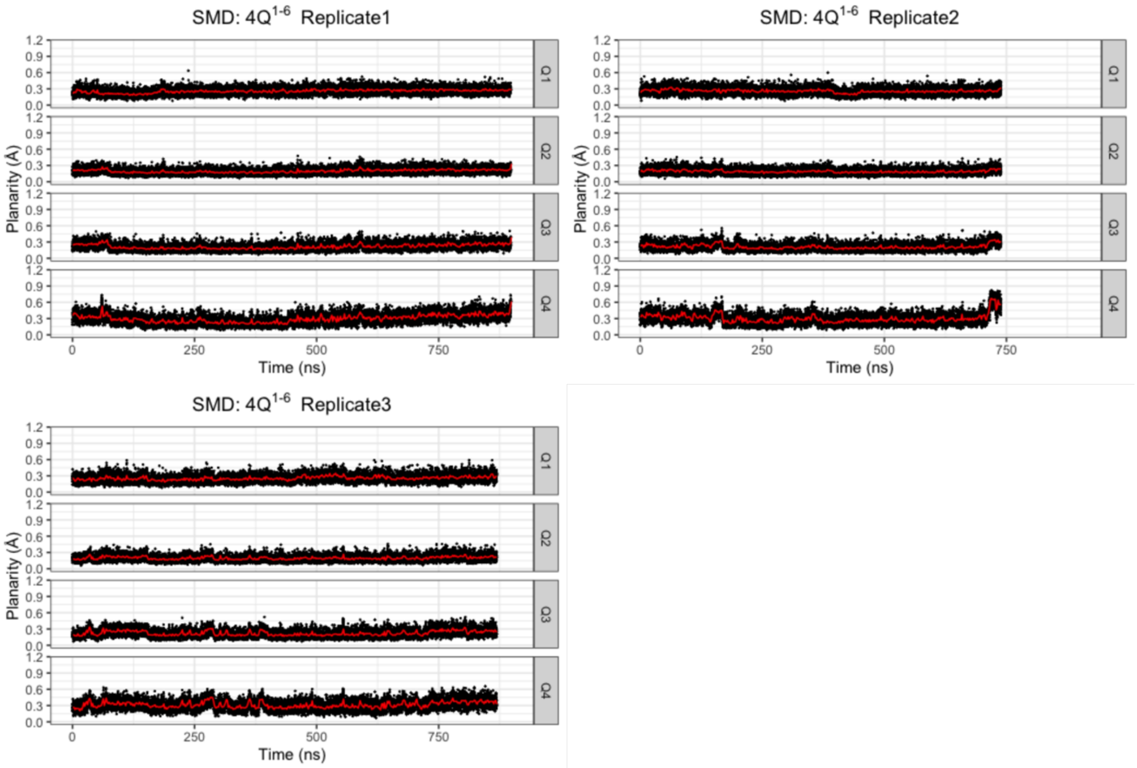


**Figure S23.** Evolution of planarity in SMD of $4Q^{1-6}$, and the red solid lines show the time-averaged values.

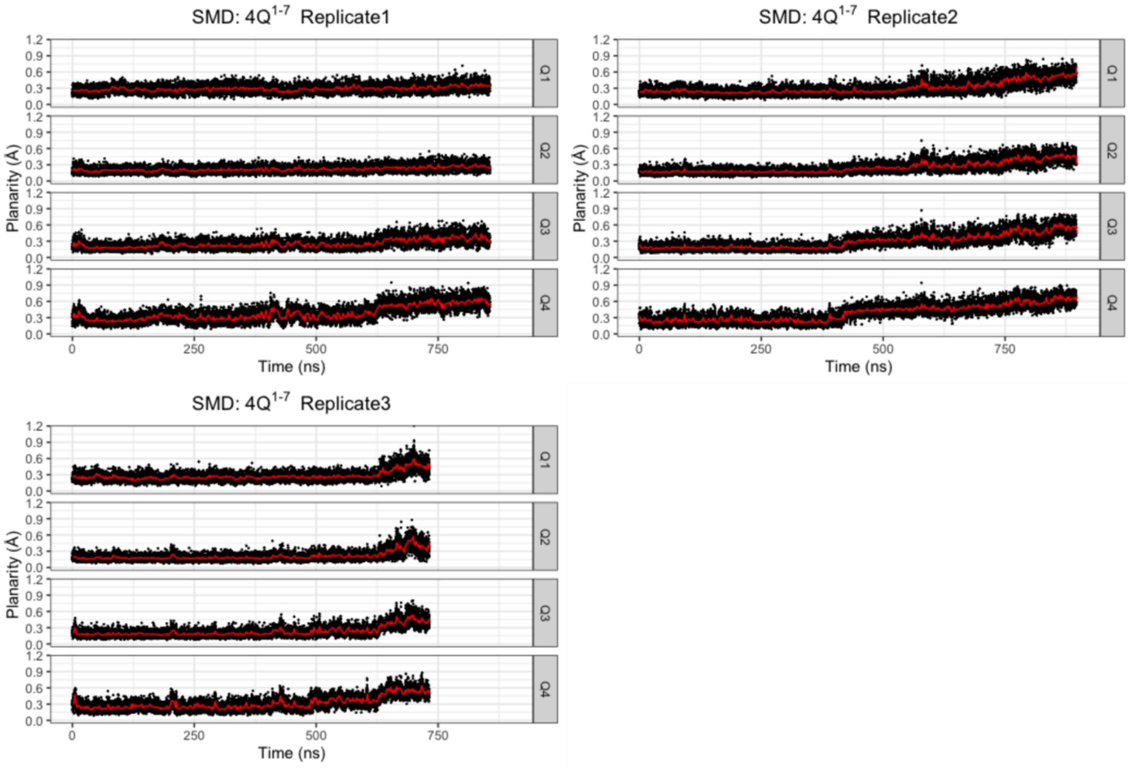


**Figure S24.** Evolution of planarity in SMD of $4Q^{1-7}$, and the red solid lines show the time-averaged values.

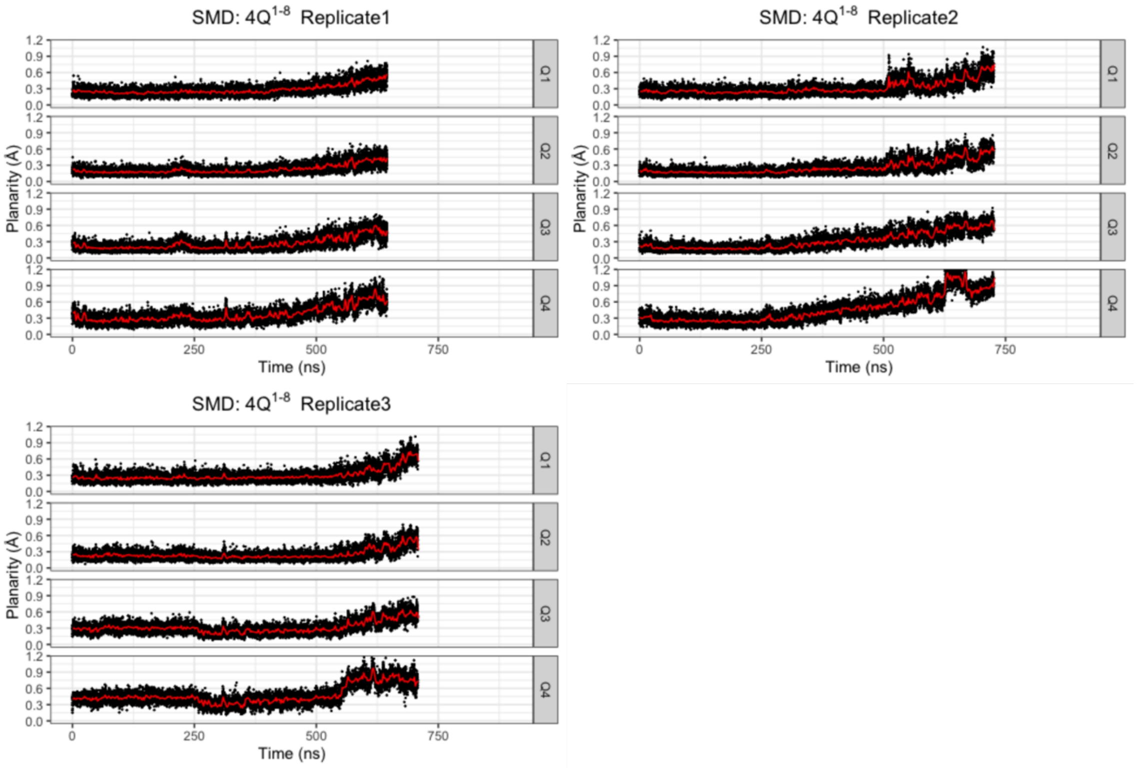


**Figure S25.** Evolution of planarity in SMD of $4Q^{1\text{-}8}$, and the red solid lines show the time-averaged values.

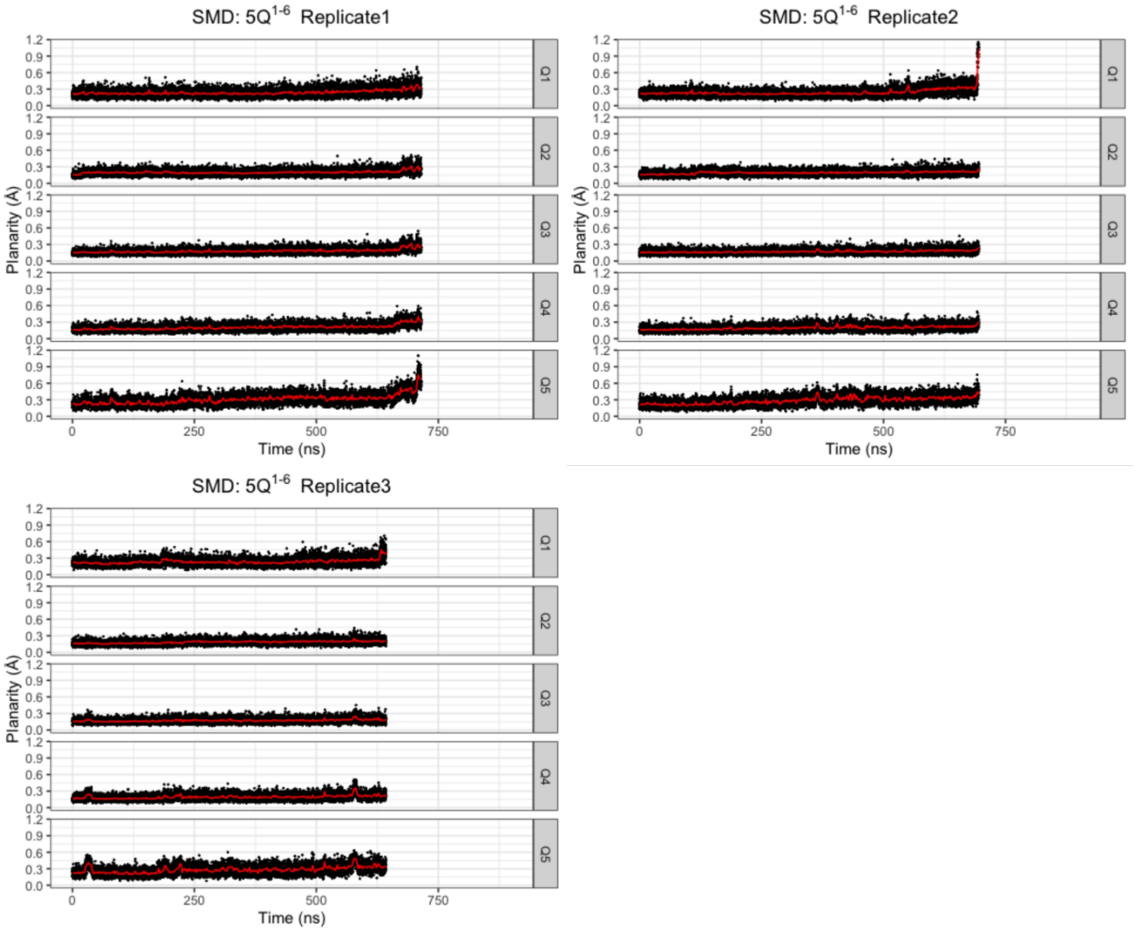

**Figure S26.** Evolution of planarity in SMD of $5Q^{1-6}$, and the red solid lines show the time-averaged values.

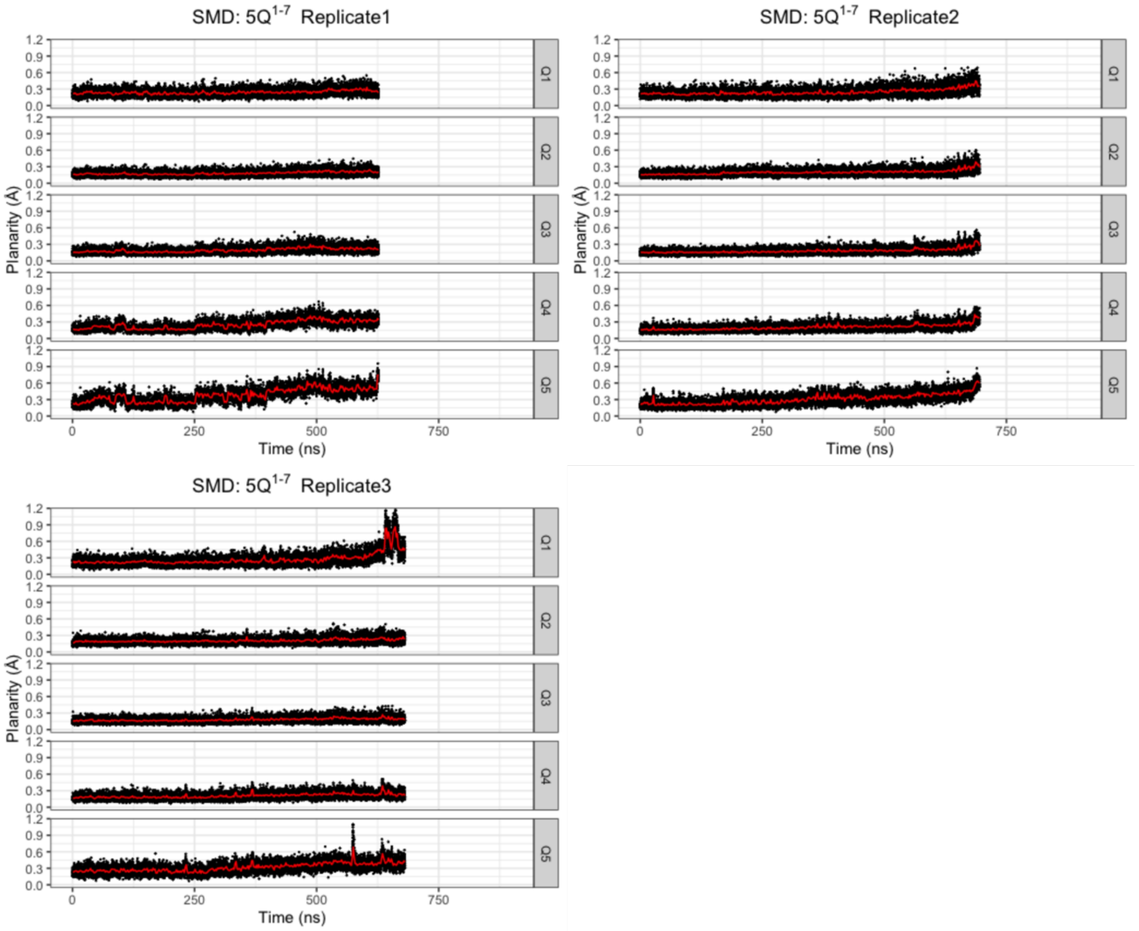


**Figure S27.** Evolution of planarity in SMD of $5Q^{1-7}$, and the red solid lines show the time-averaged values.

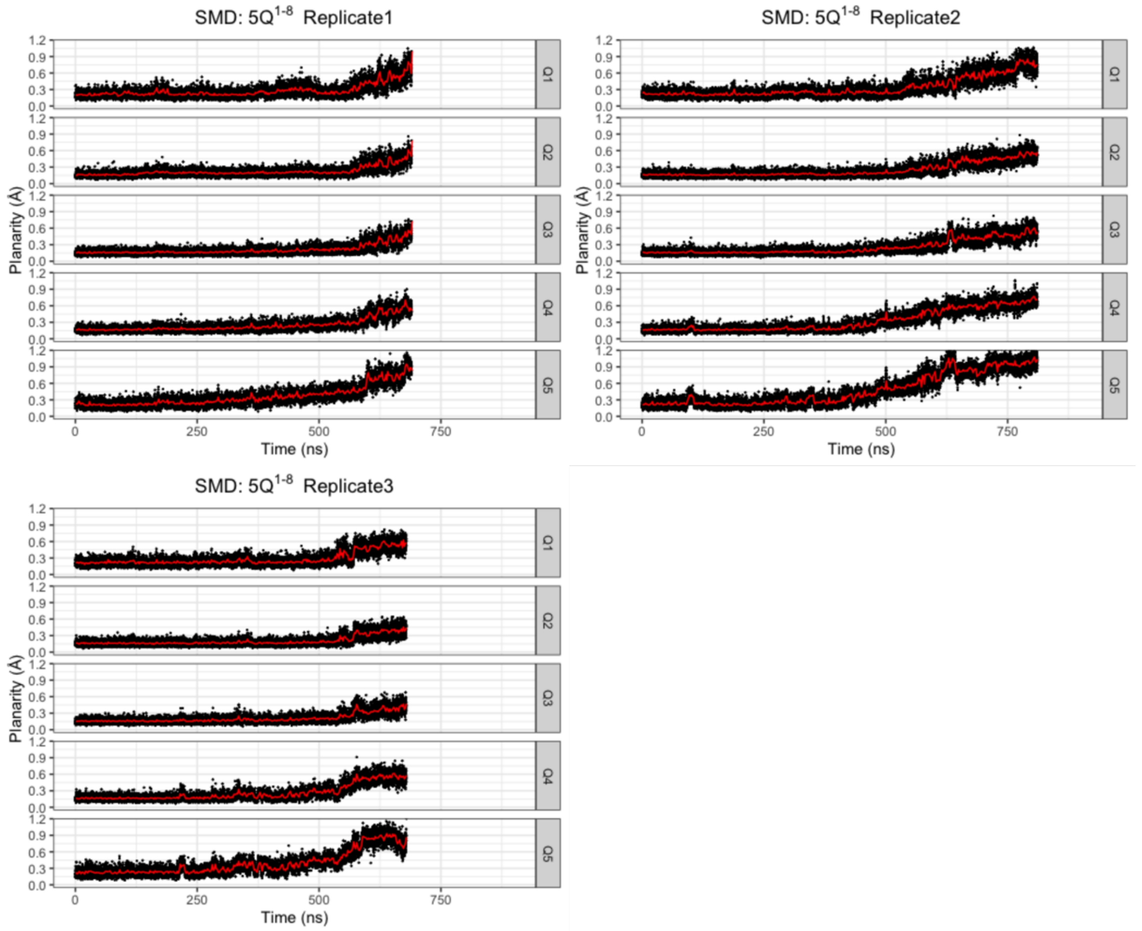


**Figure S28.** Evolution of planarity in SMD of $5Q^{1-8}$, and the red solid lines show the time-averaged values.

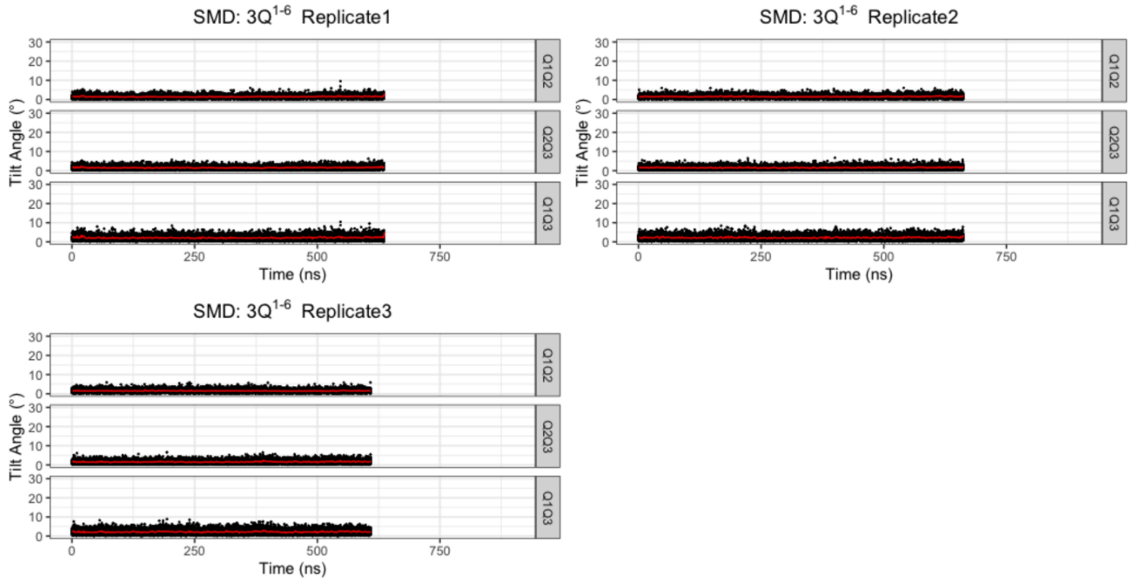


**Figure S29.** Evolution of tilt angles in SMD of $3Q^{1-6}$, and the red solid lines show the time-averaged values.

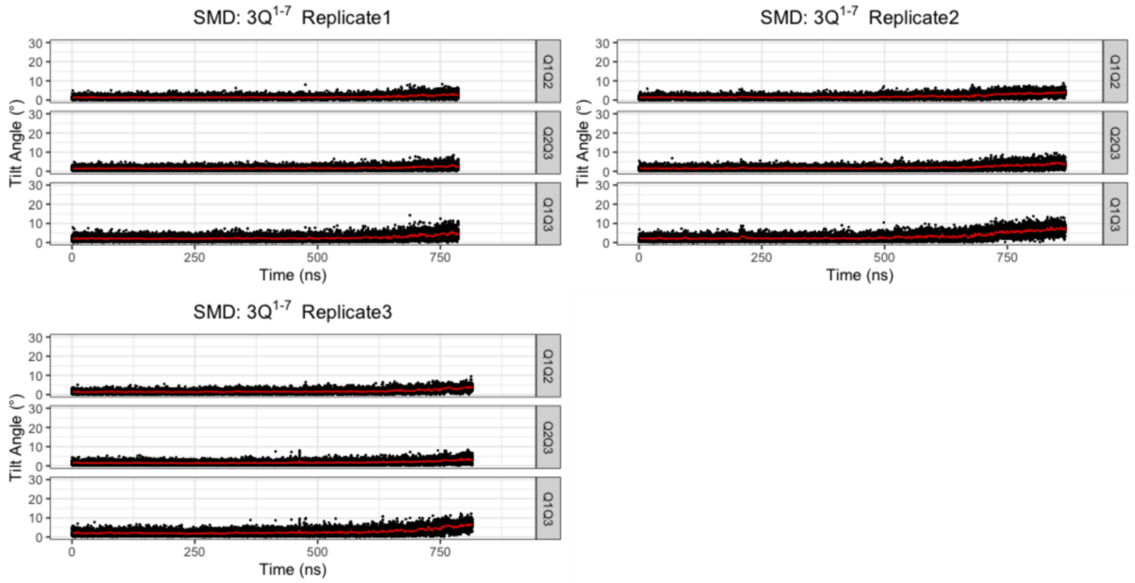


**Figure S30.** Evolution of tilt angles in SMD of $3Q^{1\text{-}7}$, and the red solid lines show the time-averaged values.

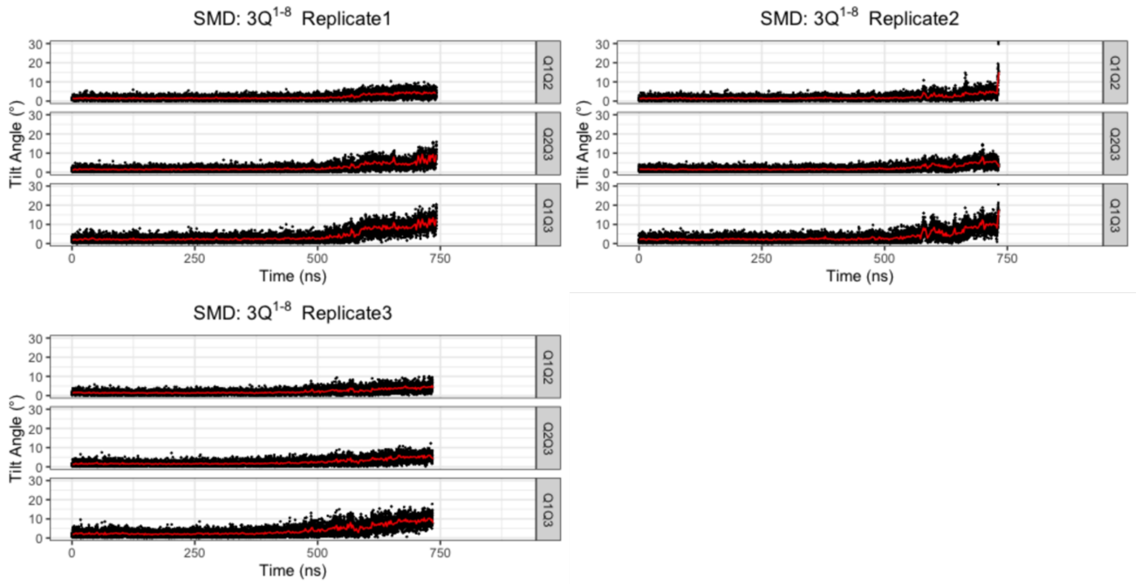


**Figure S31.** Evolution of tilt angles in SMD of $3Q^{1\text{-}8}$, and the red solid lines show the time-averaged values.

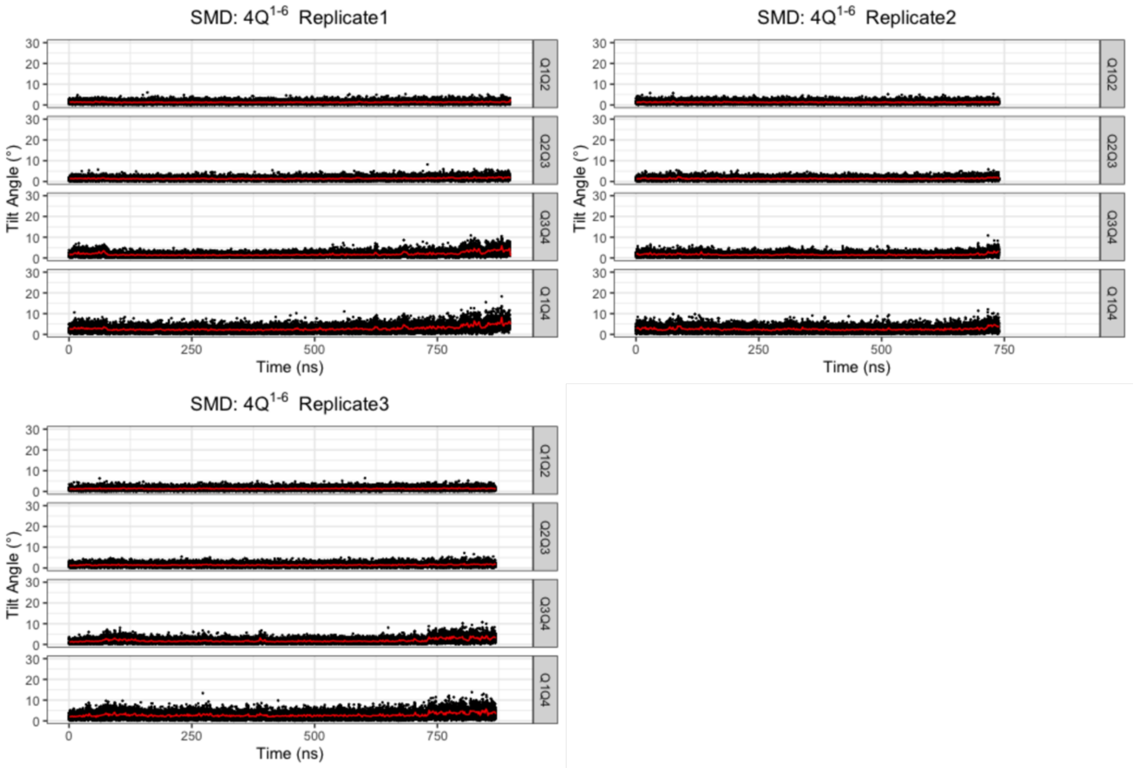


**Figure S32.** Evolution of tilt angles in SMD of 4Q$^{1\text{-}6}$, and the red solid lines show the time-averaged values.

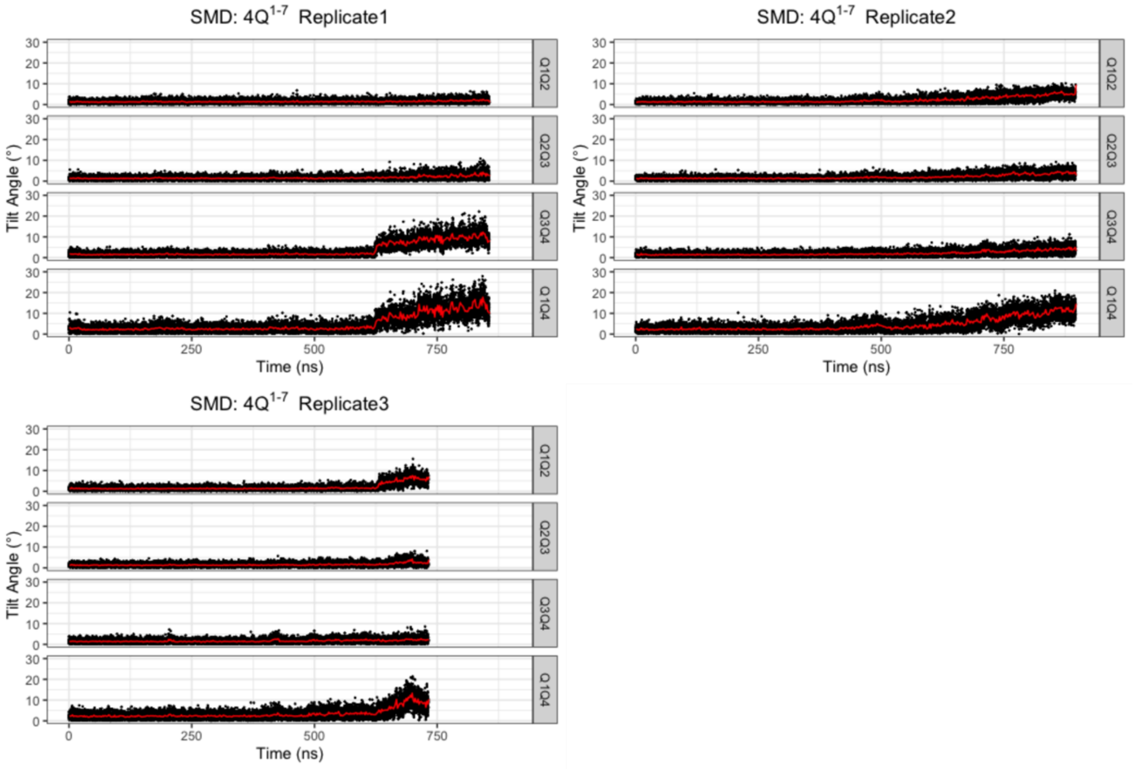


**Figure S33.** Evolution of tilt angles in SMD of 4Q$^{1\text{-}7}$, and the red solid lines show the time-averaged values.

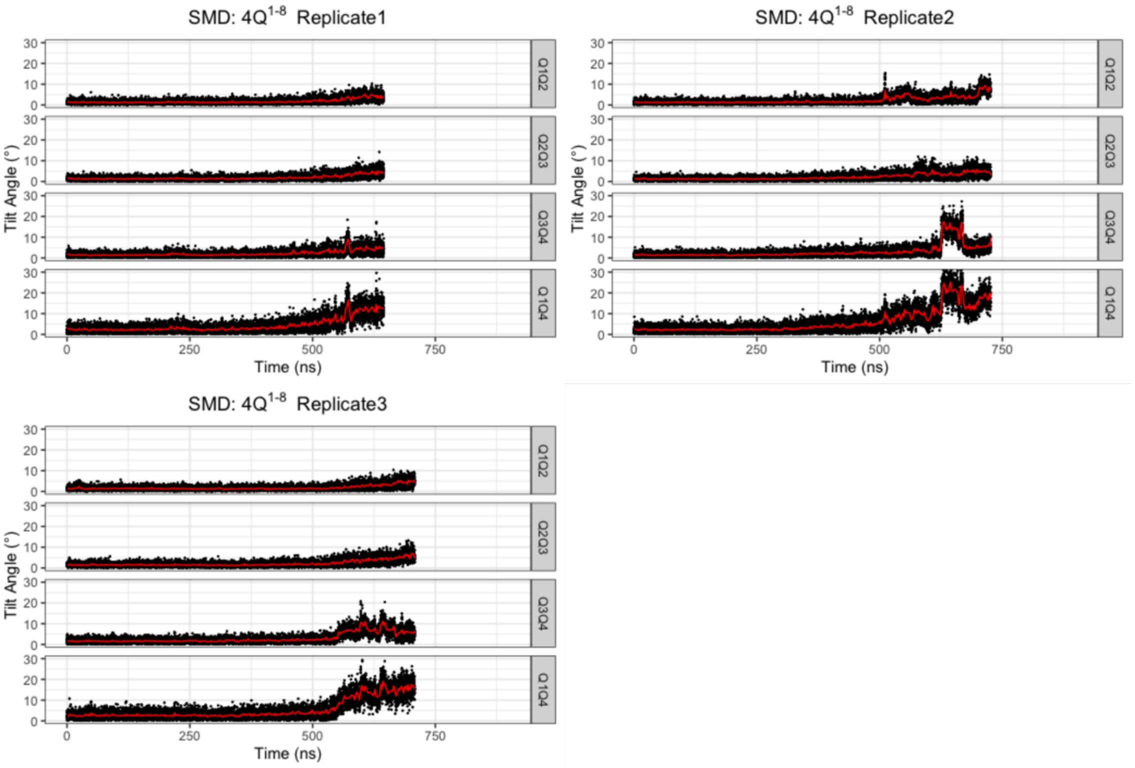


**Figure S34.** Evolution of tilt angles in SMD of $4Q^{1\text{-}8}$, and the red solid lines show the time-averaged values.

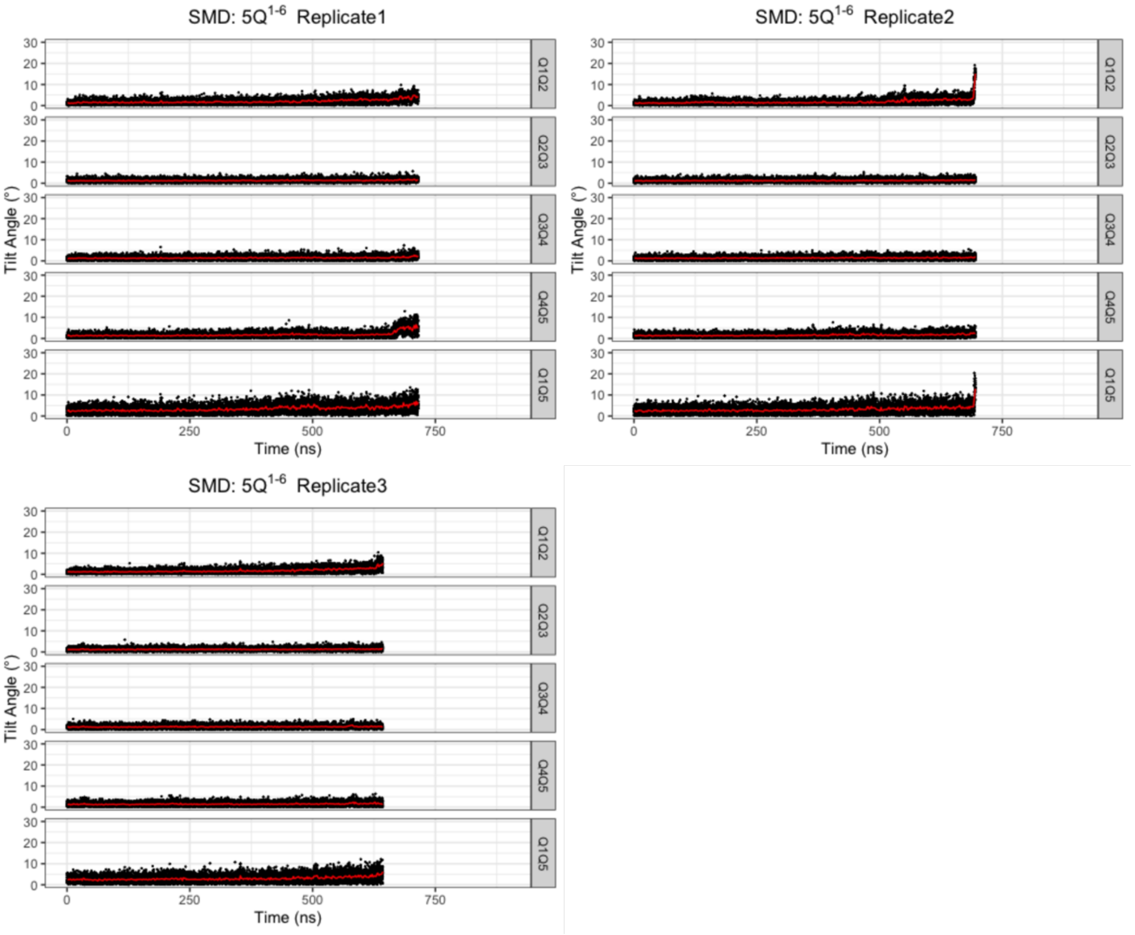

**Figure S35.** Evolution of tilt angles in SMD of 5Q$^{1\text{-}6}$, and the red solid lines show the time-averaged values.

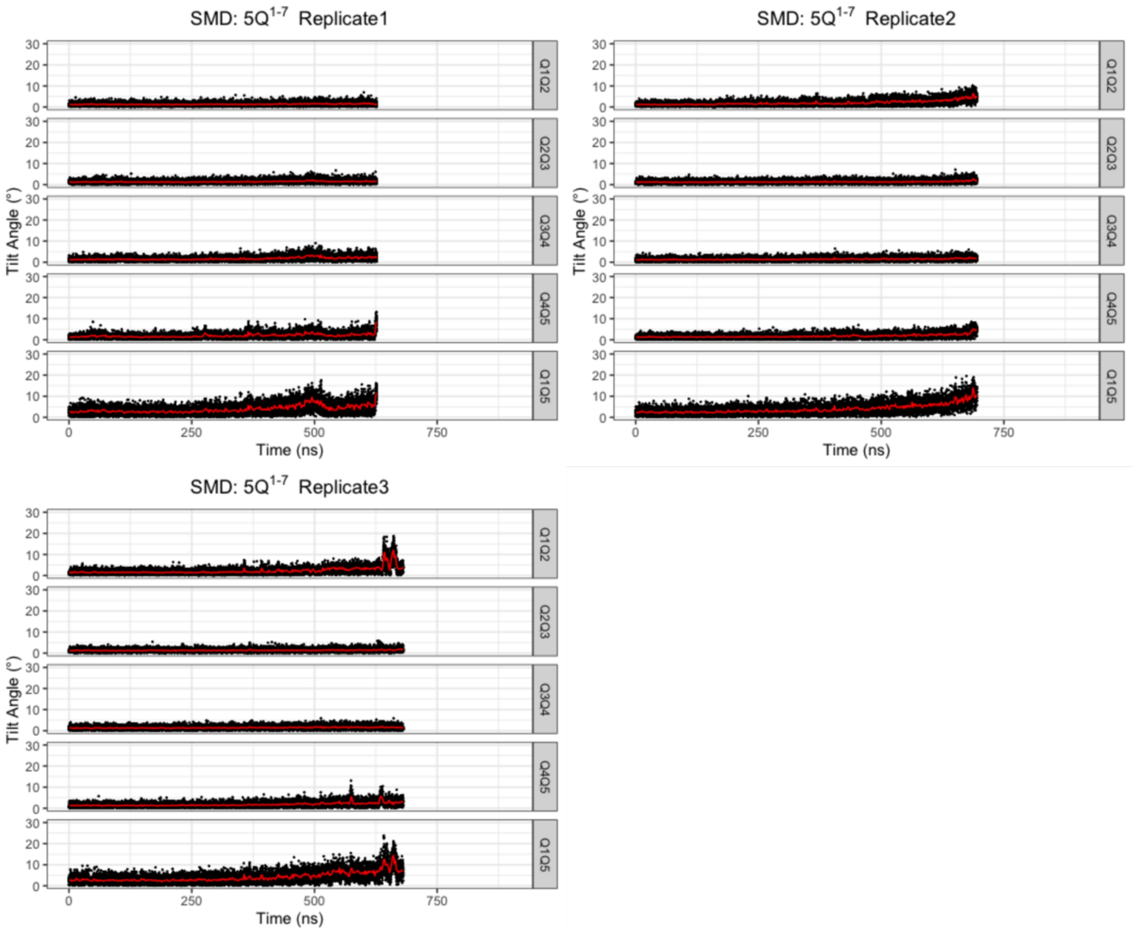


**Figure S36.** Evolution of tilt angles in SMD of 5Q$^{1\text{-}7}$, and the red solid lines show the time-averaged values.

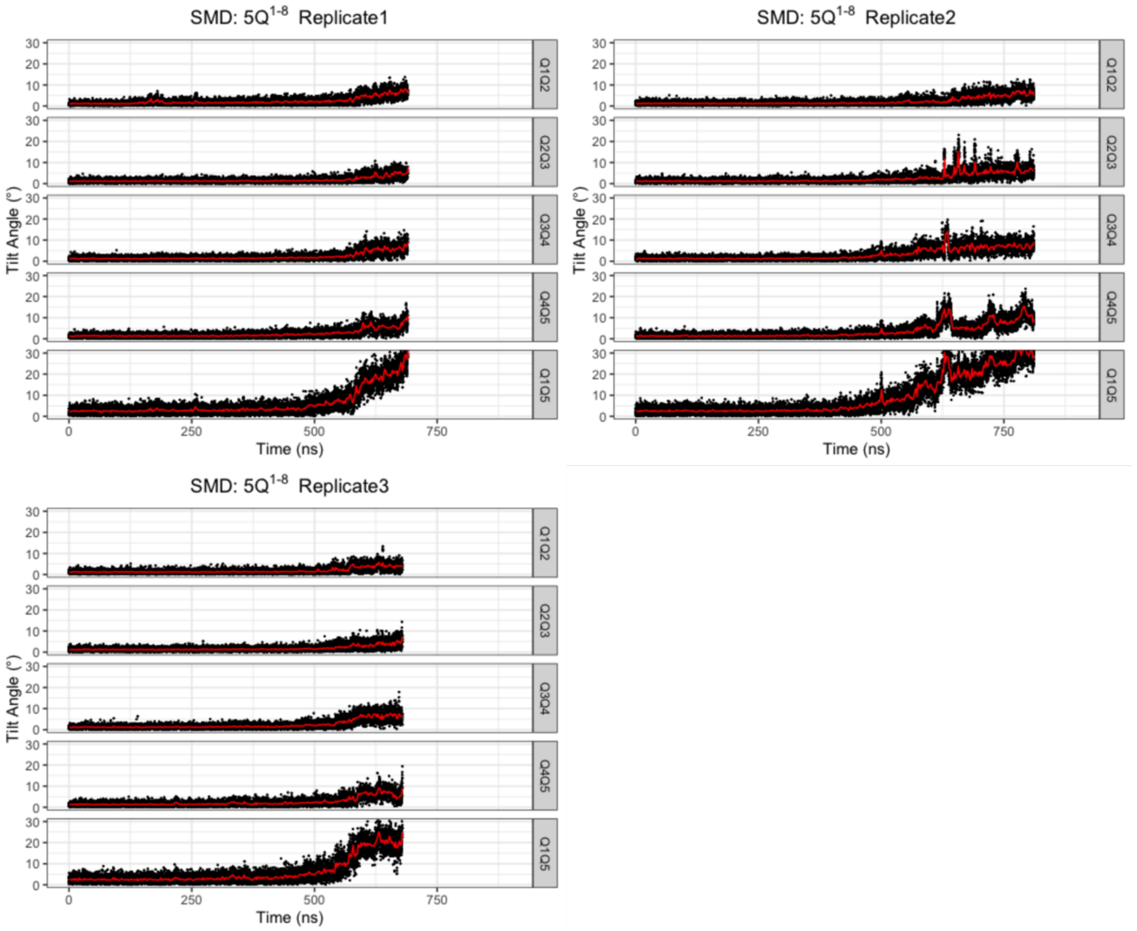


**Figure S37.** Evolution of tilt angles in SMD of $5Q^{1\text{-}8}$, and the red solid lines show the time-averaged values.

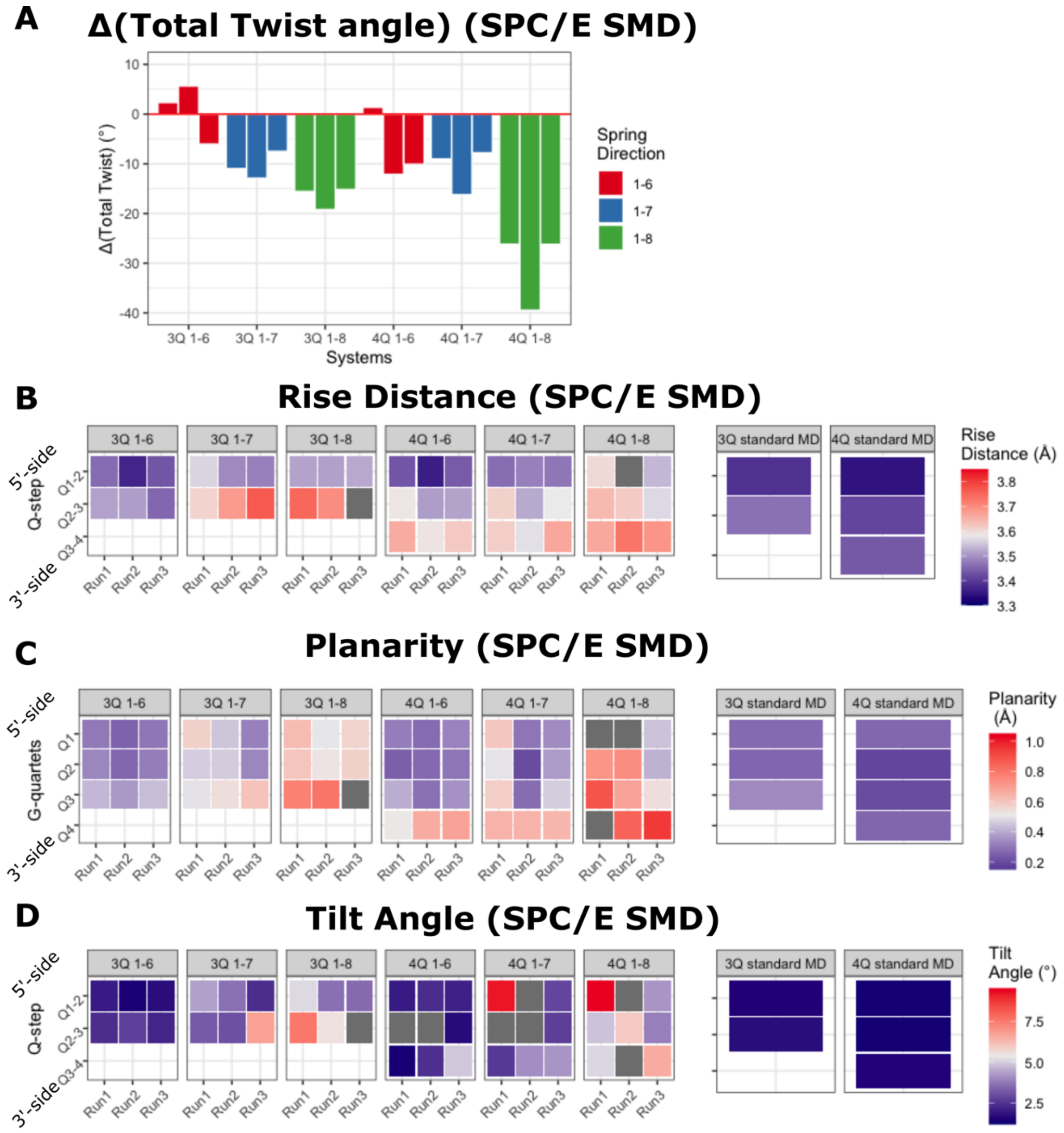


**Figure S38. GQ conformational features right before unfolding occurred (SMD in SPC/E water). (A)** The changes of total twist angles in all SMD simulations from the equilibrium GQ structure to the GQ ahead of the first unfolding event. The rise distances **(B)**, planarity **(C)**, and tilt angles **(D)** for all quartet-quartet steps in SMD simulations, with the corresponding reference values from standard MD simulations on the right side. See Tables S11-S13 for exact values.

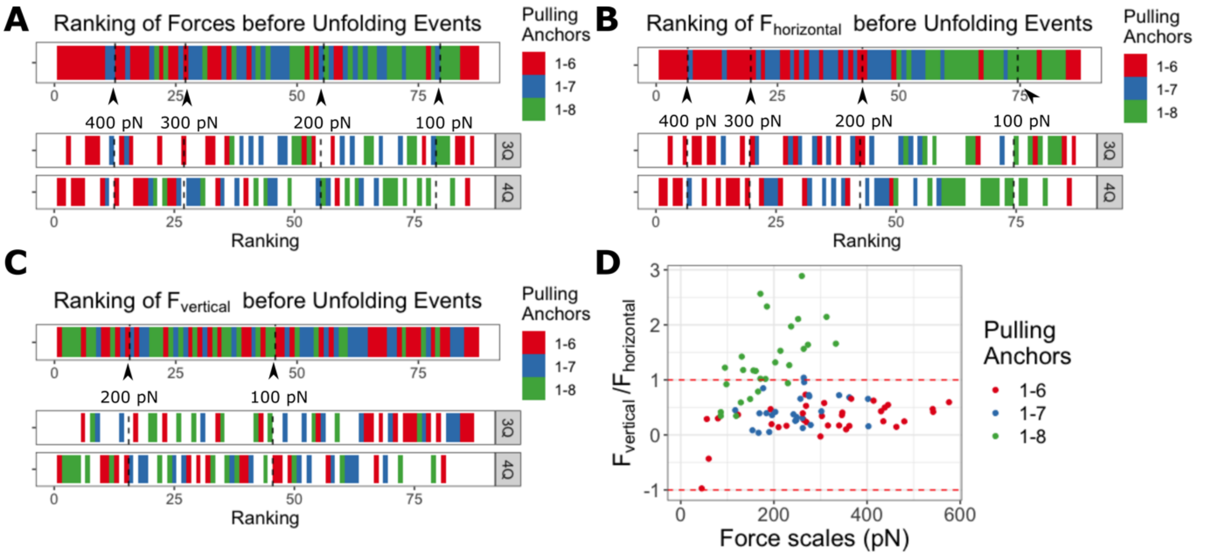


**Figure S39. Comparison of transition-inducing forces in SPC/E water. (A)** The ranking (highest to smallest) of all force magnitudes. **(B)** The ranking of the horizontal components of all transition-inducing forces. **(C)** The ranking of the vertical components of all transition-inducing forces. **(D)** The force distribution in the dimension of force magnitude and the vertical-to-horizontal force components ratio. The region between the red dashed lines has the horizontal force component higher than the vertical one. Note that the data with negative ratio have the relative positions of 5'/3'-anchors upside down with respect to the reference GQ plane.

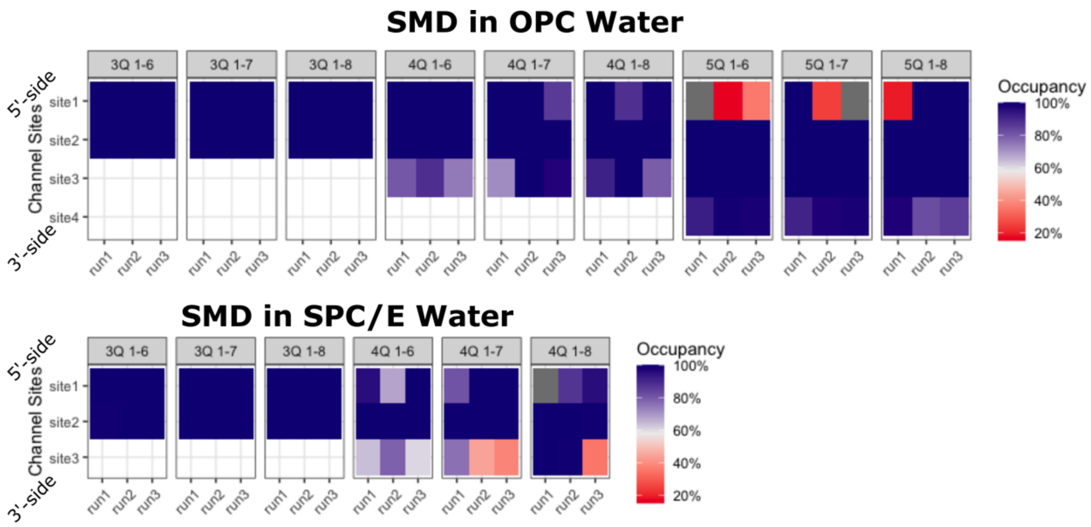


**Figure S40. Occupation rates of the GQ channel $K^+$ binding sites in SMD simulations with OPC and SPC/E water models.** The occupancies were averaged through the SMD simulation time before the starting of the unfolding.

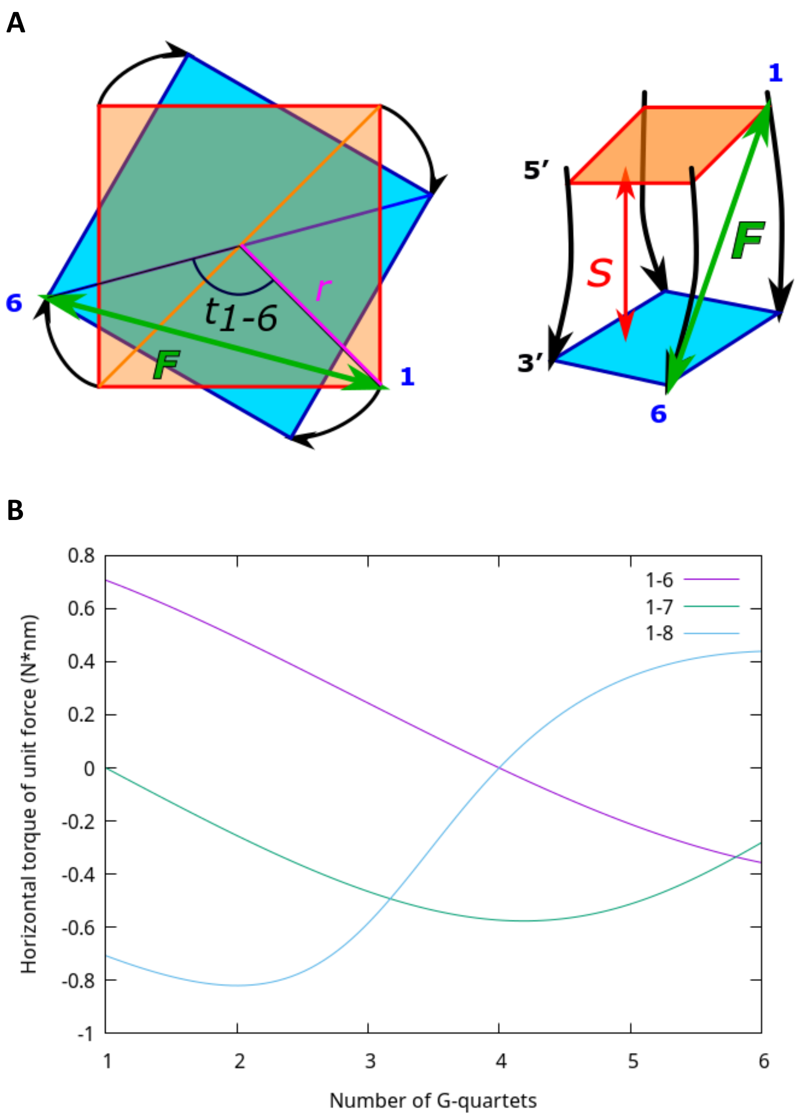


**Figure S41.** (**A**) Graphical representation of the arguments used in the calculation of horizontal torque in the case of 1-6 pulling (see the Mathematical model section in Supporting Information); the left and right panel depicts top and side view of GQ, respectively. (**B**) Horizontal torque caused by a unit force on a native GQ structure. The curves show dependence of the toque in horizontal direction on the number of G-quartets present in the native structure between the indicated anchor points. Positive torque induces GQ helix overwinding, whereas negative torque unwinding. Note that the horizontal torque inducing overwinding can be overwhelmed by the unwinding effect of the vertical force component, such as in the case of $5Q^{1\text{-}8}$.
